# ORBIT for *E. coli*: Kilobase-scale oligonucleotide recombineering at high throughput and high efficiency

**DOI:** 10.1101/2023.06.28.546561

**Authors:** Scott H. Saunders, Ayesha M. Ahmed

## Abstract

Microbiology and synthetic biology depend on reverse genetic approaches to manipulate bacterial genomes; however, existing methods require molecular biology to generate genomic homology, suffer from low efficiency, and are not easily scaled to high throughput applications. To overcome these limitations, we developed a system for creating kilobase-scale genomic modifications that uses DNA oligonucleotides to direct the integration of a non-replicating plasmid. This method, Oligonucleotide Recombineering followed by Bxb-1 Integrase Targeting (ORBIT) was pioneered in *Mycobacteria*, and here we adapt and expand it for *E. coli*. Our redesigned plasmid toolkit achieved nearly 1000x higher efficiency than λ Red recombination and enabled precise, stable knockouts (≤134 kb) and integrations (≤11 kb) of various sizes. Additionally, we constructed multi-mutants (double and triple) in a single transformation, using orthogonal attachment sites. At high throughput, we used pools of targeting oligonucleotides to knock out nearly all known transcription factor and small RNA genes, yielding accurate, genome-wide, single mutant libraries. By counting genomic barcodes, we also show ORBIT libraries can scale to thousands of unique members (>30k). This work demonstrates that ORBIT for *E. coli* is a flexible reverse genetic system that facilitates rapid construction of complex strains and readily scales to create sophisticated mutant libraries.

## Introduction

The precise manipulation of bacterial genomes for constructing strains is indispensable for many scientific fields that study or use bacteria. Despite much progress in the model bacterium, *Escherichia coli*, molecular cloning or PCR is typically required to generate substrates for each newly targeted genomic locus, and researchers must still devote significant resources to initial strain construction. Beyond individual modifications, large pools of mutants can be created and used to ask a wide range of questions when coupled to modern DNA sequencing (1). However, the types of mutations available for genome-wide mutant libraries are limited, and there are currently no techniques that enable the precise kilobase scale manipulations typical of reverse genetics (e.g. gene deletion, GFP fusion) at high throughput. Consequently, there is a need for improved genetic tools that address these shortcomings and empower researchers to accomplish complex genetics rapidly, cheaply, and flexibly at scales from individual mutants to genome wide mutant libraries.

Reverse genetic approaches generally rely on specifying a genomic modification by introducing exogenous DNA with genomic homology regions into the cell. To avoid molecular cloning of unique plasmids for each desired mutation, λ Red recombination was established for *E. coli*, which uses more easily generated linear PCR fragments. Short homology arms (< 50 nucleotide (nt)) are added to an antibiotic marker using long PCR primers with overhangs and the product is electroporated (2). This method relies on exogenous recombination machinery encoded on a helper plasmid (Beta, Gam, Exo) and has been widely used to construct individual strains, including a single gene knockout library, the Keio collection (3). Despite many improvements and adaptations, λ Red is known to have important limitations, including relatively low recombination frequency (e.g. 1 mutant per 10^4-5^ cells), that preclude more demanding applications, and it is often necessary for researchers to use a variety of alternative tools (4–9).

Advances in DNA synthesis have fueled new high throughput methods that can specifically target many different genomic sites for modification. These techniques use commercially available DNA oligonucleotide (oligo) pools (< 1 million oligos), where each oligo encodes a short region of homology that can ultimately be used to direct a mutation. λ Red and various CRISPR methods have been adapted for high throughput applications by converting these oligo pools into double stranded cassettes or cloning them into plasmids (10–14). However, to our knowledge, none of these techniques have been successfully used for precise kilobase scale modifications at high throughput.

Oligo recombineering is a unique method for reverse genetics because it does not require molecular cloning or PCR, and instead uses chemically synthesized DNA oligos directly for performing genetic modifications (15). Typically, DNA oligos encoding short homology arms (<50 nt each or 100 nt total) are electroporated into cells expressing single stranded DNA annealing proteins (SSAPs) that bind to the DNA oligo and the complementary genomic region. Oligos are known to bind the lagging strand during DNA replication and are incorporated directly into a newly synthesized genomic strand (16–18). After an additional round of DNA replication, the mutation encoded in the oligo segregates as a newly formed mutant genome (19). This method is highly advantageous in that it is fast and convenient to rely directly on commercially available oligo substrates without additional molecular biology. Oligo recombineering is also relatively efficient, with single nucleotide changes exceeding 25% efficiency (20). However, most modifications do not confer antibiotic resistance or other selectable phenotypes, therefore high efficiency is required to reasonably find a mutant through colony screening. Consequently, oligo recombineering is limited to very high efficiency modifications, which tend to be short deletions (<100 bp), insertions (<10 bp) or nucleotide changes (21). Pools of recombineering oligos have been used for high throughput applications to make many single nucleotide modifications in individual strains (21), construct amino acid mutant libraries on plasmids (22), and generate single mutant libraries with inducible T7 promoters upstream of genes of interest (23). Therefore, the promise of oligo recombineering is speed, convenience, and high throughput compatibility, but without a selection strategy, it is limited to very small genomic edits.

A solution has recently emerged to use oligo recombineering to create kilobase scale insertions and deletions, by conferring antibiotic resistance through the integration of a plasmid, making lower frequency genetic modifications easily selectable. This method, Oligo Recombineering followed by Bxb-1 Integrase Targeting (ORBIT), was pioneered in *Mycobacteria* and was successfully used to make insertions or deletions at over 100 loci (24, 25). The key advancement was the inclusion of a short attachment site on the oligo (flanked by homology arms), which directs the integration (via Bxb-1) of a non-replicating plasmid containing a cognate attachment site into the newly modified genomic locus (Fig. 1A). The plasmid and the oligo are co-electroporated, combining this two-step process into a single transformation. In this manner, new genomic modifications can be directed with a DNA oligo without the need to change the payload (i.e. integrating) plasmid.

**Figure 1.**
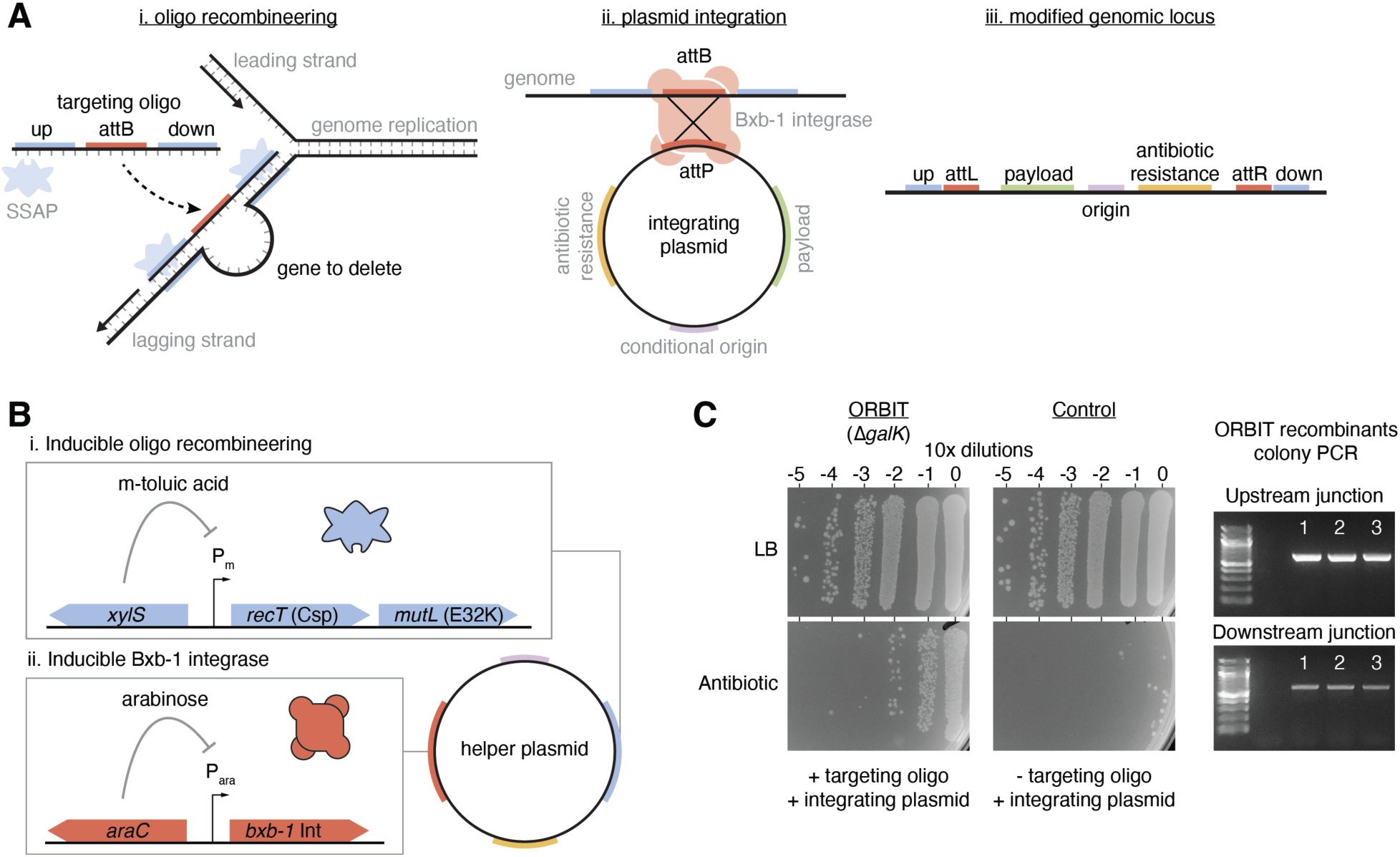
ORBIT overview and proof of principle. A) ORBIT is comprised of three steps. i) Oligo recombineering: A targeting oligo with the attB site and genomic homology arms specifying a mutation is integrated into the genome at the lagging strand through the action of the SSAP protein. ii) Plasmid integration: The newly incorporated attB site is recognized by the Bxb-1 integrase, which catalyzes recombination with the attP site on an integration plasmid. The integration plasmid cannot replicate in WT cells and carries an antibiotic resistance marker and other payloads, as needed. iii) Modified genomic locus: Following plasmid integration, recombinants are selected with the integrating plasmid antibiotic. B) The ORBIT helper plasmid (pHelper_Ec1_V1_gentR) consists of two separately inducible modules for oligo recombineering and plasmid integration. Module i expresses the SSAP, CspRecT, and the mismatch repair inhibitor allele MutL_E32K under the control of the xylS system with m-toluic acid inducer. Module ii expresses the Bxb-1 integrase under the control of the araC system with arabinose inducer. C) Serial dilution drip plates are shown for ORBIT transformations with and without a Δ*galK* targeting oligo and a kanamycin marked integrating plasmid. Many more kanamycin resistant colonies are observed in the presence of the targeting oligo. Putative recombinants were tested by PCR for the expected upstream and downstream junctions at the locus (n = 3 colonies).

Despite the advantages of ORBIT, there have not been any reports of a toolkit for use in the most common model bacterium, *E. coli*. Here, we adapt and expand on the ORBIT concept and develop an approach for achieving both low and high throughput kilobase scale genetics in *E. coli* K12 MG1655. We successfully constructed large deletions (up 134 kb) and insertions (up to 11 kb), single step double and triple mutants, markerless and scarless mutations and unprecedented gene knockout libraries (361 genes total), all directed by readily available DNA oligos. The concepts demonstrated here could be expanded in many different directions and this work advances the promise of ORBIT for achieving flexible genetics with a single toolkit, at low and high throughput, in diverse bacteria.

## Materials and Methods

### Strains and growth conditions

ORBIT experiments were performed in *E. coli* K12 MG1655 carrying an ORBIT helper plasmid. Helper plasmids were cloned into and maintained in *E. coli* DH5α, while integrating plasmids were maintained in BW25141 (Coli Genetic Stock Center, CGSC #7635), which is a pir+ host capable of replicating R6Kγ ori plasmids. Strains were grown in LB liquid cultures in 14 mL round bottom polypropylene culture tubes in a shaking incubator (Innova 42) set to 37°C or 30°C (for temperature sensitive strains) and 225 RPM. Overnight cultures were typically started directly from -80°C frozen glycerol stocks. As needed, cultures were plated on LB + 1.5% agar (RPI #L42020). Both plates and liquid cultures contained antibiotics at the following concentrations: kanamycin sulfate 35 µg / mL, gentamicin sulfate 15 µg / mL, chloramphenicol 30 µg / mL, ampicillin 100 µg / mL, carbenicillin 100 µg / mL. Carbenicillin was typically used in place of ampicillin due to its higher stability.

For quantifying ORBIT efficiencies, colony forming unit (CFU) counts were performed with a drip plate technique to fit serial dilutions on individual plates. Briefly, 150 µL of each recovery culture was added to the first well of a 96 well plate and serially diluted 7 times in neighboring wells (15 µL into 135 µL of phosphate buffered saline, PBS). PBS contained 137 mM NaCl, 10 mM PO_4_ (1.44 g / L Na_2_HPO_4_, 0.24 g / L KH_2_PO_4_), and 2.7 mM KCl, at pH 7.2. Dilutions were performed using multichannel pipettes, then 10 µL of each well was spotted onto a dry agar plate and tilted to form parallel drips. Plates were dried under a Bunsen burner and drips enabled accurate counting of 15-150 colonies per dilution. Efficiency was determined from CFU counts on LB and antibiotic media as follows: Efficiency = (CFU_antibiotic_ / CFU_LB_) * 100%.

Putative galactose and amino acid auxotrophs were phenotypically assessed on minimal media with sugars and / or amino acids added. M9 medium was used with glucose (0.4%) or galactose (0.4%), and 40 µg / mL of L-histidine, L-methionine, or L-leucine as needed. To test for a functional violacein pathway, the LuxR inducer, N-(β-Ketocaproyl)-DL-homoserine lactone (Sigma Aldrich #K3255), was added to 100 µM final from a 10 mM DMSO stock and supplemental tryptophan (100 µg / mL) was also added to the M9 glucose medium. M9 agar plates were prepared by autoclaving 15 g of agar with ∼780 mL of milliQ H_2_O. After autoclaving, 200 mL of prewarmed 5x M9 salts (64g Na_2_HPO_4_, 15g KH_2_PO_4_, 2.5g NaCl, 5g NH_4_Cl per L) were added to the agar which was cooled in a 60°C water bath. MgSO_4_ (2 mL of 1 M stock), CaCl_2_ (100 µL of 1 M stock), thiamine (100 µL of 0.5% stock), and the carbon source and required amino acids were added and the mixture was stirred on a hotplate stirrer on the bench to ensure no bubble formation. Then 20-25 mL of mixture was poured into plastic dishes.

For all pinned plates, colonies were picked into 100 µL of LB in 96 well plates using 200 µL pipette tips. 96 well plates were taped to the bottom of the shaking incubator and left to grow at 37°C. Once wells were obviously turbid, cultures were plated onto the appropriate M9 or LB antibiotic plates using a 48 pronged pinner (Millipore Sigma #R2383). The pinner was sterilized in 10% bleach, then washed in water, then 70% ethanol and flamed in between each sample. M9 plates were left to grow at 37°C for 1-2 days. Growth on M9 galactose often took particularly long. Plates were imaged on a gel documentation system (Fotodyne) with UVA light.

### Cloning procedures

To construct pHelper_Ec1_V1_gentR, the XylS inducible oligo recombineering module was cloned from pORTMAGE-Ec1 (a gift from George Church, Addgene #138474)(20), *bxb-1*, the origin and *sacB* were cloned from pKM461 (a gift from Kenan Murphy, Addgene #108320), the AraC inducible module was cloned from pKD46 (CGSC #7739) and the antibiotic marker was cloned from pMQ30 (2, 20, 24). All subsequent helper plasmids, including pHelper_noMutL_V2_gentR, were modified from pHelper_Ec1_V1_gentR as described in Table S1. pInt_attP1_kanR was constructed from pKM496 (a gift from Kenan Murphy, Addgene #109301) and pKD4 (CGSC #7632) and the attB site was changed to an attP site. All subsequent integrating plasmids were modified from pInt_attP1_kanR as described in Table S1. Fragments of the violacein operon were cloned from pBLxVi5 (a gift from Christopher Reisch, Addgene #167516) into pInt_attP1_kanR (26). The temperature sensitive Xis module for pInt_attP1_tsXis_sacB_kanR was cloned from pCP20 (CGSC #7629) and pBad-Xis/Int Reset Flipper (a gift from Drew Endy, Addgene #38207) (27). For pInt_LCS_kanR, the library cloning site with transcriptional terminators was cloned from pLibAcceptorV2 (a gift Guillaume Urtecho and Sriram Kosuri, Addgene #106250). In several cases synthetic terminators were added downstream of genes that appeared to lack transcriptional terminators (28).

Generally, Gibson assembly type reactions were used to construct plasmids by fusing 20 bp of homology as primer overhangs to PCR products. PCR reactions were performed using the high-fidelity polymerase Q5 (New England Biolabs - NEB M0492 or M0493) and purified using a DNA clean and concentrate kit (Zymo Research #D4014). Two or three fragment assemblies were performed using TEDA or a HiFi DNA assembly kit (NEB #E2621) (29).

Note that we tried to increase the yield for the integrating plasmids by using the copy up mutation pir116 host (BW19610), however we observed persistent issues with plasmid concatemers, possibly due to relaxed replication. Often, low yield integrating plasmids were midi or maxiprepped (Zymo Research #4200 / #4202) using the low copy number protocols, while higher yield replicating plasmids were miniprepped. Plasmids were fully sequenced by Primordium using Nanopore long reads. Annotated genbank files of plasmids are available as supplementary files (File S1) and at the GitHub repository (data/DNA_maps/plasmids).

### Targeting oligo design and ordering

Targeting oligos were designed to bind the lagging DNA strand and the lagging strand depends on the replichore. For *E. coli* K12 MG1655, since the origin of replication is at 3.92 Mb and the terminus is at ∼1.59 Mb, replichore 2 includes genomic positions >1.59 megabases (Mb) and <3.92 Mb and replichore 1 includes genomic positions <1.59 Mb and > 3.92 Mb (30). For replichore 1, the newly synthesized lagging strand is templated off the parental + strand, therefore the lagging strand targeted oligo should contain – strand homology sequence (Fig. S1A). For replichore 2, it is the opposite, so the targeting oligo should contain + strand homology sequence. For ORBIT, it is also important to decide which direction the attB site (and therefore the integrating plasmid) will go. Typically, attB sites were used in the same direction as the target gene. When taking both strands and attB directions into considerations, there are four possible targeting oligo structures depending on the genomic target, as shown in Figure S1B.

To simplify this process, we developed a web app that instantaneously generates targeting oligos that bind the lagging strand and face the same direction as a target gene (saunders-lab.shinyapps.io/ORBIT_TO_design_ecMG1655) (Fig. S1C)(31). This app has a simple genome browser for the *E. coli* K12 MG1655 genome, and users can jump to genes of interest using a searchable drop-down menu. Deletion (or other) targeting oligos are generated upon selection of a gene or clicking a gene in the browser window. Parameters such as homology arm length and attB direction can be customized and the targeting oligo sequence (5’ to 3’) can be copied and pasted directly into a commercial vendor website. We often used this app to design targeting oligos in conjunction with a DNA editor (e.g. Benchling) to ensure that the final ORBIT modified locus was correctly designed and to generate loci specific PCR primers for verification.

Typically, we used 120 nt targeting oligos from Millipore Sigma with no additional purification or modification (desalted only). For various experiments, we did order oligos with phosphorothioate bonds and tried different purifications (cartridge, PAGE, HPLC). We also ordered targeting oligos from Integrated DNA Technologies (IDT) and Synbio Technologies.

### ORBIT induced electrocompetent cell preparation

A complete protocol for making induced electrocompetent cells is included in the supplementary materials (Text S1). Briefly, strains with the helper plasmid were grown overnight with gentamicin (3 mL cultures). In the morning, cultures were diluted 1:1000 into larger volumes, typically 500 mL, with LB + gentamicin. Cultures were grown ∼3.5 hours to OD ∼0.3 and cultures were induced for oligo recombineering with 1 mM m-toluic acid (Sigma Aldrich #T36609) (1 M stock in 100% ethanol). After a 30 min induction continuing in the shaking incubator, flasks were put on ice and swirled periodically for ∼15 min. Cultures were distributed into 50 mL falcon tubes and centrifuged at 5, 000 rcf for 10 min in a pre-chilled centrifuge. Tubes were transported on ice to a cold room, supernatant was removed, and pellets were thoroughly resuspended in ice cold milliQ water with a serological pipettor (50 mL and then 10 mL). Tubes were inverted 5 times, then transported on ice and centrifuged again. This process was repeated two more times with 10% glycerol washes. The final wash supernatant was poured off and cells were resuspended in residual buffer typically yielded 50-100x concentrated cells in 10% glycerol. These cells were aliquoted into microfuge tubes placed open faced on ice (50 µL aliquots). Each aliquot was then chilled for >5 min on dry ice before placing in freezer boxes and placed at -80°C. Competent cells typically retained high efficiency for several months.

### ORBIT and λ Red transformation and recovery

A complete protocol for performing ORBIT transformations is included in the supplementary materials (Text S2). Frozen induced competent cell aliquots were taken from the -80°C box and thawed on ice for ∼10 min or until completely thawed. Then targeting oligo (2 µL of 25 µM stock – 1 µM final) and integrating plasmid (1 µL of 100 ng / µL) were added to each aliquot and aliquots were mixed 3 times with a 200 µL pipette before transferring to ice cold 0.1 cm electroporation cuvettes (Fisher #FB101). Cuvettes were tapped and inspected to ensure no bubbles or gaps were present and electroporated with standard *E. coli* settings (1.8 kV, Bio-Rad MicroPulser). With a 1000 µL pipette, 1 mL of recovery media (LB + 0.1-1% L-arabinose) was gently added to the cells and cells were transferred to the 3 mL recovery culture tubes. Entire experiments worth of cuvettes were transformed and transferred to recovery cultures on the benchtop. Then all culture tubes were placed in the shaking incubator for typically 1 hour before drip plating.

Generally, the standard conditions used for ORBIT transformations were 1 µM final of targeting oligo, 100 ng of integrating plasmid, with a 1 hour recovery in 3 mL of LB + 0.1% arabinose, however, many experiments deviated from these conditions, as needed. When using different sized integrating plasmids (with different violacein pathway fragments), we did not use 100 ng and instead used 1 nM final concentrations to account for the different molecular weights. For multi-mutant ORBIT experiments, targeting oligos were each added to 1 µM final concentration (i.e. 2 or 3 µM total oligo concentration for double or triple mutants). Similarly, when multiple integrating plasmids were used 100 ng of each was added to the ORBIT transformation.

For λ Red recombination, two 50 µL PCRs (Q5 hotstart) were performed for each targeted locus using 60 bp primers (40 bp homology overhangs) and pKD4 (1 ng) as the template for the kanamycin resistance marker. The reactions were checked on an agarose gel for a single band and reactions were pooled and cleaned and concentrated into a small volume of water. *E. coli* K12 MG1655 with pKD46 induced electrocompetent cells were prepared in a similar manner to the ORBIT induced electrocompetent cells described above. Briefly, cells were grown at 30°C and at OD ∼0.3 cells were induced for 30 min with 0.1% arabinose (instead of m-toluic acid).

Frozen competent cell aliquots were used for λ Red recombination in a similar manner as ORBIT competent cell aliquots. Aliquots were thawed for ∼10 min on ice and 2 µL of 150 ng / µL PCR product was added before electroporation as described above. Cells recovered in 3 mL LB at 37°C for 1 hour before plating.

### ORBIT mutant verification

ORBIT mutants were tested phenotypically, as described above, and molecularly with PCR and Sanger sequencing. Two different PCR strategies were used – a “junction PCR” that separately amplified the upstream and downstream junctions of the modified locus using a primer ∼250 bp away from the target site in the genome and another primer binding the integrating plasmid sequence. These PCRs confirmed the correct integration orientation and only showed bands for mutant strains and were used to test initial ORBIT mutants that still possessed the helper plasmid after a 1 hour recovery and plating. The second more stringent PCR strategy we used was a “spanning PCR” where the entire ORBIT modified locus was amplified with genomic primers ∼250 bp away from the target site. This spanning PCR was always expected to produce bands of different sizes from mutants and wildtype colonies. Once the helper plasmid had been cured, we used spanning PCRs for final mutant verification and Sanger sequencing, due to their higher stringency. In many cases, transformations were diluted 1:100 in LB + 0.1% arabinose + antibiotic and left to grow overnight at 37°C and streak plated the next morning on antibiotic + sucrose. This “overnight ORBIT” protocol served to cure strains of the helper plasmid quickly, facilitating more rapid spanning PCR and Sanger verification. However, ORBIT mutants were also cured of the helper plasmid by inoculating colonies overnight in LB + antibiotic and plating on LB + antibiotic + 7.5% sucrose the following morning.

Whole genome sequencing was performed on various ORBIT mutants to verify the accuracy of deletions and to quantify the background mutation rate. Genomic DNA was isolated with a gDNA miniprep kit (Zymo #D3024) and submitted to SeqCoast Genomics for short read sequencing (Illumina). Reads were compared to the U00096.3 genome sequence and variants were called using breseq (32).

### galK barcode sequencing and analysis

To create a targeting oligo containing random “N” barcodes, a Δ*galK* targeting oligo with 33 nt homology arms was designed with 16 N’s following the 38 nt attB site. The total oligo length was 120 nt and was ordered from Millipore Sigma’s standard oligo service. ORBIT was performed with this targeting oligo and standard conditions (1 µM oligo, 100 ng pInt_kan, 1 hour recovery in 0.1% arabinose). Approximately 300, 000 colonies were obtained from plating the 3 mL recovery culture (1 mL on 3 different plates). Colonies were resuspended in 1 mL of LB per plate and saved as glycerol stocks and cell pellets. Genomic DNA extraction was performed on cell pellets using a gDNA miniprep kit (Zymo #D3024). Amplicon libraries of the pInt_kan-gal locus junction were prepared in two rounds of PCR. The structure of the library and primers are shown in File S1 and the GitHub repository file galk_BC16N_orbit_seq_library.gb under data/DNA_maps/sequencing_libraries. Two 50 µL Q5 hotstart PCR1 reactions were performed with 1 µg extracted genomic DNA using the following program: 98°C for 30 seconds, then 15 cycles of 98°C for 10 seconds, 57°C for 20 seconds, 72°C for 20 seconds. Each reaction yielded a single band of ∼200 bp on a 2% agrose gel (and a high molecular weight smear of genomic DNA), which was gel extracted. Two 50 µL Q5 hotstart PCR2 reactions were performed with 10 ng of the gel extracted PCR1 product using the following program: 98°C for 30 seconds, then 5 cycles of 98°C for 10 seconds, 57°C for 20 seconds, 72°C for 20 seconds. The PCR2 reaction products were run on a 2% agarose gel, yielding a single band of ∼300 bp, which was gel extracted and subsequently clean and concentrated.

This amplicon library was loaded onto a MiSeq instrument at 7 pM (and 10% phiX control) with an Illumina Nano 300 cycle kit (reads 1 and 2 were 151 cycles). This sequencing run yielded 792, 034 paired reads, of which >99.8% passed quality filters (fastp)(33). Perfect flanking regions were extracted using tidysq in R and the intervening sequences that were exactly 16 bp long were assumed to be the random barcode regions (34). Approximately 110, 000 unique putative barcode sequences were recovered. These barcode sequences and their respective counts were provided to starcode to perform a conservative error correction using the following parameters “starcode -d 3 -s” to specify the maximum Levenshtein distance as 3 with the sphere clustering algorithm (35, 36). Complete analysis notebooks are available in the GitHub code repository under code/seq_data_processing/galK_BC_16N.

### IDT targeting oligos and transformation

The four 120 nt targeting oligos used previously for deletion of *galK*, *hisA*, *metA*, and *leuD* and 23 transcription factor deletion oligos (originally designed for the Twist oligo pool - see below for details) were ordered as an IDT oPool at 50 pmol / oligo scale. The pooled oligo library was resuspended to 100 µM total oligo in water. This targeting oligo pool was used for ORBIT with standard conditions (1 µM total oligo concentration (final), 100 ng pInt_kanR, 1 hour recovery in 0.1% arabinose). Approximately 5, 000 colonies were obtained from plating 100 µL of recovery culture in duplicate transformations. Colonies were resuspended in 1 mL LB and glycerol stocks and cell pellets were saved. See below for mutant library sequencing information.

### Twist targeting oligo design, processing, and transformation

We first searched the Ecocyc database for all proteins categorized as DNA binding transcription factors and obtained their start and stop positions, and the same was done for all annotated small RNA genes (see notebooks in the GitHub repository under /code/twist_oligo_design)(37). In Python, functions were written to create 128 nt targeting oligos that bind the lagging DNA strand for arbitrary genomic positions. This function was called for each target genes using the U00096.3 genome sequence (matching Ecocyc), resulting in targeting oligos containing upstream and downstream homology for each target gene (and the attB site).

To avoid disrupting nearby (or overlapping) upstream and downstream genes, transcription factor targeting oligos were designed to delete the sequence starting 6 bp from the start to 21 bp before the end of each gene. Because small RNA genes were shorter, they were entirely deleted from the start to stop positions. To avoid potential PCR issues, we chose to include the forward attB sequence (ggcttgtcgacgacggcggtctccgtcgtcaggatcat) in every 5’ to 3’ synthesized oligo and never the complementary reverse sequence, which would have formed a strong 38 bp duplex structure. Therefore, ORBIT integrations did not face the same relative direction when taking into account the different replichores and gene directions. Next, we added orthogonal 20 and 24 nt primer binding sites containing MboI and BtsI-V2 recognition sequences respectively (38). These sites were designed such that the targeting oligo sequences could be amplified with PCR, but then the primer sites could be entirely cleaved off (23). These targeting oligo sequences with primer sites were ordered in triplicate in a large pool from Twist Biosciences with many other unrelated experiments.

Upon receiving the oligo pool, it was resuspended according to the manufacturer’s guidelines and the three subpools were independently amplified using their specific orthogonal primer sets (see Table S2). PCR was performed in two stages and different cycle numbers were tested to use as few cycles as possible and limit unwanted products. Products were assessed on 2% agarose gels. PCR1 used 1 µL of the original oligo pool diluted 1:20 with 20 cycles (Q5 hotstart polymerase, 98°C – 15 sec, 64°C – 20 sec, 72°C – 20 sec). PCR2 was performed with 10 ng of purified PCR1 product for 7 cycles (same parameters as PCR1). Note that the forward primers had two phosphorothioate bonds at the 5’ end and the reverse primers had 5’ phosphate groups. These modifications promoted the degradation of the reverse (i.e. bottom) strand by lambda exonuclease. This digestion was performed on 1-3 µg of the PCR2 product with 1 µL of exonuclease (NEB #M0262) at 37°C for 30 min before heat inactivation. Digestion products (presumably single stranded DNA) were purified with a ssDNA clean and concentrate kit (Zymo #7010).

Next guide primers were annealed to the single stranded DNA (targeting oligo with primer overhangs), making the restriction sites duplex DNA and therefore suitable substrates for endonucleases. Annealing was performed with 4 µM (final) of each guide oligo and 0.5-2 µg of template DNA in 1x Cutsmart buffer (NEB) using a slow temperature gradient (95°C – 60°C at 0.1°C / sec, 60°C for 3 min, 60°C – 50°C at 0.1°C / sec, 50°C for 3 min, 50°C – 37°C at 0.1°C / sec, 37°C for 3 min). Then 10 units (2 µL) of each restriction enzyme (MboI and BtsI-V2) was added to the 50 µL reaction and left at 37°C overnight. Products were purified using the ssDNA clean and concentrate kit and loaded on 10% TBE-Urea gels (BioRad #4566033) with sample buffer (Thomas Scientific #C995V39) and run at 200V for ∼30 minutes. Gels were stained with 1x Sybr gold (ThermoFisher #S11494) for 20 minutes and then imaged. For size comparison, 120 nt and 90 nt targeting oligos were run alongside digestion products. This process initially yielded incomplete products, so it was repeated three times, ultimately yielding 25-50 ng of final product.

These processed targeting oligos were used for a standard ORBIT experiment with 50 µL induced electrocompetent cell aliquots and 100 ng of pInt_attP1_kanR – approximately 5 µL of targeting oligo product was added to each transformation. Cells were recovered for 1 hour in 3 mL LB + 0.1% arabinose – CFU drip plates were performed and 1 mL of recovery culture was also plated on 3 separate kanamycin plates for each transformation. Colonies were resuspended in LB and pooled glycerol stocks were made and cell pellets were saved for genomic DNA extraction.

### Mutant library sequencing and analysis

Library prep for ORBIT mutant pools was performed in a manner analogous to Tn-Seq (39, 40). Genomic DNA (gDNA) was extracted from mutant library cell pellets (IDT oPool and Twist libraries) using a gDNA miniprep kit, as for the *galK* barcode experiment above. For each library, approximately 50 µL of gDNA (2.5 – 5 µg) was sheared with a sonicator (Covaris S2) in a 130 µL cuvette with the following parameters: Duty cycle = 10%, Intensity = 5, Cycles per burst = 100, Cycle time = 60 sec, Cycle number = 3, Water bath temp = 5°C. The fragmentation was assessed on an agarose gel and fragments between 150 and 350 bp were gel extracted (Zymo #4007). End repair was performed following the manufacturer’s instructions and purified (NEB #E6050). C-tailing was performed with Terminal Transferase (NEB #M0315) and 400-800 ng of end repaired gDNA using a 1:20 dCTP:ddCTP mixture (9.5 mM dCTP Millipore Sigma #3732738001, 0.5 mM ddCTP Promega #U1225) to generate ∼20 bp C-tails.

For each ORBIT integration, there are two genome:pInt junctions, and both needed to be checked to verify that a correct deletion occurred. Therefore, two sequencing library preps were carried out for each mutant library to separately amplify these two junctions (referred to as downstream / upstream or left / right). After purification, PCR1 was performed with 200 ng of C-tailed DNA using a primer that binds pInt_kanR and a polyG primer (with overhang) that binds the polyC tail using Q5 hotstart polymerase (98°C – 15 sec, 62°C – 20 sec, 72°C – 30 sec, 25 cycles). Illumina compatible overhangs were added with PCR2 using 1 µL of the unpurified PCR1 product (98°C – 15 sec, 63°C – 20 sec, 72°C – 30 sec, 15 cycles). Products between 250 and 500 bp (350 to 600 bp for right side) were gel extracted and served as the final sequencing library.

Libraries were quantified with Qubit fluorometry (Thermofisher # Q32851), diluted to 2 nM and pooled in a 1:3:3:3 ratio, since the oPool contained fewer mutants than the Twist libraries. Libraries were denatured and diluted to 7 pM (junction 1) or 9 pM (junction 2) before loading on a MiSeq instrument (illumina) with a Nano 300 cycle kit (and 10% phiX control). The two pInt:genome junction libraries were run on separate flowcells. The structure of these libraries is diagrammed in genbank files in the supplement (File S1) and in the GitHub repository under data/DNA_maps/sequencing_libraries. To analyze the data, the pInt sequence that should be present in each read was used as an adapter sequence in Cutadapt, and the remainder of the read was assumed to be the adjacent genomic sequence (see code/seq_data_processing/ in the GitHub repository for fully documented analysis) (41). Read 1 and read 2 were 151 cycles, however, only read 1 information was used, since it was higher quality and contained the conserved pInt sequence. For the first junction (i.e. left / downstream), 44, 000 to 137, 000 reads were obtained for each mutant pool that passed initial quality filters (fastp defaults) and 64 – 73% of these reads had the correct pInt sequence with enough genomic homology to map (≥ 20 bp)(33). Of these reads with putative genomic homology adjacent to the ORBIT integration, 83-85% mapped to the *E. coli* MG1655 genome with bowtie2 (using --very-sensitive-local)(42). For the second junction (i.e. right / upstream), more reads were obtained (82, 000-237, 000 fastp filtered reads), 55-60% of reads passed the Cutadapt step, and 79-83% of reads mapped to the *E. coli* genome. The biggest loss of reads through this pipeline was due to reads being too short (< 20 bp genomic homology), likely because of the initial size distribution of genomic fragments and unexpectedly long C-tails.

Once genomic positions adjacent to the pInt sequences were mapped, the start positions could be compared to the designed targeting oligo homology arms. Every read was assigned to the nearest designed TO position, which allowed for the calculation of the fraction of “perfect” reads that exactly matched an expected start position. Designed TOs for which perfect reads could be found in both the junction 1 and junction 2 libraries were identified, and those that had zero perfect reads in the libraries were considered absent. The absent target genes were further examined by classifying them as essential, perturbing or nonessential using Ecocyc essential gene datasets (LB) and manual loci inspection (37). Complete read processing and analysis notebooks are available in the GitHub repository under code/seq_data_processing/.

### TO library sequencing and mutant abundance analysis

To get the abundance of the ORBIT targeting oligos in the original Twist oligo pool, we prepared a sequencing library starting with the PCR1 Twist products (see above). PCR1 Twist products were amplified in two rounds to add Illumina compatible overhangs. First, 10 ng of the original Twist products were used for this PCR1 (98°C –15 sec, 64°C – 20 sec, 72°C – 20 sec, 7 cycles). Products were purified and 10 ng were added to PCR2, which provided the final overhangs (98°C – 15 sec, 64°C – 20 sec, 72°C – 20 sec, 8 cycles). This reaction yielded the expected size band (∼300 bp), which was gel extracted and served as the final sequencing library. Libraries were diluted to 2 nM, pooled 1:1:1, and denatured and diluted to 7 pM. Sequencing was performed with 10% phiX on a MiSeq instrument with a Nano 300 cycle kit. Read 1 was set to 179 cycles and read 2 was set to 123 cycles. Library structure is shown as a genbank file in the GitHub repository under data/DNA_maps/sequencing_libraries. Only read 1 was used, because it contained full length amplicon information and was higher quality. 202, 000-330, 000 fastp filtered reads were obtained for each TO pool, of which ∼91% were the exact expected length of 179 bp. The function “vcountPDict” from Biostrings was used to find perfect matches between sequencing reads and designed TOs (∼90% of total reads)(43). Complete analysis notebooks are available in the GitHub repository under code/seq_data_processing.

TO abundances were compared to mutant library read counts, along with a number of other oligo parameters using standard tools in R, and linear models were fit with the “lm” function (44, 45). Oligo free energy values were calculated using XNAstring (46). Analysis notebooks walk through the creation of all supplemental and main figures that have computationally derived content (see GitHub repo code/main_figs and code/supp_figs).

## Results

### Establishing the ORBIT concept in E. coli

The original ORBIT plasmids were designed for use in *Mycobacteria*, so we started by constructing a series of new plasmids designed for use in *E. coli* and included several improvements. ORBIT requires cells to carry out two sequential actions 1) oligo recombineering and 2) plasmid integration (Fig. 1A). We constructed a new helper plasmid (pHelper_Ec1_V1_gentR) that contains separately inducible modules that correspond to these two steps. Oligo recombineering is mediated by the SSAP, CspRecT, and efficiency is further enhanced by suppressing mismatch repair with the dominant negative protein MutL E32K (20, 47) (Fig 1B). This oligo recombineering module, termed pORTMAGE-Ec1, is the most efficient SSAP system for *E. coli* published to date, and the operon is induced from the helper plasmid by the XylS / m-toluic acid system (20). Plasmid integration is catalyzed by the Bxb-1 integrase and it is induced from the helper plasmid by the AraC / arabinose system (Fig 1B). Collectively, this forms a convenient system where one can briefly induce efficient oligo recombineering and subsequently induce Bxb-1 catalyzed integration for longer periods without continued mismatch repair suppression. The helper plasmid also contains the *sacB* gene, which allows for removal of the plasmid with sucrose counter selection.

The other plasmid required for ORBIT is the integrating (i.e. payload) plasmid, which is incorporated into the genome at the new att site formed by the targeting oligo (TO). This integrating plasmid should not replicate in wildtype cells, so we constructed a simple oriR6Kγ plasmid that only replicates in specific host strains that have the *pir* gene, which encodes for the plasmid replication protein. Normal molecular cloning worked with this plasmid – it only required transformation into the appropriate *pir+* host strain. We created a series of “pInt” plasmids with different antibiotic markers, att sites and payloads, that were used for various purposes. Unlike the original ORBIT system, we chose to put the longer attP site (48 bp) on the integrating plasmid, and the shorter attB site (38 bp) on the targeting oligo. This lends flexibility for targeting oligos to have longer homology arms or shorter overall length as needed.

We tested the functionality of our new ORBIT system in *E. coli* by attempting to knock out the ∼1 kb gene, *galK* (galactose kinase), with a targeting oligo containing upstream and downstream homology arms and the attB site. To quantify the on target and off target rates for the integrating plasmid, we performed transformations with and without the targeting oligo and plated serial dilutions on permissive (LB) and selective (LB + kanamycin) media (Fig. 1C). By counting colony forming units, we calculated the efficiency of obtaining kanamycin-resistant colonies and observed a putative on-target rate ∼100x higher than the off-target control, strongly suggesting that the integration of the plasmid was enhanced by the addition of the targeting oligo. We confirmed that resistant colonies contained the expected genome modification with PCR of the upstream and downstream junctions and Sanger sequencing (Fig. 1C).

### Optimization for efficient genomic deletions

We next wanted to optimize ORBIT for high efficiency, since efficiency ultimately determined what challenging applications would be possible. Gene deletions and insertions are common genome modifications; however, based on previous literature, we assumed that deletions would be less efficient than insertions and were therefore a more challenging starting point for optimization (21). The basic ORBIT protocol (Fig. 2A) is comprised of only a few steps: 1) Prepare m-toluic acid induced electrocompetent cells from the helper plasmid-containing strain. 2) Co-electroporate the targeting oligo and integrating plasmid. 3) Recover cells in arabinose inducer. 4) Plate cells on antibiotic agar to select for recombinants. Parameters within each one of these steps were tested to develop an optimal protocol, and to provide users with a reasonable intuition about what experimental variables matter. We focused on optimizing efficiency (i.e. frequency) and we assessed accuracy (how many colonies were correct) separately.

**Figure 2.**
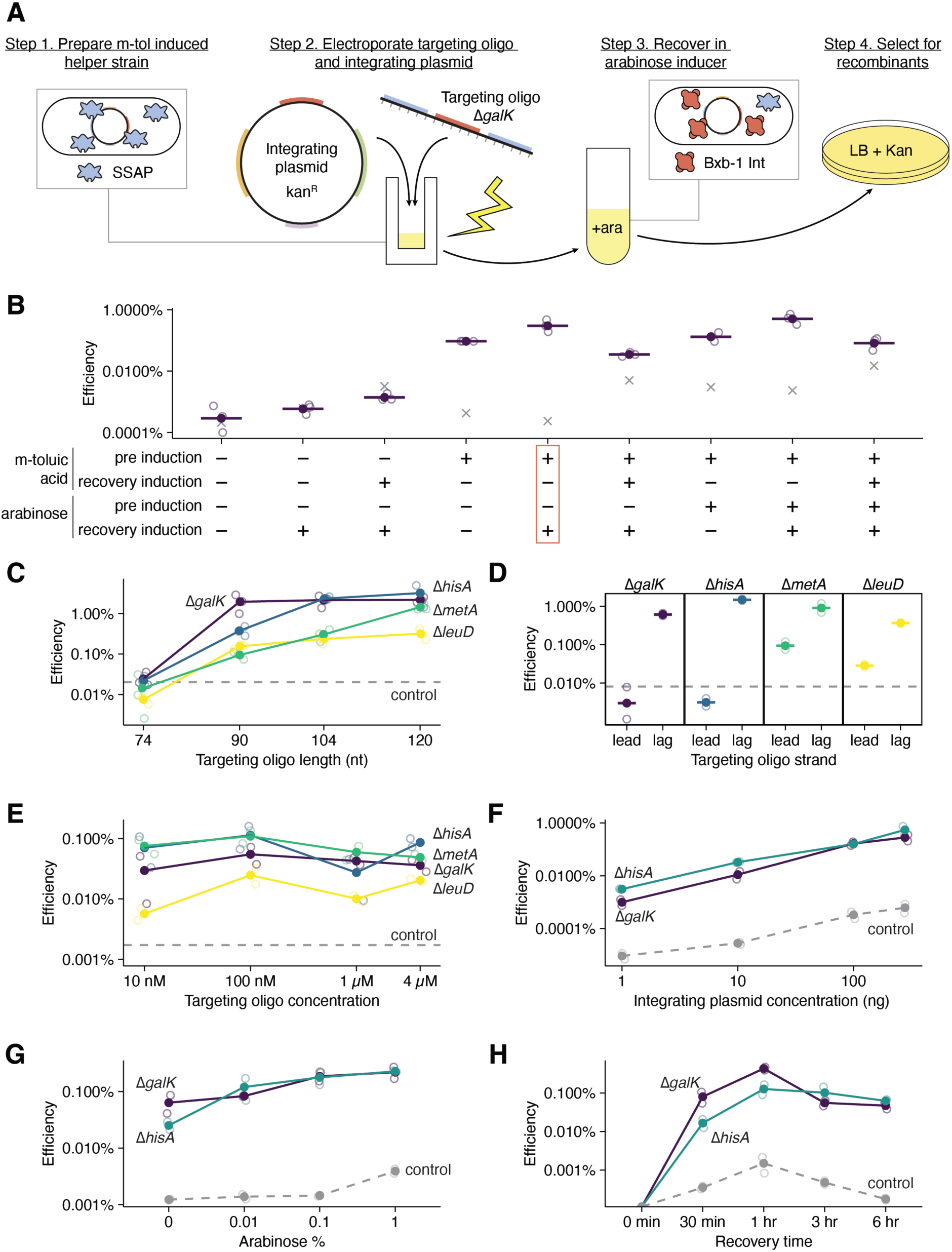
ORBIT protocol optimization. A) Diagram of the ORBIT protocol showing that m-toluic acid induced electrocompetent cells are co-transformed with integrating plasmid and targeting oligo in a single step, followed by recovery in arabinose and plating for recombinants. B) Helper plasmid induction conditions were tested for efficiency by drip plating on LB / kanamycin. M-toluic acid and arabinose inducers were included before electrocompetent cell prep (pre-induction) and / or during the recovery phase (n = 3 transformations per condition). A 90 nt Δ*galK* targeting oligo was used. Gray x’s indicate negative controls without targeting oligo. C) The effect of total targeting oligo length (including 38 nt attB site) was tested at all 4 target loci (n = 3 per condition). D) Targeting oligos against the lagging strand were compared to oligos against the leading genomic strand (i.e. reverse complement) (n = 2 per condition). E) Targeting oligos for each locus were tested at different final concentrations in 50 µL cell aliquots (n = 3 per condition). F) Integrating plasmid (pInt_attP1_kanR) was added in different amounts with Δ*galK*, Δ*hisA* or no targeting oligo (n = 2 per condition). G) Arabinose concentration during the recovery phase was varied with Δ*galK*, Δ*hisA* or no targeting oligo transformations (n = 2 per condition). Unless otherwise stated, Δ*galK*, Δ*hisA*, Δ*metA* Δ*leuD* targeting oligos were 120 nt and 100 ng pInt_attP1_kanR was used. Open circles represent replicate transformations and filled circles are mean values. Dashed gray lines are negative control transformations without targeting oligo added, but with integrating plasmid.

We started by testing induction conditions for our helper plasmid (Fig. 2B). We grew *E. coli* MG1655 with pHelper_Ec_V1_gentR and induced with either m-toluic acid (oligo recombineering), arabinose (bxb-1 integrase) or both for 30 min before pelleting and washing to make electrocompetent and storing at -80°C. Then we transformed the different “pre-induced” cells with a Δ*galK* deletion oligo and recovered cells in either m-toluic acid, arabinose or both before plating. This yielded results for every possible induction scheme and largely confirmed that our helper plasmid mediated ORBIT as expected. Without pre-induction of electrocompetent cells with m-toluic acid, efficiency was very low, while all conditions that included m-toluic pre-induction resulted in substantially higher efficiency. Adding m-toluic acid to the recovery media reduced efficiency, likely due to the prolonged induction of MutL, which could cause higher mutation rates and may be toxic. Induction of the *bxb-1* module (arabinose) was not strictly required, likely because of leaky expression, however the highest efficiencies were observed when arabinose was included in the recovery media. These results show that our helper plasmid successfully facilitates the sequential concept of ORBIT, where oligo recombineering takes place first to introduce the att site, then plasmid integration occurs to provide stable antibiotic selection.

Next, we focused on optimizing the targeting oligo. The original ORBIT work used oligos 148-188 bp in length, but we found that much shorter oligos work well with our *E. coli* system. For these optimization studies, we used our targeting oligo for deletion of the *galK* gene, as well as oligos targeting single gene deletions of amino acid biosynthetic genes *hisA*, *metA* and *leuD* whenever possible. At all 4 targeted gene loci, 90 nt oligos (26 nt homology arms + 38 nt attB) yielded efficient ORBIT well above background controls (Fig. 2C). At some loci (e.g. *metA*), significantly higher efficiency was obtained with longer homology arms, therefore we subsequently used 120 nt targeting oligos (41 nt homology arms + 38 nt attB) for most assays, because the price difference was negligible. Past work on oligo recombineering in *E. coli* has shown that oligos targeting the lagging strand during DNA replication are typically more efficient, suggesting that oligos are incorporated like an Okazaki fragment into the newly ligated lagging DNA strand (15). Consistent with this model, we observed at all 4 targeted loci that lagging strand targeting oligos were strikingly more efficient than leading strand ones (Fig 2D). Past work also recommended using phosphoorothioate (PO) bonds for recombineering oligos to make the single stranded DNA resistant to endogenous exonucleases (21). For oligos targeting the *galK* locus, we found only minor increases in efficiency by adding PO bonds (Fig S2A).

Phosphoramidite chemistry-based DNA oligo synthesis is known to yield larger fractions of incomplete products as oligo lengths increase, therefore DNA synthesis companies often recommend paying extra for high quality purifications to obtain more full-length sequences. We tested different oligo purifications (desalting, cartridge, HPLC, PAGE) for the *galK* deletion oligo and found that PAGE purification did increase efficiency (∼2.5x), but purification was not necessary for most applications (Fig S2B). We also tested 90 nt targeting oligos for all 4 loci from 3 different suppliers with different price points ($7-$18 per oligo) and found obvious differences, suggesting variable oligo quality (Fig S2C). Typically, we used 1 µM final of targeting oligo (assuming 50 µL aliquots), but we found that efficiency does not increase monotonically with increasing oligo concentration. Instead, we found that 10 nM oligo supports efficiency above background, and 100 nM oligo was consistently the most efficient concentration (Fig. 1E).

Beyond the targeting oligo, we also tested the effect of the integrating plasmid concentration on efficiency, and we saw a consistent rise in efficiency with higher concentrations (Fig. 1F). Interestingly, we also observed a rise in the putative off target rate with higher integrating plasmid concentrations, which suggests that the on vs. off target rate is largely invariant to the integrating plasmid concentration. We routinely transformed 100 ng of pInt_kanR, but this result suggests much less integrating plasmid could be used for routine applications with little downside.

Next, we specifically examined the recovery conditions following electroporation. We found that although efficiency is well above background without inducing Bxb-1 at all, adding up to 1% arabinose greatly increased efficiency. We observed that recovery times of ∼1 hour yielded the highest efficiency, but recovery times from 30 minutes to 6 hours also worked reasonably well. This time interval seemed surprisingly short for both the oligo recombineering and plasmid integration to occur before plating. However, we later discovered that our model of this process may be incomplete and that a significant fraction of plasmid integration may take place once cells have already been plated (Fig. S3A-B). Overall, our results suggest the following protocol parameters:

- ≥90 nt targeting oligo with attB against lagging strand (120 nt is our standard).
- ≥10 nM targeting oligo concentration (1 µM is our standard).
- ≥10 ng integrating plasmid (100 ng is our standard).
- 0.1-1% arabinose for 1 hour recovery (or overnight – see Fig. S3C-F).
- Eliminate helper plasmid when ready to use strain.

Our optimized protocol allows for the use of cheap, unpurified, and relatively short DNA oligos, small amounts of integrating plasmid and quick experimental times compatible with routine lab capabilities.

### Benchmarking and troubleshooting with single gene deletions

To demonstrate the utility of our protocol, we compared the efficiency of ORBIT at the 4 test loci to the current standard technique, λ-Red recombination (plasmid pKD46). λ-Red requires PCR to amplify an antibiotic resistance cassette with primers containing homology arms, so we used primers with nearly identical homology arms to the ORBIT targeting oligos. We performed 2x 50 µL PCRs with the appropriate primers for each locus and purified the products, ultimately transforming 300 ng of PCR product. ORBIT efficiencies were 2-3 orders of magnitude higher than the comparable λ-Red modifications (309x-1314x higher) (Fig 3A-B). This dramatic increase in efficiency is likely caused by several factors. First, it is significantly easier to obtain high concentrations of the ORBIT targeting oligo (direct from manufacturer tube) than a PCR product. However, the more important factor is likely that ORBIT relies on more efficient single stranded DNA recombineering, while λ-Red uses double stranded DNA, which is subsequently converted to a single stranded intermediate by the Exo protein (15, 16). Further, the ORBIT SSAP, CspRecT is known to be more efficient than the λ-Red SSAP, Beta (20).

**Figure 3.**
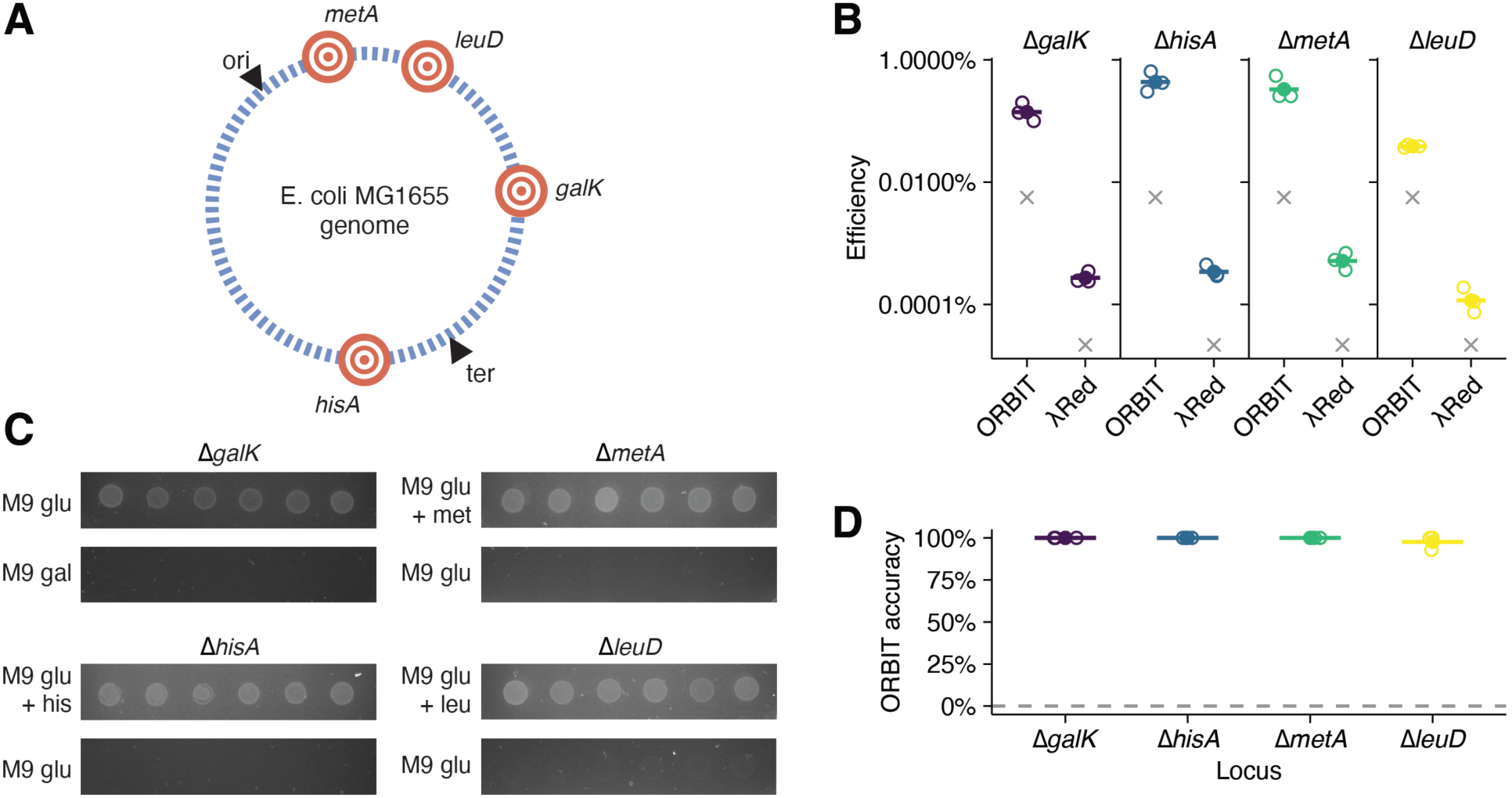
Benchmarking ORBIT. A) Genomic diagram of the four targeted loci. B) Comparison of ORBIT and λ-Red recombination (plasmid pKD46) at four loci. 300 ng of kanamycin resistant PCR amplified products (40 bp homology arms) were used for λ-Red. Open circles represent individual transformations and filled circles show mean values (n = 3 per condition). Gray x’s show negative controls without targeting oligo (ORBIT) or without PCR product (λ-Red). C) ORBIT mutants show expected growth defects on minimal media. D) Accuracy of mutations were assessed by growth phenotype on minimal media. For each transformation (n = 3), 14 separate colonies were assayed and scored for growth (yes or no) to obtain % accuracy displayed as open circles. Filled circles show the mean value of the open circle replicates. Gray dashed line shows phenotypic scores for colonies from negative control transformations in B.

Next, we assessed the accuracy of the kanamycin resistant colonies obtained from ORBIT. We grew colonies in 96 well plates and pinned cultures onto permissive or selective plates, based on the putatively deleted metabolic gene (i.e. M9 + glucose/galactose for *galK*, M9 glucose +/-histidine for *hisA*, +/-methionine for *metA*, and +/-leucine for *leuD*). Using this phenotypic assay, we scored 14 colonies, from triplicate transformations for growth on the selective media. We observed only a single *leuD* colony with an incorrect phenotype out of a total of 166 colonies tested, yielding average phenotypic accuracies of 100% for the deletions of *galK*, *hisA*, and *metA*, and 97.6% for the deletion of *leuD* (Fig. 3C-D, Fig. S4A). Colonies from each replicate were also tested by PCR and confirmed to contain the ORBIT integrating plasmid in the correct orientation (Fig. S4B-C). Therefore, our ORBIT method seems to be a highly capable reverse genetic system that can efficiently make precise gene deletions.

Despite this success, we did encounter a puzzling and persistent issue when trying to Sanger sequence PCR products that spanned the entire modified locus – sequencing results often looked perfect, except for very specific positions near the attP portions of the attR/L sequences. We found that a persistent lower band was present, which contained the modified locus with attB, but without the integrated plasmid. We discovered that by curing the helper plasmid, the lower band was eliminated. We confirmed that the helper plasmid was the source of this issue, by testing a Δ*galK* mutant that was cured of pHelper and then retransforming this strain with pHelper, which caused the lower band to reappear (Fig. S3C-F). This suggests that pHelper-derived leaky expression of the Bxb-1 integrase excised the integrated plasmid at a low, but detectable rate, which confounded our Sanger sequencing. Therefore, when preparing mutants for experiments or sequencing, we suggest diluting recovery cultures 1:100 in LB + kanamycin + 0.1% arabinose overnight and then plating on LB + kanamycin + sucrose the next day to immediately cure the helper plasmid. Once strains were cured of the helper plasmid, the ORBIT integrations at the *galK, hisA, metA,* and *leuD* loci were stable without antibiotic selection for at least 70 generations of liquid culture passaging (Fig. S4D).

We also performed whole genome sequencing on our Δ*galK* mutant and discovered that it harbored an unexpectedly high number of single nucleotide polymorphisms (SNPs), in addition to the intended deletion. We suspected that random mutations were occurring because of the *recT* or the *mutL* genes on the helper plasmid, so we tested three strategies to reduce the background mutation rate. 1) We swapped CspRecT for λ-Red Beta, 2) we removed the *mutL* gene from the helper plasmid and 3) we added strong degron tags to both *recT* and *mutL* to reduce their protein levels following induction. Figure S5A-C shows that all three new versions of the helper plasmid worked with reasonable efficiency to make our standard deletion mutants. Next, we performed whole genome sequencing on *galK*, *hisA* and *metA* mutants made with the original (V1) helper plasmid and the three new derivatives. We observed that the original helper plasmid strains again accumulated many mutations (3-8 SNPs) beyond the ones present in the WT parent. All three strategies showed improvement in the mutation rate, however, the best result was the “no MutL” helper plasmid, which showed zero new mutations across all three mutant strains (Fig. S5C-D). This result made sense, since *mutL* serves to inhibit part of the mismatch repair pathway (47). During this troubleshooting phase, we also discovered two SNPs present in the original helper plasmid *bxb-1* gene and a suboptimal ribosome binding site.

Therefore, we decided to make a version 2 of the helper plasmid that fixed the *bxb-1* issues and removed *mutL*. We remade competent cells and compared pHelper_v1 and pHelper_v2 efficiency on different days and we saw that V2 was slightly more efficient, although it also showed a higher pInt only off target rate (Fig. S5E). We confirmed that all four deletion mutants were made properly by PCR, with only a single *leuD* mutant seemingly incorrect for a total of 35/36 correct colonies (Fig. S5F). Consequently, we believe that pHelper_V2 will be a useful ORBIT tool moving forward to avoid elevated background mutation rates. However, for the remainder of this work, we continued to employ the V1 helper plasmid, since all previous work was with this plasmid.

### Generating large genomic deletions and insertions using ORBIT

Past work suggested that smaller genomic modifications would be more efficient than larger genomic modifications (21). Since our single gene deletions were achieved at relatively high efficiency, we reasoned that ORBIT may be useful for particularly long deletions or large insertions, which may be difficult to construct with λ-Red (48, 49). To test the effect of deletion size, we designed targeting oligos to delete smaller (100 bp) and larger (6-13 kb) regions around 3 test loci (*hisA*, *metA*, *leuD*) (Fig 4A). We also designed a larger set of deletions spanning from 1 to 49 kb surrounding the *galK* locus, making sure to avoid obviously essential genes. As expected, efficiencies broadly declined as deletions became larger, however, sufficient colonies were easily obtained from each transformation (Fig 4B). Interestingly, efficiency did not decline monotonically for increasing deletion size at each locus, suggesting other targeting oligo parameters play important roles in determining efficiency. We phenotyped these deletion mutants (n=7-8 for duplicate transformations) on M9 agar as described above and observed accuracies of 87-100% for all 14 mutants, with only 8 putatively incorrect colonies out of a total of 206 tested (Fig 4C, S6). Representative colonies were also tested by PCR and Sanger sequencing, which confirmed that recombinants had the exact expected locus structures (Fig. S7-8).

**Figure 4.**
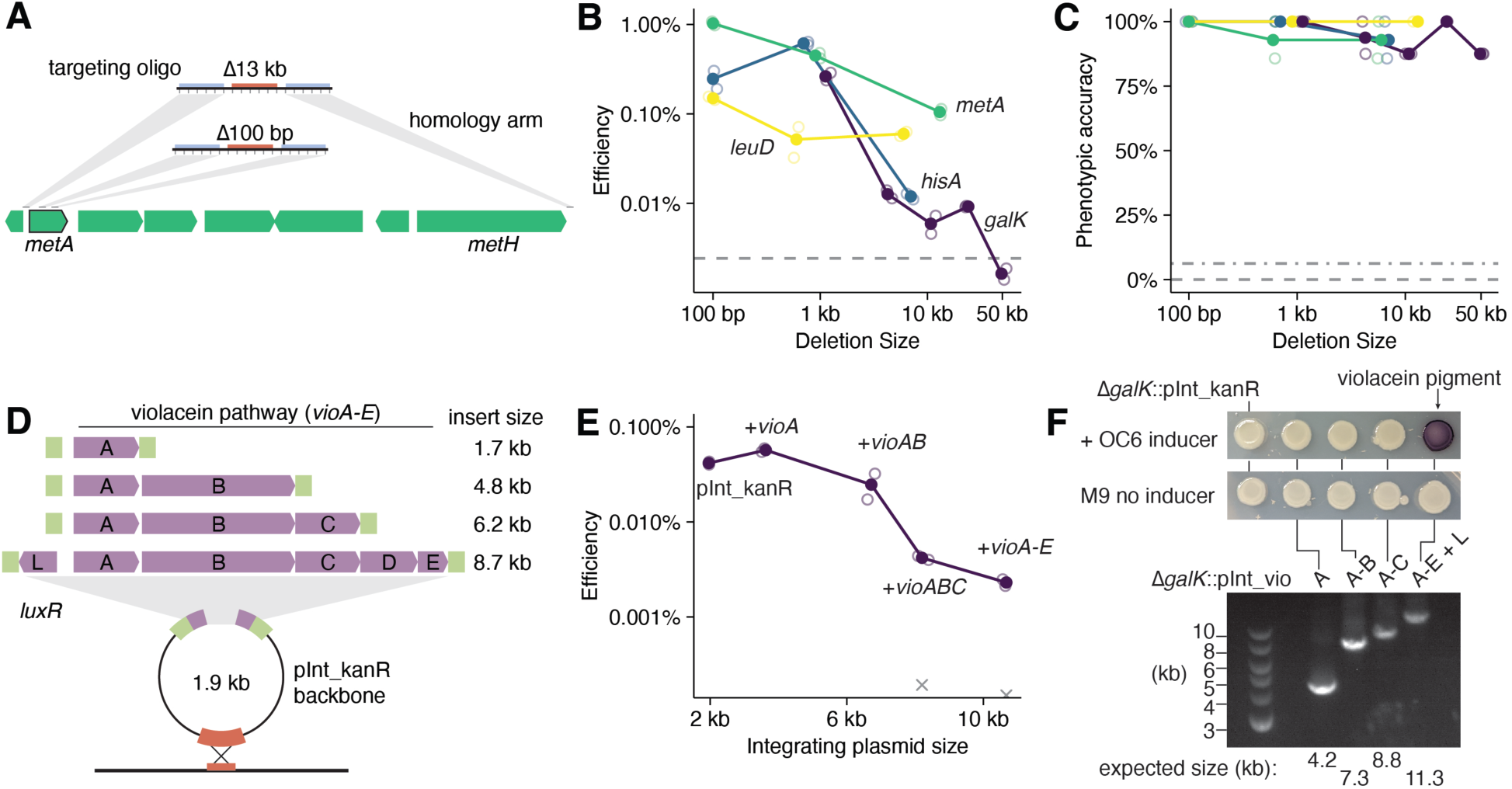
Large deletions and insertions with ORBIT. A) Targeting oligos were designed to delete different sized genomic regions, for example, around the *metA* locus. B) The efficiency of each different sized mutation was tested with duplicate (n = 2) transformations (open circles). Filled circles show mean values and the gray dashed line shows transformation efficiency without a targeting oligo. C) For each transformation, 7 colonies were individually phenotyped on minimal medium to obtain accuracy scores for each mutation (open circles). Filled circles show mean values. Dashed gray lines show phenotypic scores for colonies obtained from negative control transformations – controls for *galK* are shown separate from the other three loci. Different sized fragments of the violacein pathway (under *luxR* inducible control) were cloned in the integrating plasmid backbone of pInt_attP1_kanR. E) Different sized integrating plasmids were tested for efficiency with the Δ*galK* targeting oligo with duplicate transformations (n = 2, open circles) and filled circles show mean values. Gray x’s show efficiency without targeting oligo. F) Putative recombinants were tested via PCR and phenotypically on minimal media with and without the *luxR* inducer, homoserine lactone. Purple pigment is observed with the inducer and the full length violacein operon.

Given the ease with which a 49 kb deletion was constructed, we wondered if larger deletions were possible with ORBIT. Past work has predicted many of the largest dispensable genomic segments (free of essential genes) and obtained corresponding deletion mutants (50). We designed and ordered targeting oligos that deleted one of the largest known dispensable fragments, a 134 kb region near the *galK* locus, and a predicted possible fragment of 226 kb (51). The 226 kb deletion did not yield recombinants, possibly due to the proximity to the replication terminus, which could interfere with the oligo incorporation mechanism. However, the Δ134 kb targeting oligo yielded correct recombinants that were verified by PCR, Sanger sequencing and whole genome sequencing (Fig. S7).

We next tested the ability of ORBIT to integrate large constructs onto the genome by cloning various parts of the violacein pathway (*vioA-E*) into the integrating plasmid (Fig. 4D) (26). Violacein is a purple pigment that is only produced if the entire 8.7 kb pathway (with *luxR* regulator) is present and functional and was therefore a convenient phenotypic readout. We accounted for the significant size differences of the *vio* integrating plasmids (∼2-11 kb) and transformed with a final concentration of 1 nM in 50 µL cell aliquots. We did observe decreased efficiency for large integrating plasmids, however, this decline was relatively modest – 18x lower for the longest (pInt_luxR_vioA-E_kanR, 10, 671 bp) vs. the shortest (pInt_kanR, 1, 958 bp) integrating plasmid (Fig. 4E). We tested colonies from each transformation by PCR (and nanopore sequencing) and demonstrated successful integration of the full violacein pathway as shown by production of the purple violacein pigment in the presence of the LuxR inducer (Fig. 4F). Since Bxb-1 natively acts to integrate a 50 kb circular phage genome, and plasmid integration only relies on Bxb-1, we suspect that larger integrations could be made with ORBIT.

### Multi-mutant construction with single ORBIT transformations

Strains with multiple modifications are usually constructed by introducing single modifications serially, which can become a time-consuming strategy. ORBIT is compatible with this approach, however we wanted to take advantage of our high modification efficiency to enable the simultaneous introduction of multiple modifications. First, we tried to make a double deletion mutant by adding a second targeting oligo and a second integrating plasmid (with a different selectable marker) to an ORBIT transformation. Specifically, we added 4 components to the electroporation: Δ*galK* oligo, Δ*hisA* oligo, pInt_kanR and pInt_chlorR (Fig. 5A). We performed ORBIT normally and plated for recombinants on double antibiotic plates (LB + kanamycin + chloramphenicol). We observed doubly resistant colonies that proved to be true double mutants. However, an issue with this approach is that there is no control over which integrating plasmid goes into which locus – in other words there is random assortment of the multiple integrating plasmids into the multiple attB sites introduced.

**Figure 5.**
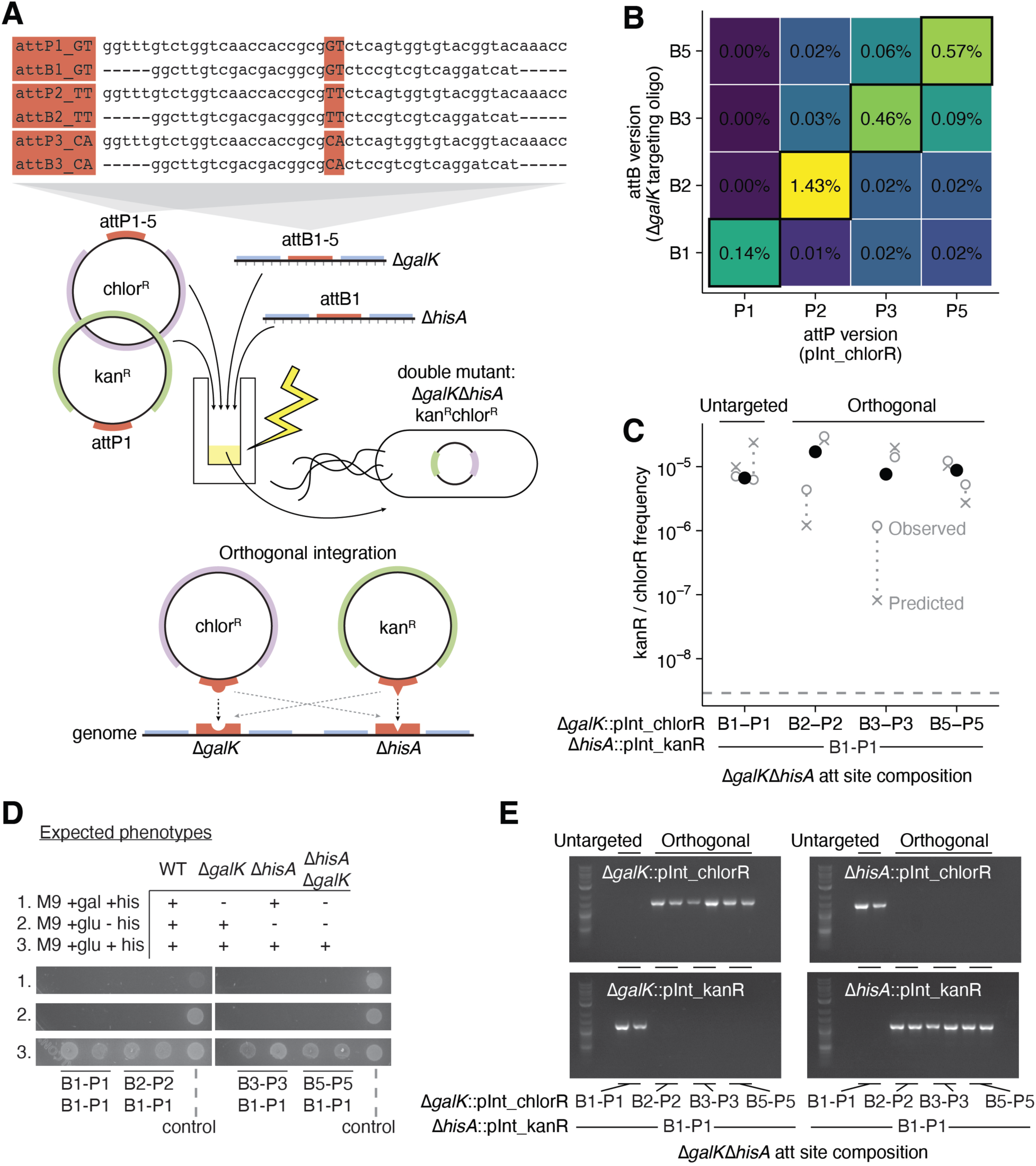
Orthogonal double mutants. A) Double ORBIT mutants are constructed by transforming both Δ*galK* and Δ*hisA* targeting oligos with both kanamycin and chloramphenicol marked integrating plasmids. Orthogonal att sites are specified by changing the central di-nucleotide where the Bxb-1 catalyzed recombination occurs (highlighted in red). These orthogonal attB and attP sites should preferentially recombine to specify integration of plasmids into defined loci, instead of randomly assorting. B) Testing single mutant efficiency Δ*galK* attB variants with pInt_chlorR attP variants. Matching att sites are outlined in black on the diagonal and the colorscale shows efficiency (n = 1 per condition). C) Double mutant frequency (kanamycin and chloramphenicol resistant colonies) for both untargeted and orthogonal double mutants with Δ*galK* (attB1, B2, B3, B5) + pInt_chlorR (attP1, P2, P3, P5) + Δ*hisA* (attB1) + pInt_kanR (attP1). Open circles show double mutant frequencies from duplicate transformations (n = 2 per condition) and filled black circles show mean values. Gray x’s show the expected double mutant frequencies calculated from the single mutant frequencies assuming independence (kanR frequency X chlorR frequency). The dashed gray line shows negative controls with no targeting oligos, but with pInt_attP1_kanR and pInt_attP1_chlorR. D) Colonies from transformations in C were phenotyped on minimal media. The expected growth pattern for single and double mutants is shown above the plate images. Control refers to a pInt only colony that is assumed to be wildtype at the test loci. E) Colonies from D were tested by PCR using primers specific to pInt_kanR or pInt_chlorR and *galK* or *hisA* to test for all four possible loci-plasmid assortments. Orthogonal transformations show the expected Δ*galK*::pInt_chlorR Δ*hisA*::pInt_kanR bands.

To overcome this random assortment issue, we generated 5 different mutant attB / attP sites on targeting oligos and pInt_chlorR plasmids, which past work suggested were mostly orthogonal and all still recognized by Bxb-1 (52) (Fig. 5A). For clarity, we now refer to the wildtype att sites as attB1 and attP1. When testing the variant targeting oligos and integrating plasmids for single locus integration efficiency, we observed many attP sites that only show high efficiency with the matching attB sites and therefore exhibit high orthogonality (Fig. 5B). Some attB/P combinations did not appear particularly orthogonal, and we did not pursue them further, but it is possible they are useful when given their preferred site in a multi mutant context. We then used orthogonal combinations of the variant pInt_chlorR plasmids (attP2, attP3, attP5), with the original kanR integrating plasmid (attP1) and the matching attB targeting oligos and observed similar efficiency as the untargeted double mutant (pInt_attP1_chlorR + pInt_attP1_kanR) (Fig. 5C). We found that nearly all tested double mutants were phenotypically correct, since they required both glucose and histidine to grow (Fig. 5D, S9). With the orthogonal transformations, we observed the specified assortment into the correct loci in all tested cases (Δ*galK*::pInt_chlorR Δ*hisA*::pInt_kanR), whereas the untargeted transformations both yielded the opposite configuration by PCR (Fig. 5E).

By plating on single and double antibiotic media, we were able to predict double mutant efficiency from observed single mutant efficiency. We found that double mutants are made at approximately the expected frequency, indicating that multi mutations should be viewed as independent events (Fig. 5C). With this expectation, we can infer the limits of this approach since the recombinant frequency must be greater than the transformed population. For our typical efficiencies of 0.1 – 1%, this indicated triple mutants should be possible (expected triple frequency 10^-9^ - 10^-6^) and we were able to make both untargeted and orthogonally targeted *ΔgalKΔhisAΔmetA_*100 bp triple mutants in a single co-electroporation, as verified by PCR (Fig. S10). Thus, our orthogonal attB/P toolkit combined with the high efficiency of ORBIT enables relatively rapid and complex multi-mutant construction strategies.

### Strategies for markerless and scarless ORBIT mutants

Frequently, it is desirable to create mutants that do not have antibiotic markers (markerless) or deletion mutants that have no trace of introduced DNA (scarless). Here, we demonstrate three different strategies for creating markerless or scarless mutants with ORBIT at each of the test loci (*ΔgalK, ΔhisA, ΔmetA, ΔleuD*) and verified the accuracy by PCR and Sanger sequencing (Fig. 6). First, all integration plasmids have flippase recognition target sites (FRT) sites that can be used in conjunction with flippase (FLP) to excise the plasmid backbone, leaving at minimum a 172 bp scar (Fig. 6A). To create these markerless strains, ORBIT mutants (without helper plasmid) were transformed with the separate FLP plasmid (pCP20 at 30°C), colonies were grown in liquid at 37°C to induce FLP expression and then plated on no antibiotic media. We observed very high efficiency of FLP excision and counter selection was not required. However, it is well known that FRT scars can cause issues when creating multi-mutants as recombination can occur between FRT scars. Consequently, we developed a clean deletion protocol that leaves no scar, by incorporating *sacB* onto the integrating plasmid (pInt_sacB_kanR) and using a second clean deletion oligo (Fig. 6B). This oligo is identical to the targeting oligo, but it does not contain an attB site. After ORBIT with pInt_sacB_kanR, helper plasmid-containing recombinants were grown and induced with m-toluic acid (oligo recombineering), transformed with the clean deletion oligo and plated on sucrose media for plasmid counter selection. Sucrose resistant mutants lost both the helper and integrating plasmids, but in addition to the clean deletion, we also observed mutants with the attB scar (Fig. S4). However, true clean deletion mutants were easily found for all four loci using colony PCR and Sanger sequencing for confirmation.

**Figure 6.**
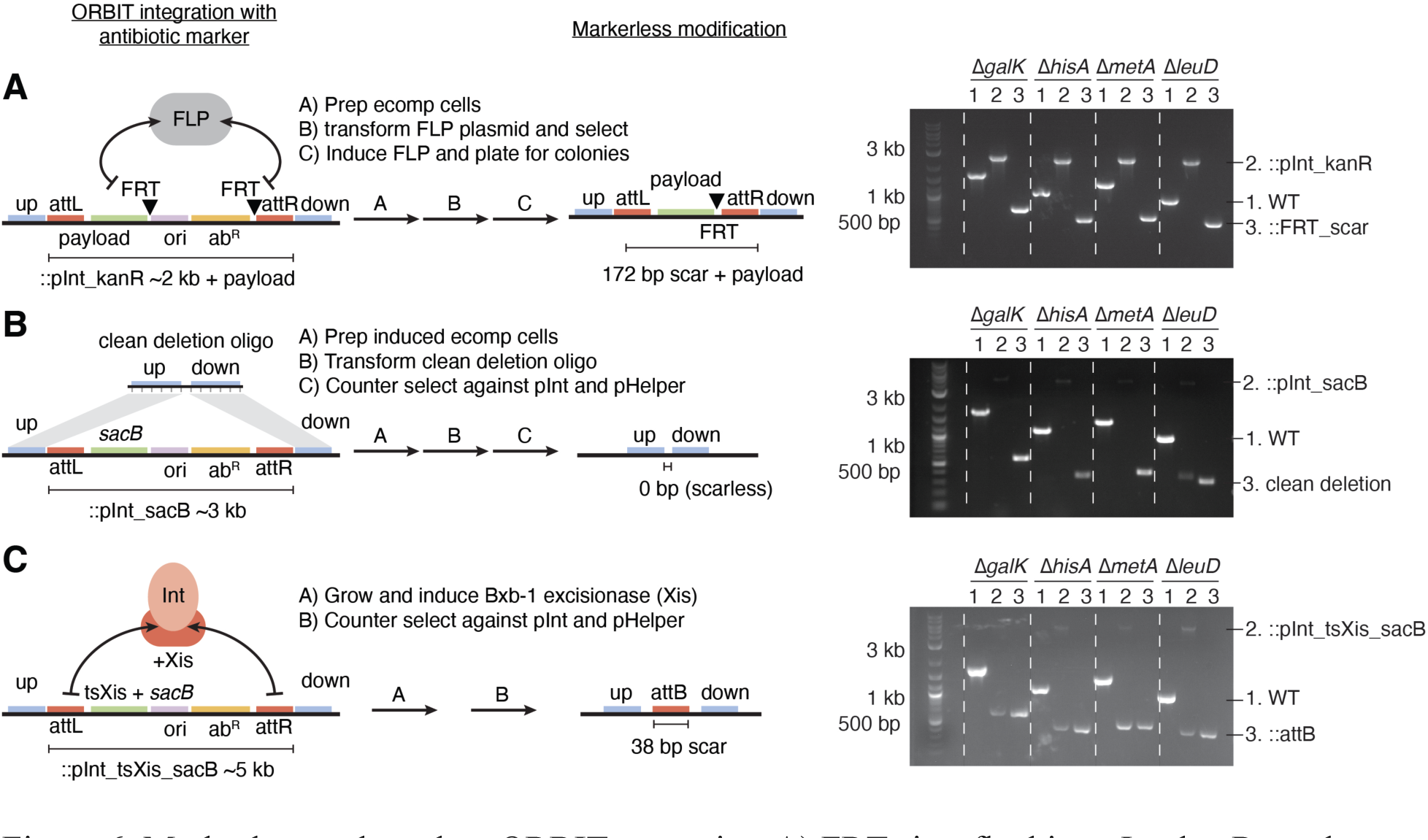
Markerless and scarless ORBIT strategies. A) FRT sites flanking pInt_kanR can be used to excise the antibiotic marker and origin, by transforming in a plasmid carrying the FLP recombinase. Representative data are shown for wildtype, Δgene::pInt_kanR and Δgene::FRT_scar loci. B) Clean deletions can be constructed by transforming with a second oligo, lacking attB, and selecting against the sacB gene on integrated pInt_sacB_kanR. Representative data are shown for the wildtype, integration and clean deletion products at four loci. C) A markerless modification with only an attB (38 bp) scar can be obtained by expressing the Bxb-1 excisionase factor (Xis) off an integrated pInt_tsXis_sacB_kanR plasmid. Excision is catalyzed by the Bxb-1 Int (from pHelper) complexed with Xis. Excision products are selected for on sucrose. Representative data are shown.

Both these strategies required prepping electrocompetent cells from recombinants, so we developed a markerless mutation strategy that was simpler to execute. Bxb-1 integrase can be turned into an excisionase by adding the directionality factor, Xis (24). We added Xis under a temperature sensitive promoter to pInt_attP1_sacB_kanR (pInt_attP1_tsXis_sacB_kanR) and used it for ORBIT at 30°C. To create the markerless mutant with only an attB scar (38 bp), we grew cells at 37°C to induce Xis (relying on leaky expression of Bxb-1 integrase) and then plated on sucrose to counter select for the excision product (and loss of helper plasmid). We anticipate that this excision-based markerless strategy may be useful for constructing multi mutants when paired with the orthogonal att sites (Fig. 5), since orthogonal attB sites may serve as relatively inert scars.

### Making mutant libraries directly from commercial targeting oligo pools

Next, we applied ORBIT for constructing pooled mutant libraries, by using many different targeting oligos in a single transformation. To estimate the mutant diversity in an ORBIT experiment, we included a random ‘N’ barcode in the Δ*galK* targeting oligo sequence (adjacent to the attB site). When used in ORBIT this oligo should yield a complex set of genomic barcodes at the defined position (Fig. 7A). We recovered ∼300k colonies from a single 50 µL transformation and then prepared an amplicon sequencing library by amplifying the pInt_kanR-genome junction spanning from the adjacent gal operon through the barcode and attR into pInt_kan. From ∼686k reads containing perfect flanking sequences, we observed ∼110, 000 unique barcode sequences. However, to be sure that barcodes were unique we performed a conservative error correction that identified ∼30k barcode sequences (Fig. 7B) (35, 36). This experiment proves that ORBIT can generate libraries with tens of thousands of distinct mutants.

**Figure 7.**
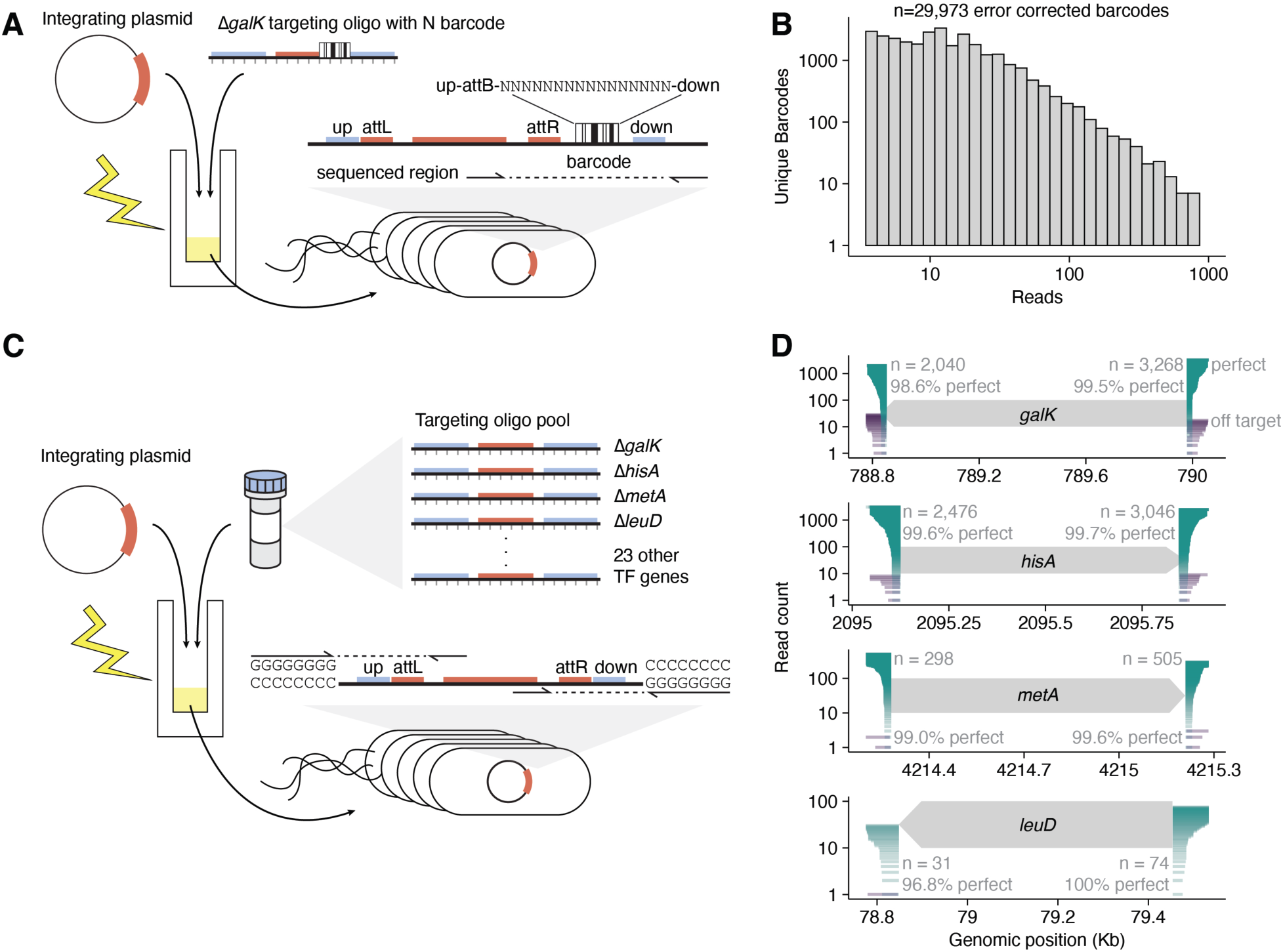
Barcoded and multi-locus mutant libraries. A) Diagram for creating a genomic barcode library at the galK locus. Barcodes are encoded as 16 N’s in a targeting oligo and the genomic region was amplified and sequenced. B) Unique barcodes observed for increasing read counts. C) Diagram for creating a 27 member mutant library of different gene deletions directly from a commercial oligo pool. Mutants were sequenced at the integrating plasmid-genome junctions by adding polyC tails to genomic fragments and amplifying with a pInt specific primer. D) Representative sequencing reads for the 4 test loci, organized by read length. Read count and the percent perfect are reported for both upstream and downstream junctions.

We then aimed to construct a more traditional mutant library that might be used in a molecular microbiology lab – a collection of 27 deletion mutants (Fig. 7C). In this set we included the 4 test genes used previously (*galK, hisA, metA, leuD*) and 23 randomly chosen transcription factor genes. This set of targeting oligos was synthesized as an IDT oPool (∼$3 / oligo), with a prep size of 50 pmol / oligo. This was enough material to transform directly into an ORBIT experiment. Therefore, we transformed 1 µM of the library directly from the IDT tube and obtained thousands of colonies from plating. To assess the accuracy of the mutant library we prepared and sequenced short read libraries of the genome-pInt_kanR junctions (upstream and downstream) using a protocol adapted from Tn-Seq (53–55). By treating the integrating plasmid sequence that replaced each gene body as an adapter and looking at where the adjacent DNA sequence mapped to the *E. coli* genome, we were able to find all the expected upstream and downstream junctions, with many “perfect” reads starting at the exact correct genomic coordinate. This indicates that every deletion mutant designed in the TO library was correctly constructed at single bp accuracy in the mutant library. Examples of this data are shown in Figure 7D for the four test genes (all loci shown in Fig. S11), and interestingly the low *ΔleuD* read counts match the relatively low individual efficiency from Fig. 3B. Given the simplicity of this mutant library construction, we expect that this workflow may be useful for small to medium scale mutant library construction.

### Creating high throughput gene knockout libraries from inexpensive oligo pools

Finally, we explored how higher throughput ORBIT mutant libraries might be constructed to understand specific features at genomic scales. As a proof of principle, we designed three targeting oligo libraries to create deletion mutations across the *E. coli* genome (Fig. 8A). The first two were focused on transcription factors (TFs) and collectively aimed to knock out every predicted DNA binding TF. Based on our earlier optimization, we decided to split the 302 TFs by gene size, since we knew that deletion length affected efficiency and we wanted to avoid extreme efficiency differences. Therefore library #1 was all TFs shorter than 550 bp (76 TF deletions 175 – 547 bp) and library #2 was all TFs longer than 550 bp (226 TF deletions 550 – 3937 bp). For library #3, we designed targeting oligos to delete all small RNA genes (90 targets 54 – 370 bp), which are small enough that they can be difficult to disrupt with transposon mutant libraries.

**Figure 8.**
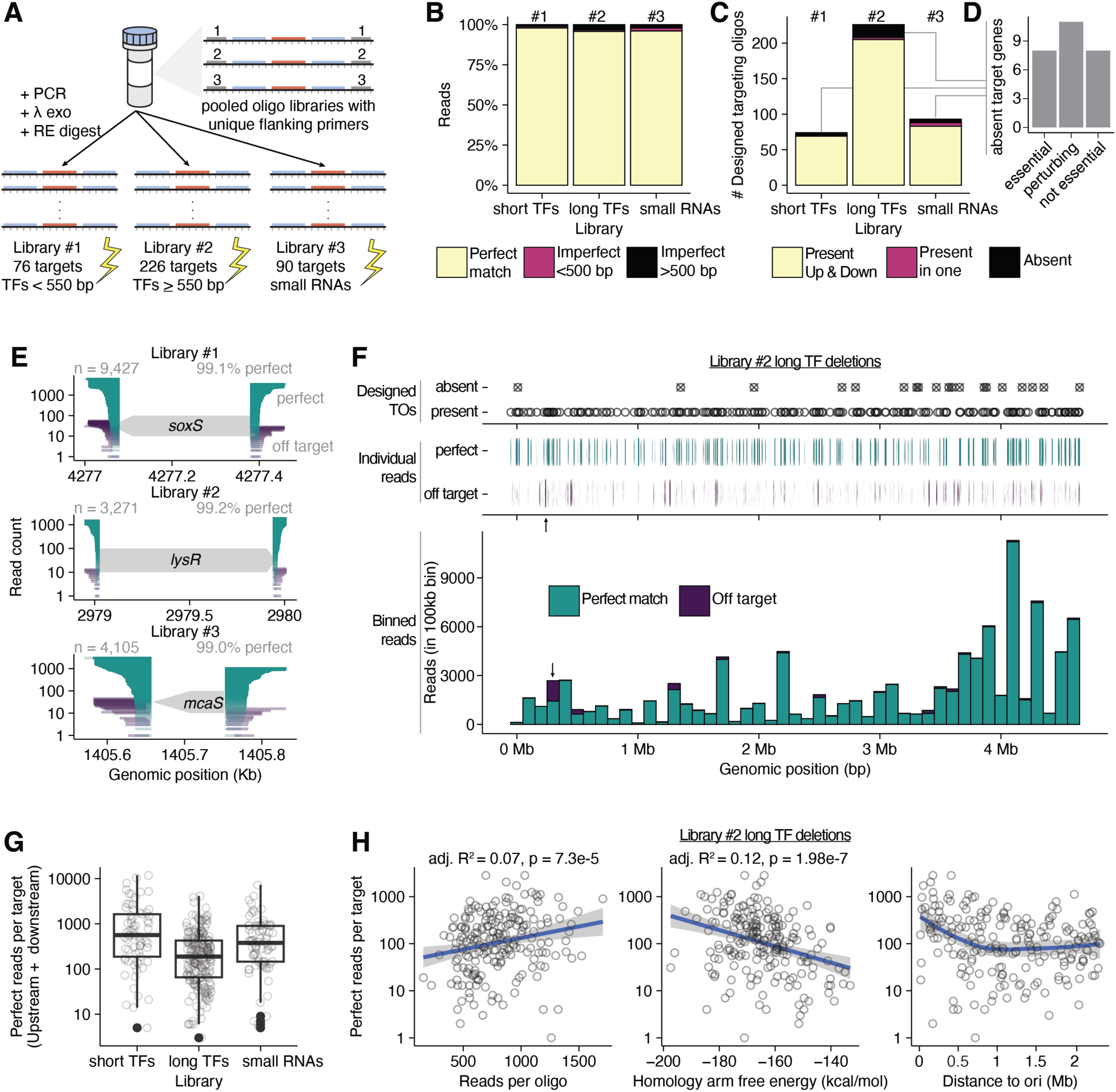
High throughput knockout libraries for transcription factor and small RNA genes. A) Diagram showing the encoding of ORBIT targeting oligos in a low abundance oligo pool. Multiple libraries are encoded in the same order by adding unique flanking primer sites. Single stranded DNA oligos are PCR amplified and then digested to get single stranded targeting oligos without flanking sites. Three different libraries were created – short TFs, long TFs and small RNAs. B) Upstream and downstream reads that passed quality controls were assessed for how well they matched target sites and >95% are perfect matches, with imperfect reads found both nearby (<500 bp) and far away (>500 bp) from target sites. C) For each set of designed targeting oligos, > 89% of targets were recovered based on the presence of perfect reads in the upstream and downstream libraries. D) For targets that were absent from the sequencing datasets, a significant fraction were annotated as essential genes or were obviously perturbing. Only 8 targets were not found at all and are not thought to have growth defects. E) Example loci from each library are shown, with perfect reads shown in green and imperfect reads shown in purple. Read counts (upstream + downstream) and percentage of perfect reads are shown. F) The genome wide distribution of perfect and off target reads is shown for Library #2, long TF deletions. The open circles show the positions of the designed targets, individual reads are shown below, and reads are binned into 100 kb regions in the bottom plot. The black arrow shows the strongest off target site in the dataset. G) Boxplots show the distribution of sequencing reads per target (upstream + downstream) for each of the three libraries. H) The variation in reads per target is compared to three variables: Reads per targeting oligo (before transformation), homology arm free energy, and the genomic distance to the origin of replication (ori). Adjusted R^2^ and p-values are shown for the linear models (blue lines) in the first two panels. The third panel shows a loess smoothing that highlights the increased reads per targets observed near the ori.

To scale up our mutant libraries to hundreds of different loci, we used low cost, low yield, multiplexed oligo pools from Twist Biosciences ($0.10 – $1.00 / oligo), which required several molecular biology steps to use as ORBIT targeting oligos. All three libraries were synthesized in a single pooled order along with other constructs for different projects. Each library had unique sets of flanking primer sites, so each targeting oligo library was amplified separately with two rounds of PCR from the original oligo pool. With the constructs now double stranded with primer sites, we needed to process them to get single stranded DNA without primer overhangs. We did this by adapting a protocol from Bonde et al. 2015, and digested away the bottom strand with lambda exonuclease (Fig. S12A)(23). Then short guide oligos were annealed to the primer sites and restriction enzymes were used to cleave the top strand at the edge of the bound guide. We found three rounds of digestion reactions were necessary to accumulate sufficient quantities of the 128 nt targeting oligo products (assessed by denaturing PAGE), and some primer overhangs remained uncleaved (Fig. S12B). Then we transformed these mixtures of correct product and primer overhang product in ORBIT experiments. A single 50 µL transformation was carried out for each library, yielding efficiencies of 0.085% (∼120k total colonies), 0.018% (21k total colonies), and 0.195% (240k total colonies) for libraries 1, 2, and 3 respectively (Fig. S12C). As expected, the longer deletions encoded in library #2 yielded lower efficiency on average than the other two libraries, however, plenty of colonies were still obtained (estimated coverage ∼92 colonies / designed mutation).

Following the same library prep, sequencing, and analysis as described above for the IDT oPool library (Fig. 7B), we assessed the accuracy of the three Twist mutant libraries. Overall, >95% of the filtered reads (i.e. high quality, adapter present, genome mapped) for each library were reads that perfectly matched a designed deletion at the exact expected nucleotide (Fig. 8B). Overall, 392 targeting oligos were designed, and 357 (90.8%) were recovered with a perfect representative in the upstream and downstream sequencing libraries (Fig. 8C). Of the few designed mutants that were not present in either the upstream or downstream sequencing libraries (n = 27), many were definitively essential genes, had known growth defects in LB, or were likely perturbing (next to essential genes) (Fig. 8D). This left only 8 mutations that were not obviously essential or perturbing that were not recovered, suggesting that ORBIT mutations failed or were below the limit of detection. Individual examples further support the high “perfect rate” of the summary statistics. Fig 8E shows example genes from each library and all target loci are shown in Figures S13-15. Perfect rates can be calculated for each target locus and median perfect rates for targets within each library are > 97%. These results indicate that ORBIT can make highly accurate mutant libraries genome wide. This is exemplified by the most complex library #2 (226 long TFs), which is shown in Fig 8F. Individual reads align to very narrow regions which correspond to on target sites and there are no obvious genomic regions where ORBIT does not work. Off target sites are rare across the genome, although library #2 does show a single particularly strong off target site. Off target sites were not conserved across the three libraries, suggesting that each targeting oligo has different off target propensities or off target integration is highly random, but further investigation of off target effects is warranted.

One potential issue from these results is that mutant abundances within the population vary dramatically, ∼1000 fold for the most and least abundant targets in each library (Fig. 8G). This can cause challenges when tracking populations in pooled assays or recovering individual low abundance mutants, therefore it would be useful to understand what drives the dramatic differences in oligo efficiency. Upstream and downstream read counts correlate (R^2^ = 0.69, p = 2.2e-16), indicating that reads per target is a reasonable proxy for relative abundance in the mutant population, and not strongly skewed by random library prep or sequencing artifacts (Fig. S16A-C). Our first hypothesis for the Twist libraries was that the original TO abundance that was transformed could strongly affect the final “perceived” efficiency of each oligo. We sequenced the original targeting oligo pools to get each TO abundance and then compared to the mutant abundances in our library. For all three libraries, there was a statistically significant correlation between initial TO abundance and mutant abundance (Fig. 8H, Fig. S17), however, we found the explanatory power to be quite weak (adjusted R^2^ = 0.04 for library #2).

We next wondered if other oligo properties could explain differences in efficiency. Past work showed that oligo recombineering efficiency was enhanced by stronger homology arm free energy (hybridization energy) (21). We calculated the minimum free energies for the homology arm hybridization energy and observed statistically significant correlations for libraries 1 & 2 (Fig. 8H, Fig. S17). The distance from the target gene to the origin of replication (*ori*) also showed a statically significant correlation for libraries 2 and 3, although it is likely this relationship is not linear, as shown for library 2 (Fig. 8H). We fit the mutant abundance data with a multiple linear regression model using oligo abundance, homology arm free energy and the distance to *ori* as independent variables. Although all three libraries were fit with statistically significant models, the explanatory power was still limited (adjusted R^2^ ≤ 0.13) (Fig. S16D), suggesting that we are still very far from having a highly predictive model based on intrinsic oligo parameters or technical parameters like oligo abundance. Therefore, targeting oligo efficiency may be explained by other biological (e.g. chromatin environment, fitness defects) or technical (e.g. read mapping, PCR efficiency) factors that remain to be explored.

## Discussion

Reverse genetics in bacteria is almost entirely dependent on generating homologous constructs through PCR or molecular cloning and is limited by recombination efficiency and throughput. Here we established ORBIT as a reverse genetic toolkit for *E. coli*, where efficient mutations are specified by DNA oligos that are transformed directly into cells. We showed that ORBIT could generate highly accurate genomic deletions of various sizes at several loci, and we demonstrated three different strategies that can be used to obtain markerless or scarless modifications. Therefore, ORBIT should be broadly useful for making common deletion mutations ranging from removal of small TF binding sites to individual genes and large operons. Our largest deletions (49 and 134 kb) also indicate that ORBIT may be useful for larger genomic modifications used in applications like genome minimization (51). To facilitate the construction of multi mutants, we made pInt derivatives with different antibiotic markers and we showed that these plasmids can be used to rapidly create double and triple mutants in a single transformation (given high efficiency). To overcome the random assortment of multiple integrating plasmids, we created orthogonal att sites, which can be used to simultaneously target multiple pInts to multiple specified loci. These orthogonal att sites may also prove useful for serially constructing markerless multi mutants with Xis, since orthogonal attB scars should be relatively inert. We anticipate that this toolkit could greatly streamline workflows where sets of multi-mutants are needed.

Beyond deletions, we established that ORBIT could insert large payloads encoded on the integrating plasmid (up to 11 kb). Large pathways or complex reporters could be integrated onto the genome at nearly any locus using ORBIT, and we created an integrating plasmid with transcriptional terminators and a cloning site flanked by FRT sites specifically for this purpose (pInt_attP1_LCS_kanR, Table S1.). Further, ORBIT may be compatible with a rapid “clonetegration” strategy, where pInt DNA assembly reactions are used directly for ORBIT without ever cloning pInt as a replicating plasmid in the pir+ host (56). An important future application is translational fusions where a fluorescent protein (or other sequence) is encoded on the integrating plasmid next to the attP site and the pInt is targeted directly upstream of the stop codon of a native gene. Murphy et al. 2018 demonstrated that constructs of this type can be used to generate translational GFP fusions in *Mycobacteria*, where the intervening attR/L scar is translated and acts as a polypeptide linker (24). Future work should test this application and the suitability of the attR/L scar as a linker in *E. coli*. Another option for smaller translational tags (e.g. his / FLAG) could be to encode the tag on the targeting oligo to avoid the linker/scar sequence issue.

One unexpected issue we observed is the integrated plasmid can be excised at detectable levels in the presence of the Bxb-1 Integrase, encoded on pHelper (Fig S3). This stands in contrast to the literature, which suggests that integration is essentially unidirectional (24, 57, 58). This phenomenon should be explored further, but in practice this issue simply requires that strains should be cured of pHelper before final verification and use in experiments. Removal of the helper plasmid is easily done by either sucrose counter selection or by mobilizing the ORBIT modification into another strain background with P1 transduction (59). Upon curing of pHelper, we established that the ORBIT modifications are highly stable. Our V1 helper plasmid also showed a high background mutation rate, which may be suboptimal for various applications. In general, we recommend using independently generated replicates and complementation to ensure background mutations are not responsible for phenotypes of interest. This issue could be circumvented by using P1 transduction to move marked ORBIT mutations into different strain backgrounds. However, we believe the best solution would be to use the V2 helper plasmid, which has similar efficiency and should accumulate few, if any, background mutations.

Probably the closest method to ORBIT is λ Red recombination, which has arguably been the standard technique for manipulating the *E. coli* genome since the early 2000s (2). Both ORBIT and λ Red rely on electroporation to introduce DNA containing homology and an antibiotic marker into the cell, however, ORBIT does not require PCR (and product purification) to generate a homology containing product and instead relies directly on the targeting oligo.

Despite this advantageous starting point, there may be reasons to use λ Red over ORBIT (e.g. to avoid flanking att scar sequence), considering it has been successfully used for over 20 years. In our hands, ORBIT is significantly more efficient than λ Red modifications at the same loci, which is consistent with the known efficiencies of single vs. double stranded DNA recombineering. λ Red is also known to be somewhat limited in the deletion size (10-200 colonies total with 5-90% accuracy for 7-82 kb)(49) and extremely limited in the integration size (< 3 kb) (4, 16, 48), whereas ORBIT easily surpassed those limitations (Fig. 3). Many alternative tools have been developed to integrate larger payloads– either designed landing pads, integrase attachment sites or transposon insertion sites, however, they typically target a single locus (5–9). More broadly, genomic integrations offer significant improvements over replicating plasmids, which require continual antibiotic selection, have single cell heterogeneity in copy number, and can result in complex genotypes due to co-transformation (60). Therefore, we believe genomic integrations should be used instead of replicating plasmids, whenever possible.

A variety of high throughput genetic methods have been published for use in *E. coli*, each with advantages and disadvantages compared to ORBIT. CRISPR-Cas methods for making genomic modifications typically involve designing a gRNA to cut the genome near a recognized PAM site and supplying a repair template that encodes the desired modification and a silent mutation that destroys the PAM site. In this manner, Cas9 supplies the initial cut and the selection against the wildtype genotype (cutting without repair is toxic) (61). At high throughput, these methods have worked well for introducing libraries of amino acid changes into specific proteins of interest without unwanted scars (11, 12), however, these techniques have not been used for kilobase scale insertion or deletion. Furthermore, these techniques rely on multipart cassettes that must be synthesized (≥ 200 nt) and cloned into replicating plasmids. There may be some applications where this intermediate in vivo cloning step may be suboptimal and unmarked mutations may be difficult to verify and track. Another technique, CRISPR-transposons may soon achieve targeted kilobase-scale insertion at high throughput, but it currently cannot achieve single nucleotide accuracy and has limited ability to direct deletions (13). Retrons are a recent extension of oligo recombineering, where ssDNA can be generated *in vivo* from a replicating plasmid inside the cell (62). These tools may become very useful in the future, but currently require extended growth periods during which the mismatch repair systems are inactivated to achieve efficient modification and are also limited to highly efficient mutations, as with classical oligo recombineering. Ultimately, each method has advantages and disadvantages, and we believe there may be interesting possibilities to explore by combining these techniques with ORBIT.

At high throughput, we established that ORBIT successfully made >350 perfect gene knockout mutants of TFs and small RNAs in three pooled transformations. To our knowledge, these precise kb-scale modifications are unprecedented at this scale. Single mutant ORBIT libraries could likely contain tens or hundreds of thousands of unique mutants, considering that we obtained ∼30, 000 uniquely barcoded strains when targeting the *galK* locus with a randomly barcoded targeting oligo. Interestingly, genomic barcoding is used to track evolving populations, and this simple ORBIT technique may be a significant improvement over existing methods (63). Further, libraries of integrating plasmids could be targeted to a single genomic site with high efficiency, providing a valuable alternative to replicating plasmid libraries and multistep landing pad methods. However, we believe the truly unique promise of ORBIT lies in targeting many different loci, genome-wide to ask new questions about how bacterial genomes function. We anticipate that the ORBIT integrating plasmid could be used as a sort of “custom transposon” for Tn-seq-like assays, where the fitness effects of pooled mutations are monitored by short read sequencing. For example, we could subject our TF or small RNA libraries (Fig. 8) to a condition of interest and read out the relative fitness of each mutant through pooled sequencing. With single nucleotide accuracy and kilobase scale modifications, this approach could be extended beyond single gene deletions in countless ways, and sophisticated logic could be executed using precisely specified controls and replicates (14). We believe these future applications will be further aided by the fact that many low throughput strategies described here may work well at high throughput (e.g. multi mutants, markerless strategies). With these new tools, there will likely also be a need to develop new ways to assay and monitor pooled libraries to answer biological questions, including sequencing strategies, fluorescence readouts (via FACS or microscopy), and biochemical readouts (via sequencing or mass spectrometry) (64).

One potential issue we encountered was the wide range of abundance between mutants in the pooled libraries, which implies different ORBIT efficiencies for each targeting oligo or another biasing factor. We anticipate that more sophisticated oligo design and a deeper understanding of recombineering efficiency could improve these outcomes. Further optimization is also required to efficiently remove the flanking primer binding sites from the targeting oligo libraries (Fig. S12). Besides ultra-cheap, low abundance oligo pools that require PCR and processing, we also demonstrated ORBIT with an oligo pool that is more expensive, but higher abundance and can be transformed directly without molecular biology (IDT oPool - Fig 7C). We believe oligo pools of this sort may be extremely useful for constructing medium throughput libraries (10s to 100s of mutants) at potentially higher frequency. With different options for targeting oligo sources, there may be new workflows that are enabled by ORBIT. For example, high throughput screens could use ultra cheap oligos, secondary screens for interesting hits might use higher abundance oligos and final hits may be constructed individually with separate oligos. Further, creating arrayed libraries with ORBIT may be an interesting avenue, since pooled mutants can be obtained and tested in 96 well plates and low abundance or missing mutants could be created separately in pools or individually. This may be a very efficient strategy compared to arraying random transposon libraries or constructing mutants one by one (3, 65).

We envision ORBIT as a single unified toolkit that can rapidly achieve genomic deletions and insertions of arbitrary size, at low and high throughput, in a way that is not possible with any other single method. Several labs across the United States have already found ORBIT useful for a variety of applications (personal communications). To facilitate uptake of these powerful tools into the research community, we have released a number of public resources, including protocols (see supplementary materials), advice (saunderslab.org/research/ec_orbit), and plasmids on Addgene (Table S1), that should be sufficient for a laboratory to start using ORBIT quickly. To simplify the targeting oligo design process, we also created a TO design app that allows users to instantly obtain targeting oligos that bind the lagging DNA strand and direct the integrating plasmid to insert in the correct orientation for gene deletions and other modifications (Fig. S1C)(saunders-lab.shinyapps.io/ ORBIT_TO_design_ecMG1655).

Beyond *E. coli* MG1655, these ORBIT tools should work in other bacteria, to varying degrees. Anecdotally, these plasmids can work in other Enterobactericae, so researchers working in closely related organisms may be able to use these tools as is. For other organisms, there are straightforward ways to try and develop an ORBIT system, with the primary limitation being an oligo recombineering system that works well, including efficient electroporation conditions.

Moving forward, ORBIT tools should enable new genetic approaches at low and high throughput that work seamlessly across bacterial strains and species and reveal robust biological insights.

## Data and material availability

Raw data are available at a GitHub repository (github.com/saunders-lab/ecoli_orbit) and the Short Read Archive (BioProject # PRJNA989253), for Illumina sequencing data. The code in the repository walks through the transformation of raw data into near final figures generated in R. Code notebooks are also publicly available as html documents hosted at saunders-lab.github.io/ecoli_orbit. Most plasmids used in this work are available on Addgene (addgene.org/browse/article/28238366). See table S1 for specific plasmid identifiers.

## Supporting information

Table S2. Oligonucleotides

Text S1. Induced electrocompetent cell protocol

Text S2. ORBIT Integration protocol

File S1. DNA maps (.gb)

## Acknowledgments

We thank Kimberly Reynolds (UTSW), Rob Phillips (Caltech), Dianne Newman (Caltech) and members of their laboratories for feedback and scientific support at various stages of this project. Specifically, we acknowledge Tom Röschinger and Manuel Razo-Mejia for their troubleshooting efforts, and Griffin Chure for his illustrator style guide. We appreciate the Center for Environmental Microbe Interactions (Caltech) for financially supporting the initial phase of this project and the Lyda Hill Department of Bioinformatics (UTSW) for supporting the Distinguished Fellow program. We would like to acknowledge the McDermott Center NGS Core (UTSW) for providing access to a Covaris instrument. We also thank Kenan Murphy, Christopher Reisch, Drew Endy, Guillaume Urtecho and the Coli Genetic Stock Center for plasmids.

## Notes

https://github.com/saunders-lab/ecoli_orbit

### Competing Interest Statement

The authors have declared no competing interest.

### Summary of Updates

Minor typos, supplemental files, SRA identifiers, and Addgene identifiers.

