## Supplemental Tables and Figures for "ORBIT for *E. coli*: Kilobase-scale oligonucleotide recombineering at high throughput and high efficiency"

**Supplemental Material:**

Table S1. *E. coli* ORBIT plasmids.
Table S2. Oligonucleotides (separate csv file).

Figure S1. Target oligo design principles and web app.
Figure S2. Commercial oligo parameters and pricing.
Figure S3. Troubleshooting attB only colonies.
Figure S4. Single gene deletion accuracy and stability.
Figure S5. Helper plasmid efficiency and off target SNP mutation rate.
Figure S6. Variable deletion size phenotypic accuracy.
Figure S7. Sanger sequence confirmation of *galK* deletions of various sizes.
Figure S8. Sanger sequence confirmation of *hisA, metA* and *leuD* deletions of various sizes.
Figure S9. Phenotypic accuracy of orthogonal and untargeted double mutants.
Figure S10. Orthogonal att sites and triple mutant construction.
Figure S11. oPool library - Upstream and downstream reads for target loci.
Figure S12. Low abundance (Twist Biosciences) targeting oligo library processing.
Figure S13. Short transcription factor library - Upstream and downstream reads for target loci.
Figure S14. Long transcription factor deletion library - Upstream and downstream reads for target loci.
Figure S15. Small RNA deletion library - Upstream and downstream reads for target loci.
Figure S16. Explaining mutant library abundances.
Figure S17. Correlations of oligo parameters and observed reads per target (perfect reads).

Text S1. Protocol for ORBIT electrocompetent cells with induced helper plasmid. (separate pdf file)
Text S2. Protocol for ORBIT integrations. (separate pdf file)

File S1. DNA maps for plasmids and sequencing libraries. (separate genbank files, compressed)

Code repository: https://github.com/saunders-lab/ecoli_orbit

| Name | Purpose | Inducer | Resistance | sacB counterselection | Host strain | Identifier |
|---|---|---|---|---|---|---|
| **Helper plasmids** | | | | | | |
| pHelper_Ec1_V1_gentR | Replicating ORBIT V1 helper plasmid w/ inducible oligo recombineering module and bxb-1 integrase | m-toluic acid / arabinose | Gentamicin | Yes | DH5α | Addgene #205291 |
| pHelper_Ec1_V1_ampR | Replicating ORBIT V1 helper plasmid w/ inducible oligo recombineering module and bxb-1 integrase | m-toluic acid / arabinose | Ampicillin / Carbenicillin | Yes | DH5α | Addgene #205292 |
| pHelper_NoMutL_V2_gentR | Replicating ORBIT V2 helper plasmid. No mutL gene for low background mutation rate | m-toluic acid / arabinose | Gentamicin | Yes | DH5α | Addgene #205293 |
| **Integrating plasmids** | | | | | | |
| pInt_attP1_kanR | Nonreplicating ORBIT integrating plasmid with WT attP (P1) sequence. Replicates in pir+ strains for cloning. | NA | Kanamycin | No | Pir+ | Addgene #205294 |
| pInt_attP1_chlorR | Nonreplicating ORBIT integrating plasmid with WT attP (P1) sequence. Replicates in pir+ strains for cloning. | NA | Chloramphenicol | No | Pir+ | Addgene #205295 |
| pInt_attP1_ampR | Nonreplicating ORBIT integrating plasmid with WT attP (P1) sequence. Replicates in pir+ strains for cloning. | NA | Ampicillin / Carbenicillin | No | Pir+ | Addgene #205296 |
| pInt_attP2_kanR | Orthogonal version of pInt_attP1_kanR with attP2 sequence. | NA | Kanamycin | No | Pir+ | Addgene #205297 |
| pInt_attP2_chlorR* | Orthogonal version of pInt_attP1_chlorR with attP2 sequence. | NA | Chloramphenicol | No | Pir+ | NA |
| pInt_attP3_chlorR | Orthogonal version of pInt_attP1_chlorR with attP3 sequence. | NA | Chloramphenicol | No | Pir+ | Addgene #205298 |
| pInt_attP5_chlorR* | Orthogonal version of pInt_attP1_chlorR with attP5 sequence. | NA | Chloramphenicol | No | Pir+ | NA |
| pInt_attP5_ampR | Orthogonal version of pInt_attP1_ampR with attP5 sequence. | NA | Ampicillin / Carbenicillin | No | Pir+ | Addgene #205299 |
| pInt_attP1_LCS_kanR | pInt_attP1_kanR with Library cloning site (LCS) flanked by transcriptional terminators for reporters / constructs. FRT sites excise backbone, not payload. | NA | Kanamycin | No | Pir+ | Addgene #205300 |
| pInt_attP1_sacB_kanR | pInt_attP1_kanR with the sacB gene (sucrose sensitivity), for clean deletions or marker excision. | NA | Kanamycin | Yes | Pir+ | Addgene #205301 |
| pInt_attP1_tsXis_sacB_kanR | pInt_attP1_kanR with the sacB gene (sucrose sensitivity) and Bxb-1 excisionase (Xis) under temperature sensitive regulation. For markerless modifications (leaves attB scar). | High temperature (37-42°C) | Kanamycin | Yes | Pir+ | Addgene #205302 |
| **Other** | | | | | | |
| BW25141* | Pir+ host for cloning / replicating ORBIT integrating plasmids | NA | None | No | Pir+ | CGSC #7635 |
| pCP20* | Temperature sensitive plasmid with temperature sensitive FLP induction. For marker excision between FRT sites. | High temperature (37-42°C) | Ampicillin / Choramphenicol | No (Temp Sensitive) | DH5α | CGSC #7629 |

Table S1. *E. coli* ORBIT plasmids. Green rows indicate the standard plasmids used throughout the paper, unless otherwise specified. Blue rows note integrating plasmids with orthogonal attP sites. Yellow rows show integrating plasmids that are specifically designed to facilitate markerless or scarless mutations. All plasmids, except those indicated with an asterisk are available on Addgene.

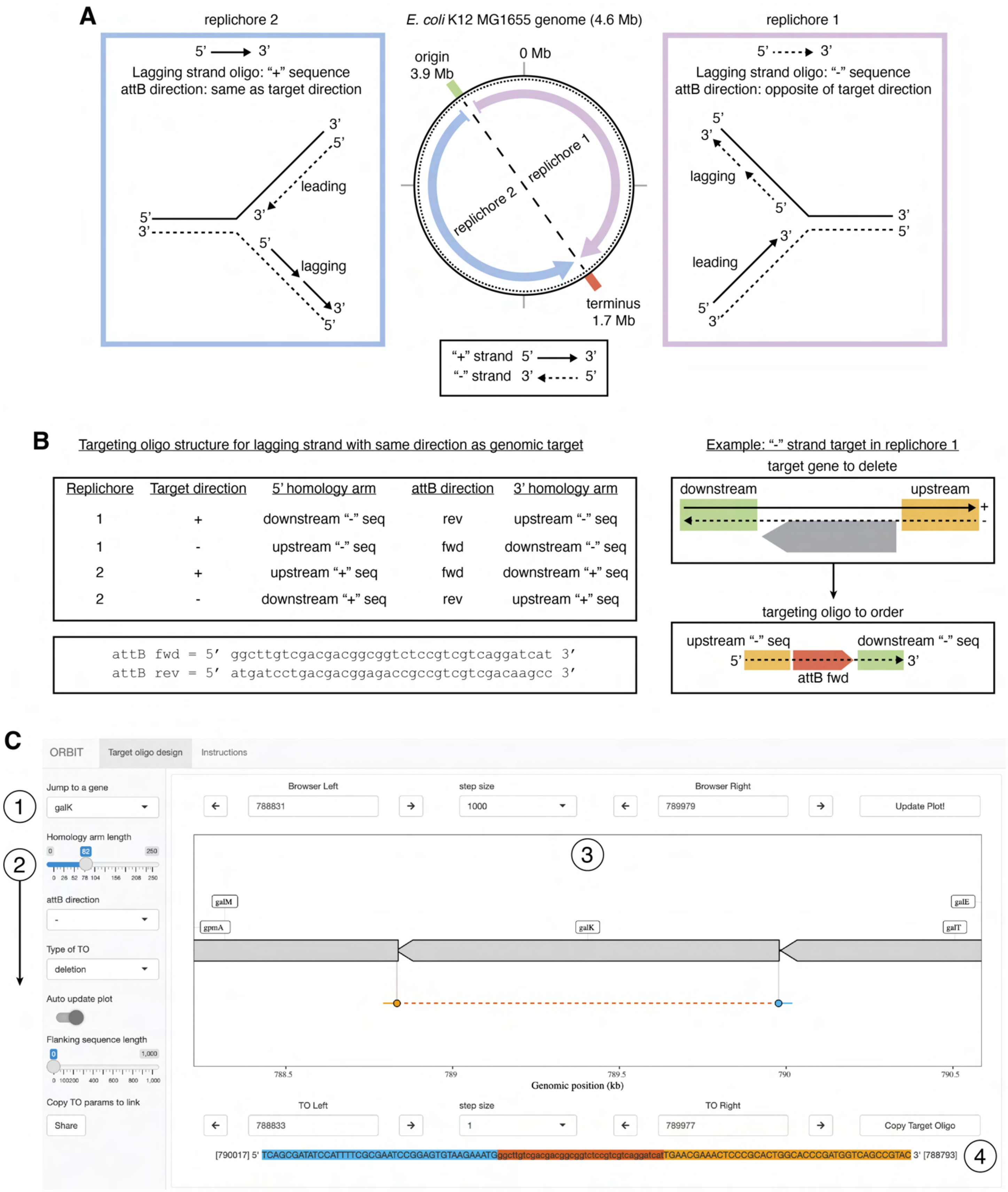


Figure S1. Target oligo design principles and web app. A) The *E. coli* K12 MG1655 genome is split into two replichores starting at the origin and ending at the terminus. The origin occurs at 3.9 Mb, not 0 Mb. The forward or "+" genomic strand is shown as a solid line and the reverse or "-" genomic strand is shown as a dashed line. For the genomic sites in replichore 1, the replication fork will advance as shown with the newly synthesized lagging strand encoded as the “-” genomic sequence. The replication fork proceeds in the opposite manner for replichore 2, therefore the lagging strand is encoded as the “+” genomic sequence. B) To design a targeting oligo that is incorporated into the lagging strand and inserts the ORBIT pInt_attP plasmid into the same orientation as a genomic target, both the replichore and target direction need to be accounted for. Oligonucleotides are typically specified and ordered as 5’ to 3’ sequences, and the table provides all four possible targeting oligo structures in this format. An example of a “-” strand gene on replichore 1 is shown with its corresponding ORBIT targeting oligo structure. C) A web app for targeting oligo design simplifies the design process for targets in the *E. coli* K12 MG1655 genome. (1) Users can jump to a gene of interest and immediately obtain a targeting with parameters specified in the sidebar (2), including homology arm length, attB direction (typically same direction as gene) and type of TO (deletion, N terminal, C terminal or insertion). The main genome browser window is shown in (3) and users can click on genes, use arrows or type in genomic coordinates to move the plot window and targeting oligo positions. (4) The resulting targeting oligo is instantly displayed, which can be copied and ordered directly from commercial sources. Flanking genomic sequence can also be displayed to facilitate colony PCR primer design. URL: saunders-lab.shinyapps.io/ORBIT_TO_design_ecMG1655

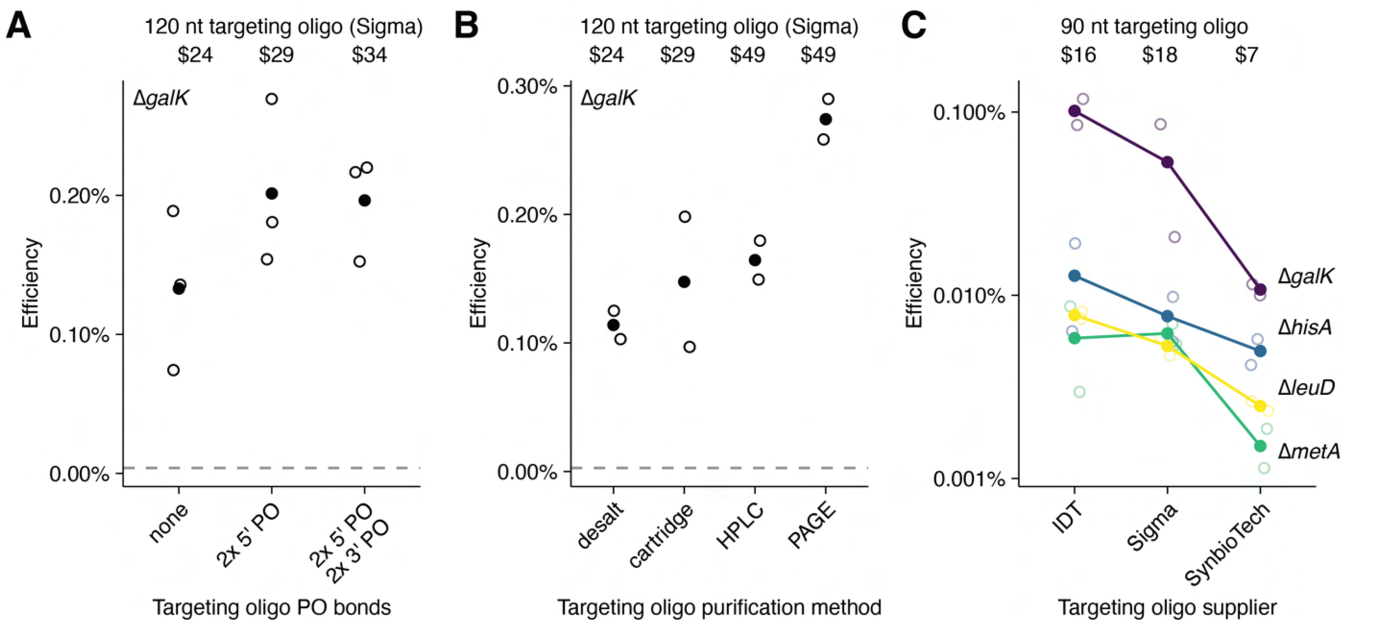


Figure S2. Commercial oligo parameters and pricing. A) The efficiency of targeting oligos (Δ*galK*) with phosphorothioate bonds (2x) at the 5' end or both the 5' and 3' ends (n = 3). B) The efficiency of ΔgalK targeting oligos with different purification options from Millipore Sigma (n = 2). C) The efficiency of targeting oligos from three different suppliers (n = 2). All prices reflect the current academic pricing offered to UT Southwestern Medical Center in 2023. Open circles show individual transformations and filled circles show mean values.

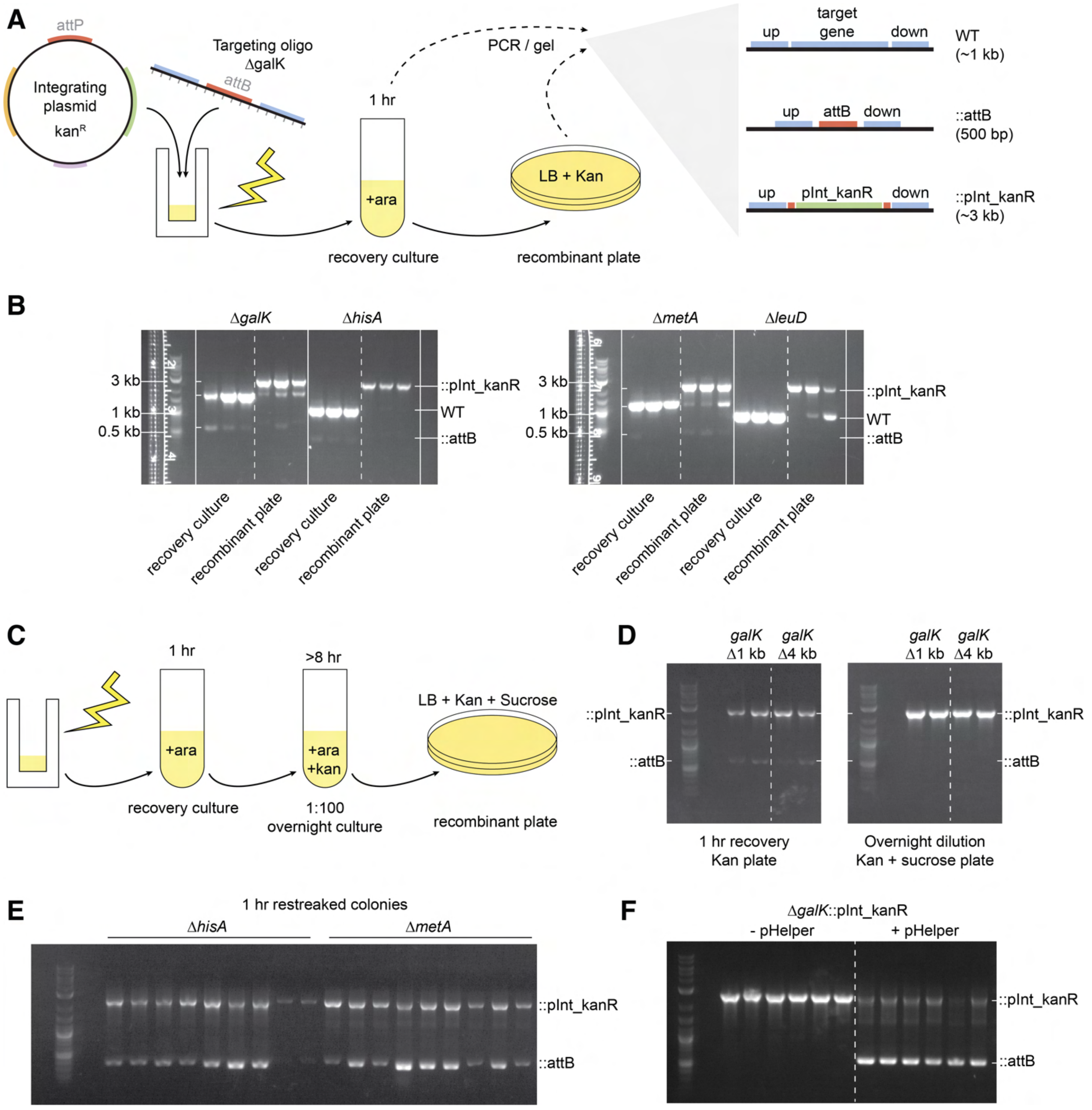


Figure S3. Troubleshooting attB only colonies. A) Assessing the three possible locus structures during the recovery and plating of an ORBIT experiment. PCRs spanning the loci were performed and wildtype (WT), Δgene::pInt_kanR, and Δgene::attB were differentiated by size. B) Gels show attB is visible during the recovery culture, while pInt_kanR is not, however upon plating Δgene::pInt_kanR becomes the dominant band at all 4 loci. C) Diagram showing the workflow for 1 hour recovery cultures, which typically retain pHelper, and an overnight protocol for obtaining recombinants without pHelper. D) Comparing 1 kb and and 4 kb deletions at the galK locus using the 1 hr and overnight protocols. The attB band disappears for the overnight colonies, leaving only the Δ*galK*::pInt_kanR band. E) Restreaking Δ*galK* colonies from 1 hour recovered experiments that show attB bands resulted in many colonies that still had attB and pInt_kanR bands. F) pHelper was retransformed into a Δ*galK*::pInt_kanR strain, which had previously been cured of pHelper. Spanning PCRs were performed colonies from both strains, and the attB is only visible when pHelper was added back.

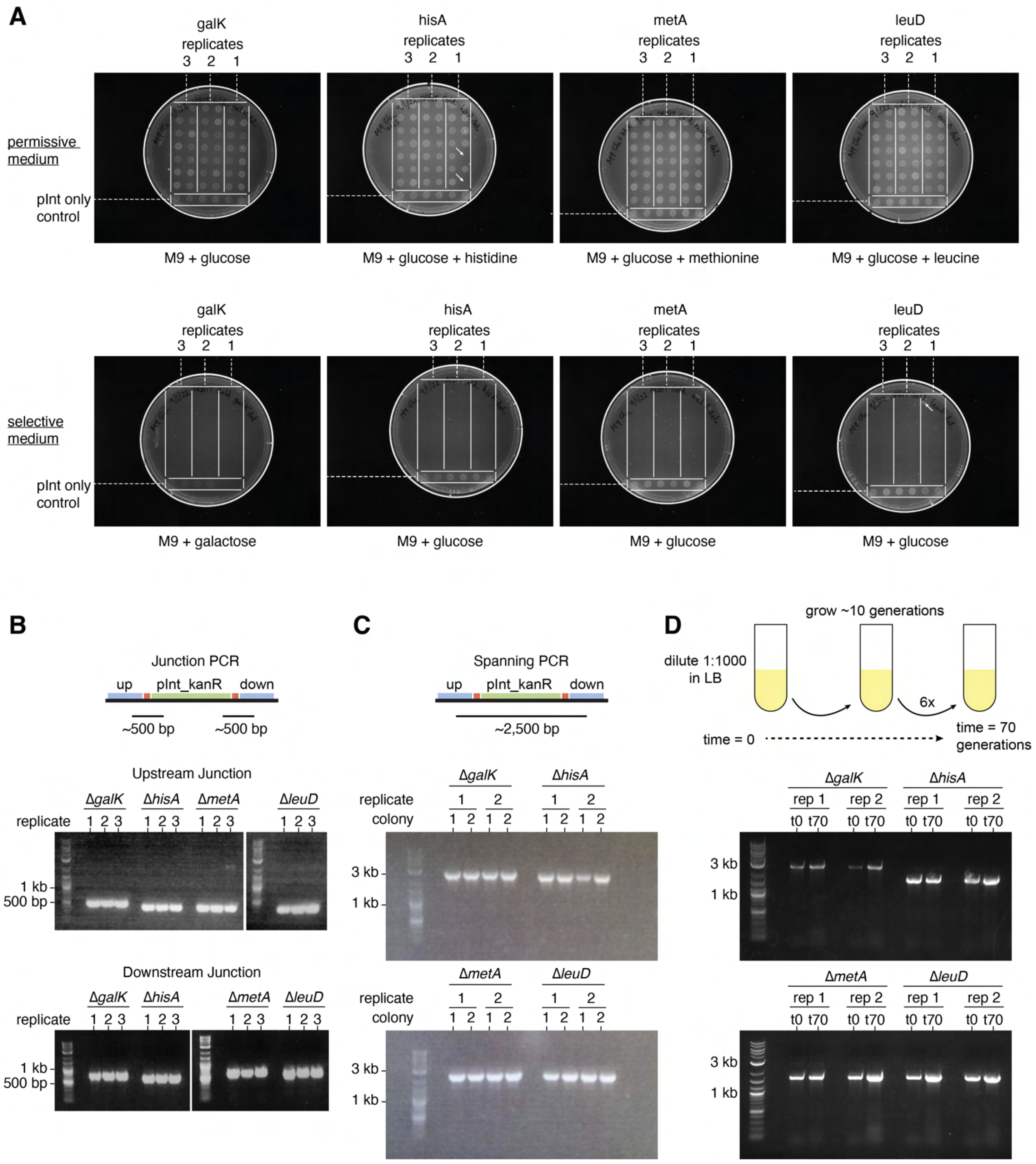


Figure S4. Single gene deletion accuracy and stability. A) Phenotypes on selective and permissive minimal media for gene knockouts (n=14 colonies for 3 replicate transformations.), including negative (pInt only) controls (n = 6) that are assumed to be wildtype at the targeted loci. B) PCR of pInt_kanR-genome junctions (n=3 colonies) and C) PCR spanning the entire locus (n = 4 colonies from two transformations). D) Stability test of Δgene::pInt_kanR helper cured strains over 70 generations in the absence of antibiotic selection. Tested by PCR spanning the loci (n = 2 passaged cultures per locus).

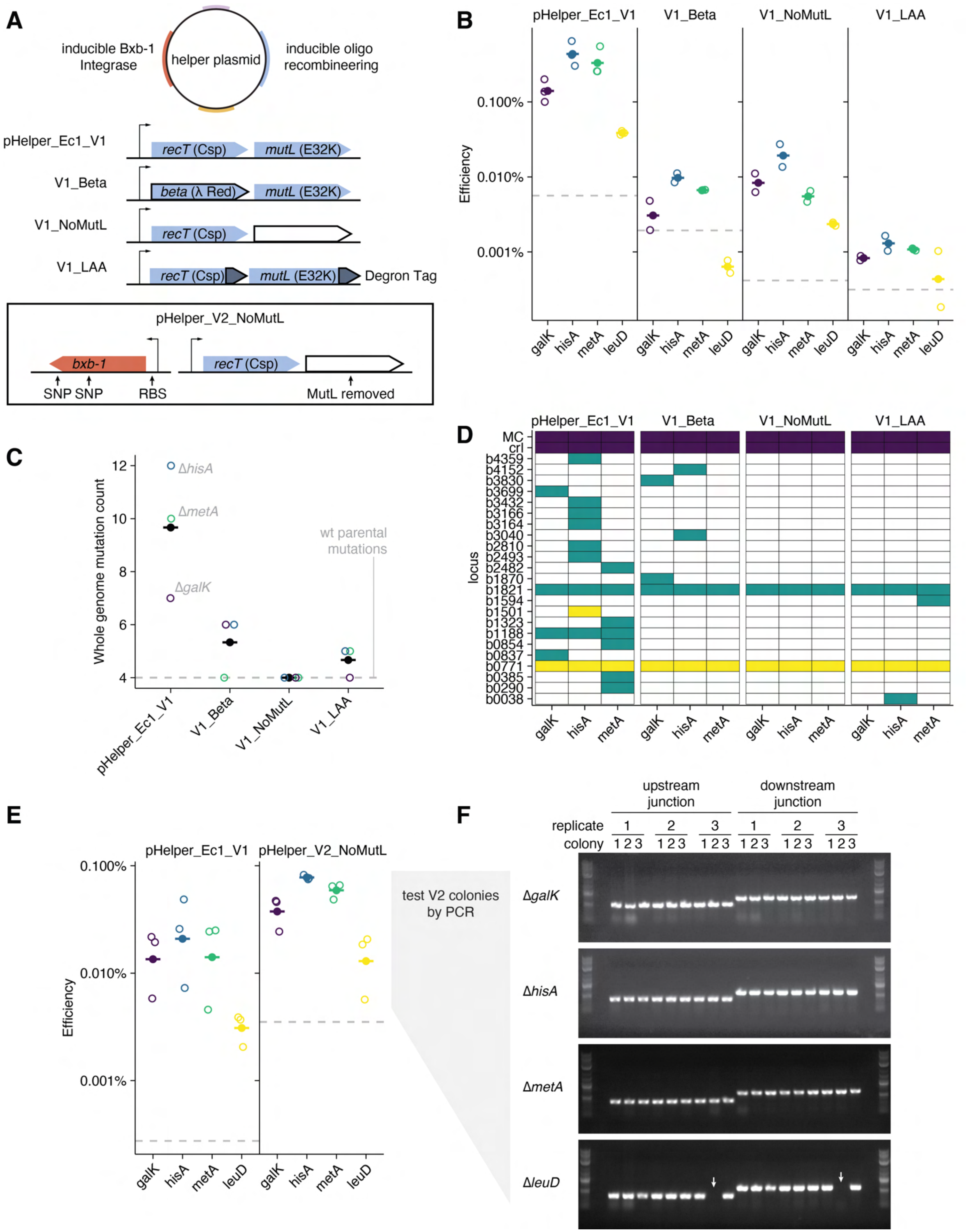

A
inducible Bxb-1 Integrase
helper plasmid
inducible oligo recombineering
pHelper_Ec1_V1
recT (Csp)
mutL (E32K)
V1_Beta
beta (λ Red)
mutL (E32K)
V1_NoMutL
recT (Csp)
V1_LAA
recT (Csp)
mutL (E32K)
Degron Tag
pHelper_V2_NoMutL
bxb-1
recT (Csp)
SNP SNP
RBS
MutL removed
B
pHelper_Ec1_V1
V1_Beta
V1_NoMutL
V1_LAA
Efficiency
0.100%
0.010%
0.001%
galK hisA metA leuD
C
Whole genome mutation count
12
10
8
6
4
ΔhisA
ΔmetA
ΔgalK
wt parental mutations
pHelper_Ec1_V1
V1_Beta
V1_NoMutL
V1_LAA
D
pHelper_Ec1_V1
V1_Beta
V1_NoMutL
V1_LAA
locus
MC
crl
b4359
b4152
b3830
b3699
b3432
b3166
b3164
b3040
b2810
b2493
b2482
b1870
b1821
b1594
b1501
b1323
b1188
b0854
b0837
b0771
b0385
b0290
b0038
galK hisA metA
E
pHelper_Ec1_V1
pHelper_V2_NoMutL
Efficiency
0.100%
0.010%
0.001%
galK hisA metA leuD
test V2 colonies by PCR
F
upstream junction
downstream junction
replicate 1 2 3 1 2 3
colony 1 2 3 1 2 3 1 2 3 1 2 3 1 2 3 1 2 3
ΔgalK
ΔhisA
ΔmetA
ΔleuD

Figure S5. Helper plasmid efficiency and off target SNP mutation rate. A) Designs for four different helper plasmids in addition to the original pHelper_Ec1_V1_gentR plasmid. The modifications were: V1_Beta had the λ-Red *beta* gene instead of *cspRecT,* V1_NoMutL had the *mutL_E32K* gene removed, and V1_LAA had degron tags added to both *cspRecT* and *mutL_E32K.* pHelper_noMutL_V2 was made from V1_NoMutL and also included two fixed mutations in *bxb-1* and added a more optimal ribosome binding site. B) ORBIT deletion efficiency for each of the four V1 helper plasmids at each of the four test loci (pHelper_Ec1_V1 is same data as fig 3). Open circles represent individual transformations and filled circles show mean values (n = 2 or 3 per condition). Gray dashed lines show negative controls without targeting oligo. C) For each V1 plasmid, three recombinants (1x ΔgalK, 1x ΔhisA, and 1x ΔmetA) made with each helper plasmid were sent for whole genome sequencing. In addition to the correct ORBIT target mutation, single nucleotide polymorphisms (SNPs) were detected randomly throughout the genomes. The number of those SNPs for each mutant is reported in open circles, and filled points show the mean number of mutations found from each set of helper plasmid created mutants. The gray line shows the number of SNPs detected in the parental WT strain, and are therefore unrelated to ORBIT. D) The specific distribution and type of mutations across loci is shown, where blue refers to deletion, green refers to transitions and yellow refers to transversions. E) Deletion efficiency for the original V1 compared to the NoMutL V2 helper plasmid (circles and dashed lines are same as B). F) Using pHelper_NoMutL_V2, three replicate ORBIT experiments were performed for each locus and three colonies from each experiment were tested by junction PCR. White arrow shows a *leuD* colony that is missing a band, and is putatively incorrect.

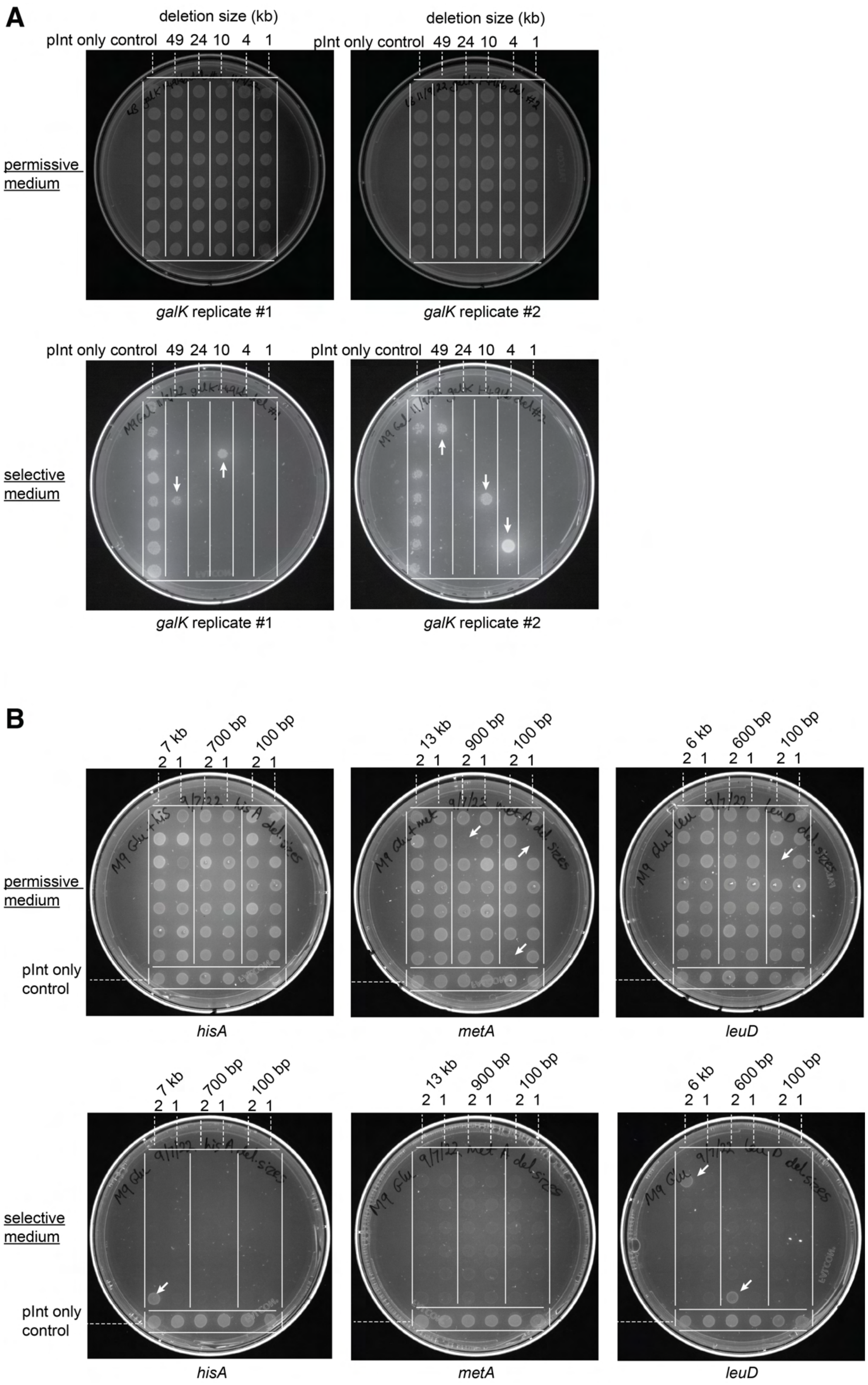
A
deletion size (kb)
pInt only control 49 24 10 4 1
permissive medium
galK replicate #1
galK replicate #2
selective medium
B
7 kb 700 bp 100 bp
2 1 2 1 2 1
13 kb 900 bp 100 bp
6 kb 600 bp 100 bp
pInt only control
hisA
metA
leuD

Figure S6. Variable deletion size phenotypic accuracy. A) Deletion mutants of various sizes at the galK locus (1 – 49 kb) were tested for phenotypic accuracy on permissive (LB) and selective plates (M9 + galactose) (n=8 colonies for 2 transformations per deletion). LB was used, because biotin biosynthesis was predicted to be disrupted in the 49 kb mutant. Negative controls (pInt only) were included and assumed to be wildtype at the galK locus (n=8). B) Deletions mutants of various sizes at the hisA, metA and leuD loci were tested for phenotypic accuracy on permissive (M9 glucose + histidine, methionine, or leucine) or selective plates (M9 glucose) (n = 7 colonies for 2 transformations per deletion). Negative controls (pInt only) were included and assumed to be wildtype at the tested loci (n=6). Putatively incorrect colonies are highlighted with white arrows.

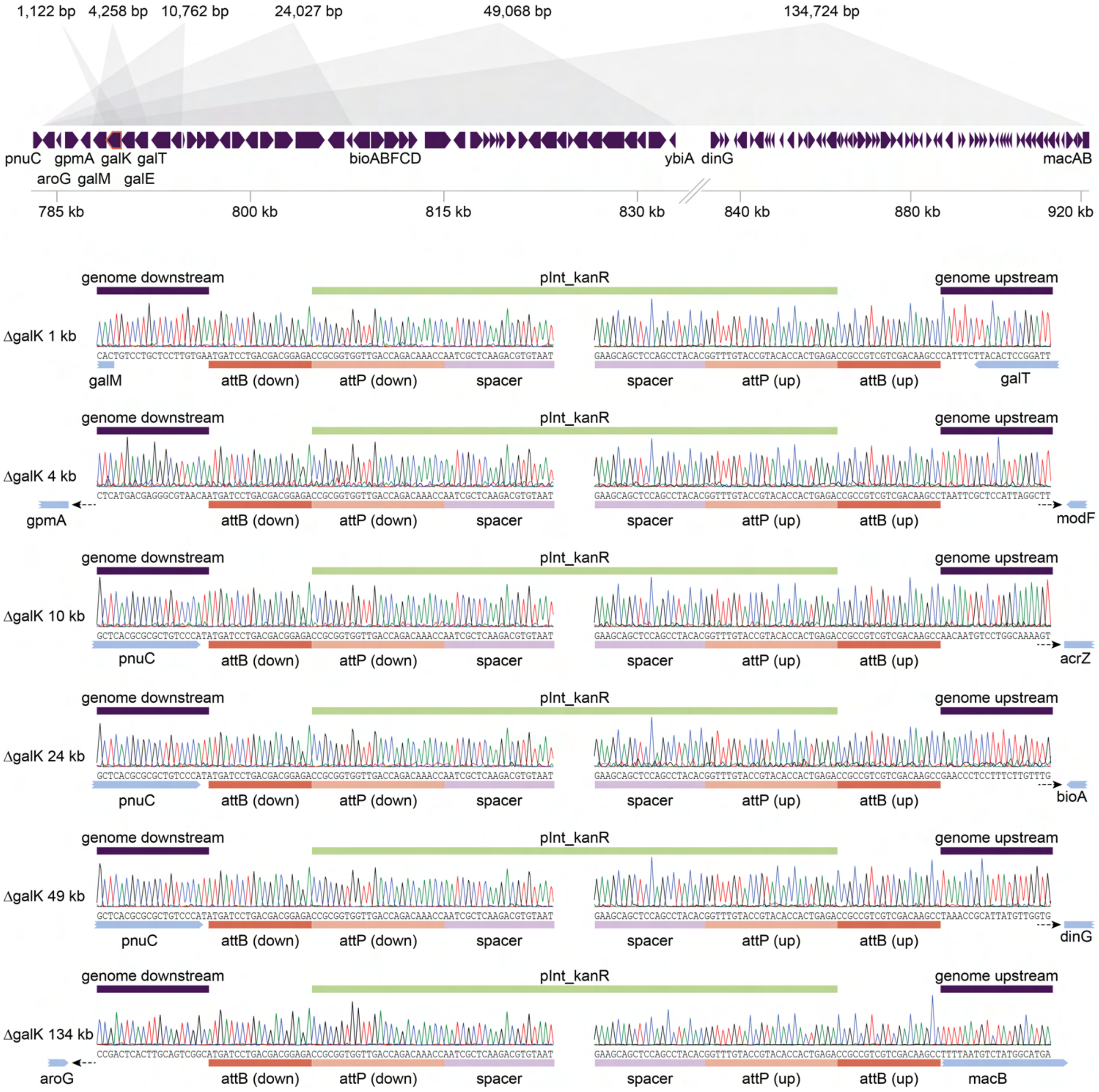


Figure S7. Sanger sequence confirmation of *galK* deletions of various sizes. The gene diagram shows the designed deletions of different sizes surrounding the *galK* (highlighted red) locus. Beyond the 1-49 kb deletions tested in Fig 4, a 134 kb deletion that was designed later is also shown. Due to the difference in scales, a break is included in the axis for genomic position. PCRs spanning the entire loci were performed and sent for sanger sequencing from either end to cover the genome – pInt_kanR junctions. Representative sanger traces show the expected sequences.

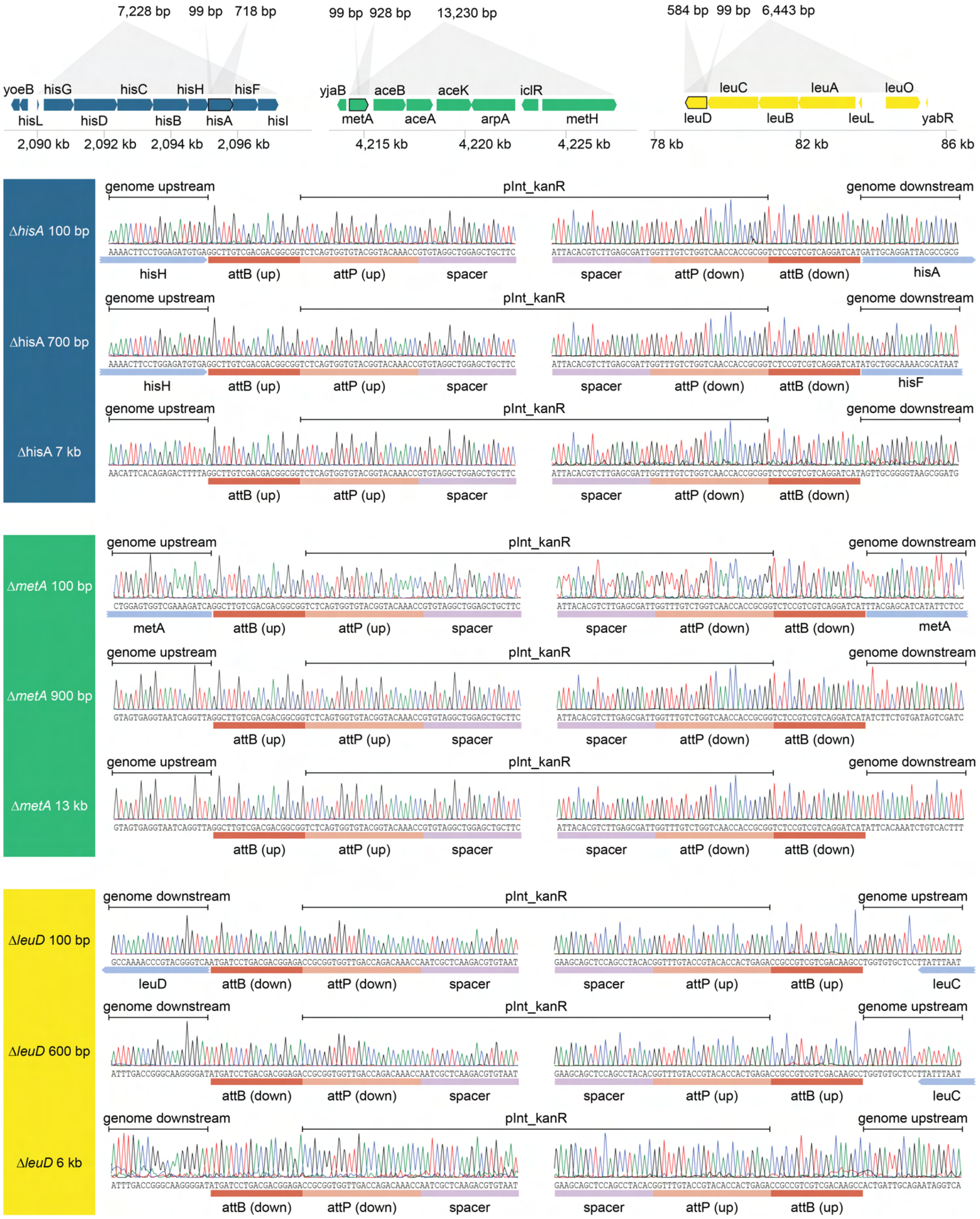


Figure S8. Sanger sequence confirmation of *hisA, metA* and *leuD* deletions of various sizes. The gene diagrams shows the designed deletions of different sizes surrounding the target loci (genes from Fig 3 are outlined in black). PCRs spanning the entire loci were performed and sent for sanger sequencing from either end to cover the genome – pInt_kanR junctions. Representative sanger traces show the expected sequences.

**A**

| | Expected phenotypes | | | |
|---|---|---|---|---|
| | off target | ΔhisA | ΔgalK | ΔhisAΔgalK |
| LB | + | + | + | + |
| M9 glucose | + | - | + | - |
| M9 glucose + histidine | + | + | + | + |
| M9 galactose | + | - | - | - |
| M9 galactose + histidine | + | + | - | - |

**B**

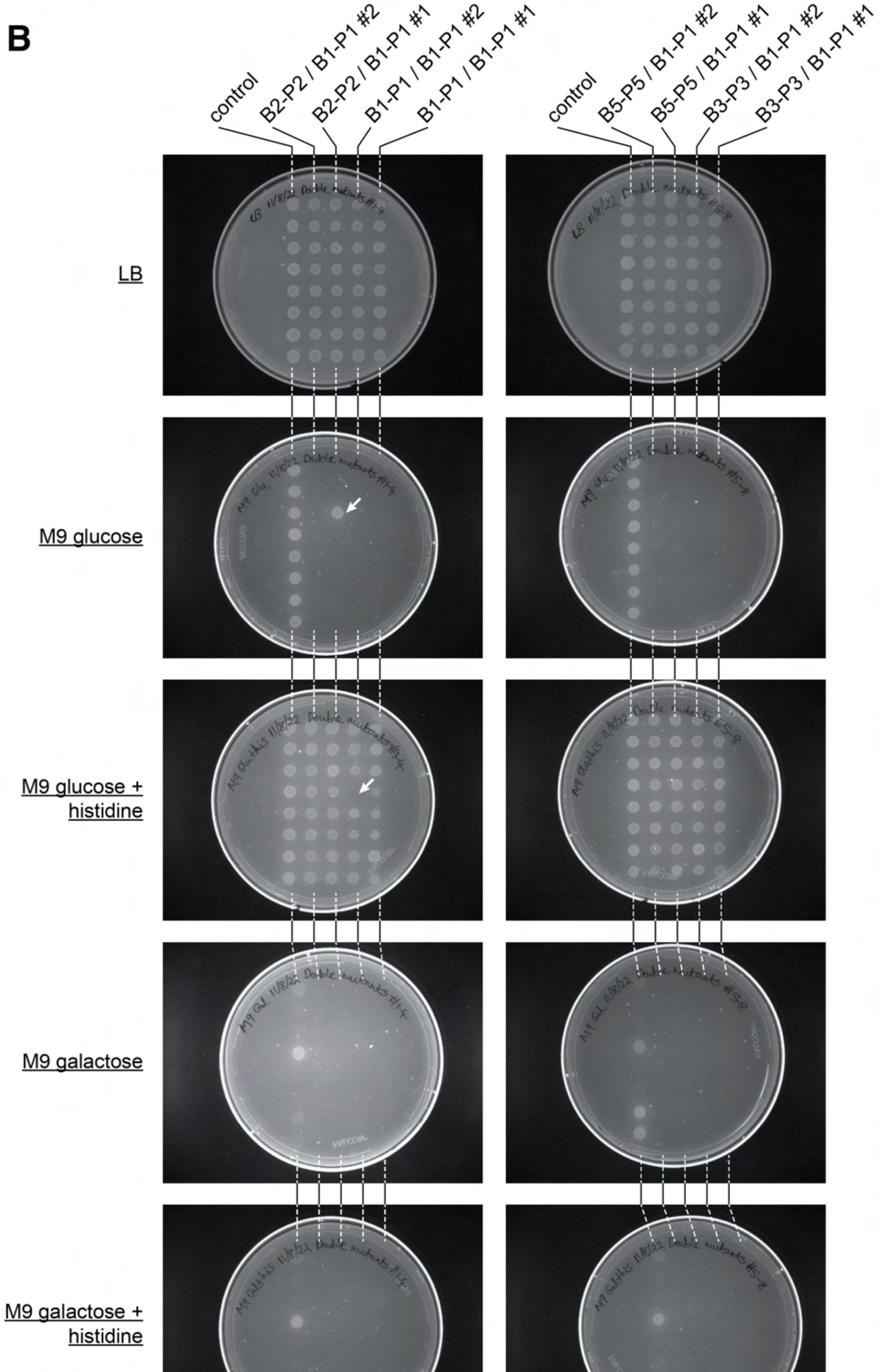


Figure S9. Phenotypic accuracy of orthogonal and untargeted double mutants. A) The table shows the expected growth phenotypes for wildtype, single, and double mutants (Δ*hisA*Δ*galK*) on various minimal media. True double mutants were expected to only grow on LB and M9 + glucose + histidine. B) Colonies were phenotyped on all 5 media and scored for growth (n = 8 for 2 transformations per double mutant). Negative controls (pInt only) were included and assumed to be wildtype at the tested loci (n=8). Putatively incorrect colonies are highlighted with white arrows.

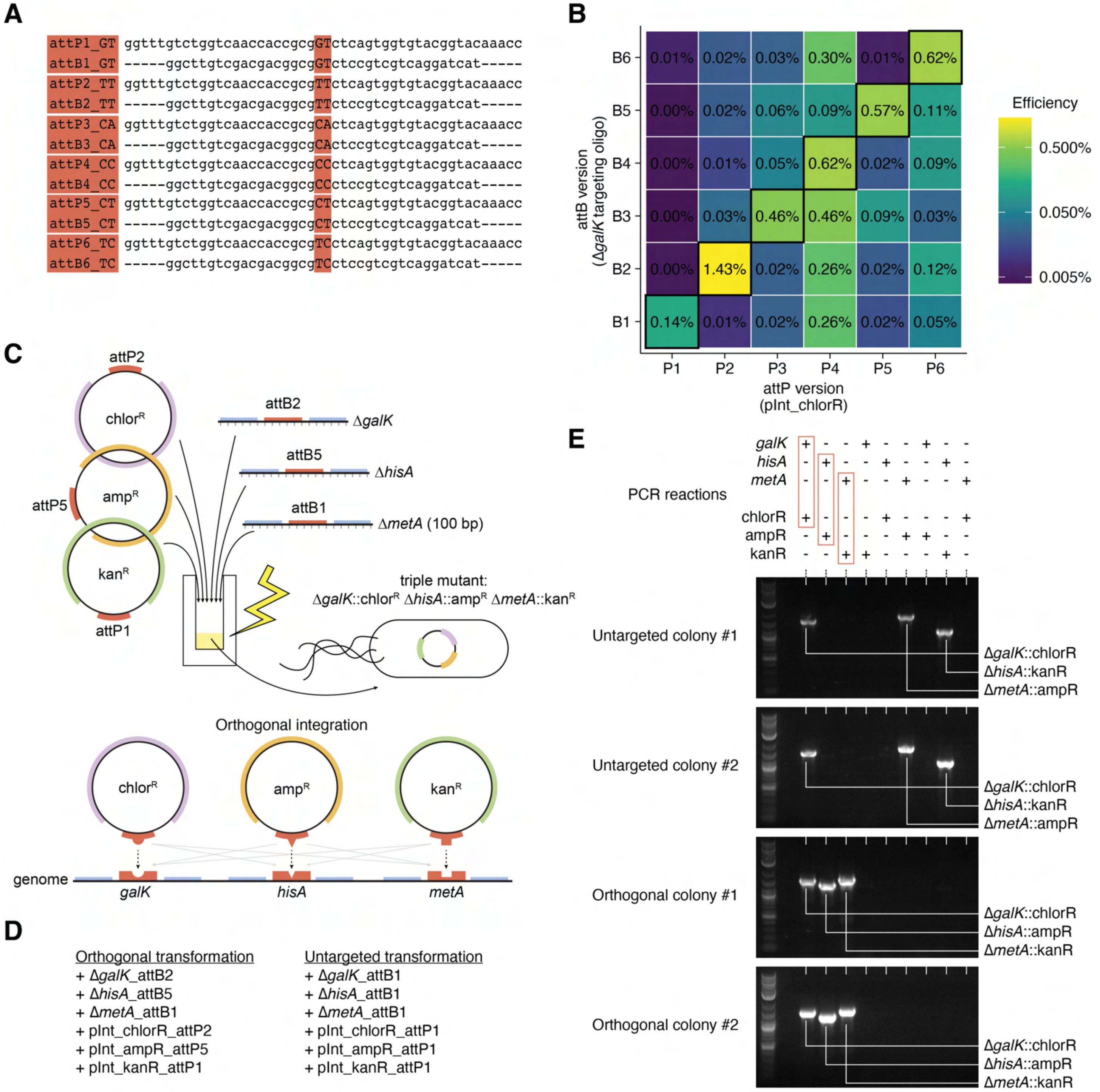


Figure S10. Orthogonal att sites and triple mutant construction. A) Wildtype (attB1 / attP1) and mutant att sites are shown, which vary at the central di-nucleotide. B) Same as Fig. 5B, but for the complete set of attB/P1-6 sites. Single mutant efficiencies were tested at the galK locus with the variant attB sites and the variant attP sites (on pInt_chlorR) (n=1 per attB/attP pair). The matching attB/attP pairs are shown on the diagonal and outlined in black. The color scale shows efficiency and the numbers in each tile show the exact efficiency value. C) Orthgononal triple mutants were contronstructed by adding three targeting oligos (ΔgalK_attB2, ΔhisA_attB5 and ΔmetA_attB1) and three integrating plasmids with different resistance markers (pInt_attP2_chlorR, pInt_attP5_ampR, and pInt_attP1_kanR). Bxb-1 integration should occur between matching orthogonal sites to give the specified integration pattern: ΔgalK::pInt_chlorR ΔhisA::pInt_ampR ΔmetA::pInt_kanR, although off target integration could occur in any combination. D) Orthogonal and untargeted transformations were performed. The untargeted triple mutant is the same three targeting oligos and integrating plasmids, but with the wildtype att sites (attB1/attP1) for all components. E) PCRs were performed for each possible target locus – pInt pair. The expected orthogonal genotype is highlighted in red. Both orthogonal transformation colonies (n=2) show the expected result, while both untargeted transformation colonies (n=2) show a different integration pattern (ΔgalK::pInt_chlorR ΔhisA::pInt_kanR ΔmetA::pInt_ampR) than orthogonal, but the same as each other.

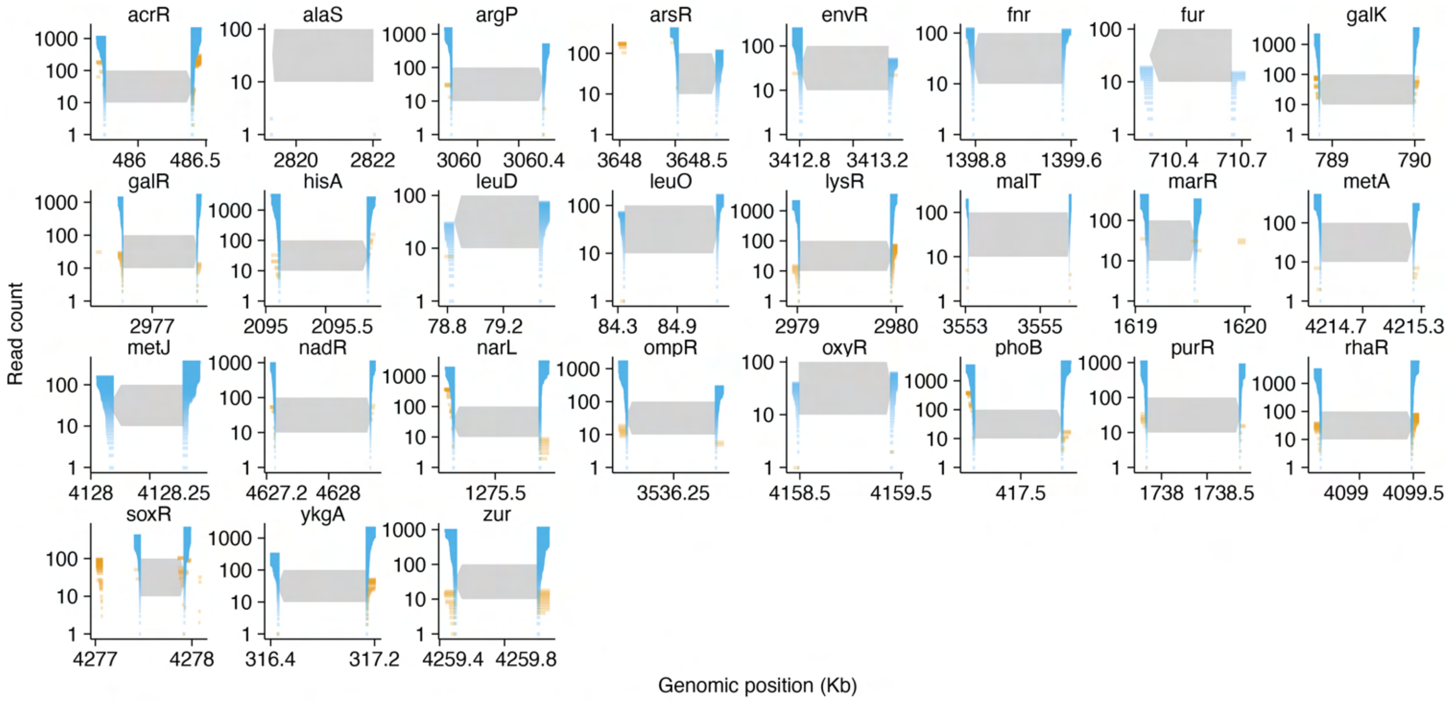


Figure S11. oPool library - Upstream and downstream reads for target loci. Plots show all reads mapping within 500 bp of either end of the target genes. Orange segments show imperfect reads and blue segments show perfectly matched reads. The number of reads in each category, upstream – perfect, upstream – imperfect, downstream – perfect, downstream – imperfect are shown on the y-axis and reads are ordered by read length. Genomic coordinates are shown on the x-axis.

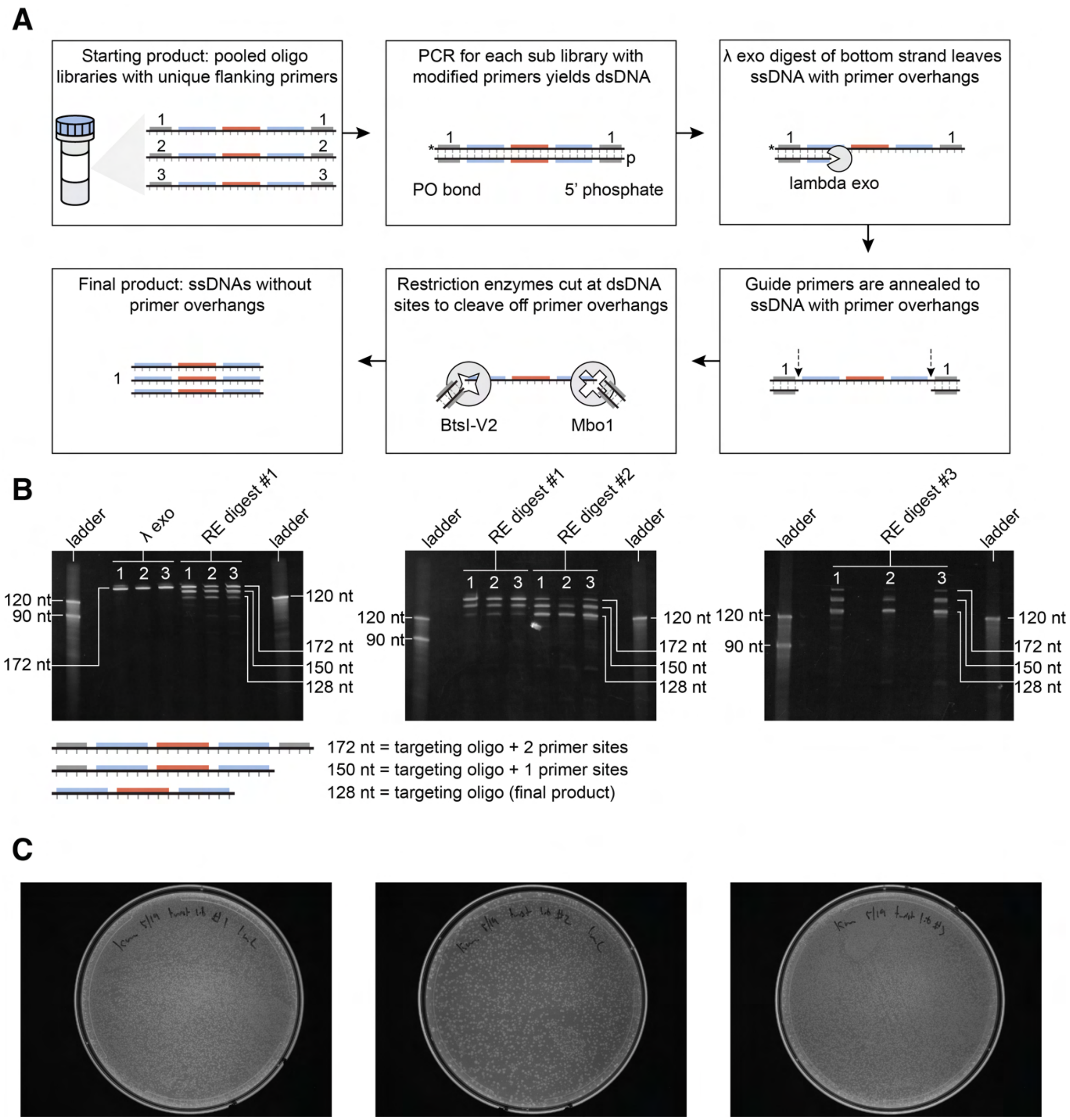


Figure S12. Low abundance (Twist Biosciences) targeting oligo library processing. A) A low abundance oligo pool was ordered encoding all three mutant libraries with unique flanking primer sites. PCR was performed with modified primers, resulting in a double stranded DNA product. The primer modifications promoted the digestion of only the bottom strand by lambda exonuclease to return the libraries into single stranded form. Then guide primers binding to the flanking primer sites were annealed and restriction enzymes were added that cleaved only at the double stranded sites on the flanking primer sequences. This resulted in a final single stranded product without PCR overhangs. This protocol was adapted from Bonde et al. 2015. B) PAGE gels show three products following exonuclease and restriction enzyme treatment – targeting oligo + both primer sites, targeting oligo + 1 primer site and the final targeting oligo alone. These are assumed to represent incomplete digestion products. The initial digest was repeated twice for a total of three digests, which mostly final targeting oligo product. C) Transformation of these libraries in ORBIT experiments yielded colonies as shown by the plate images.

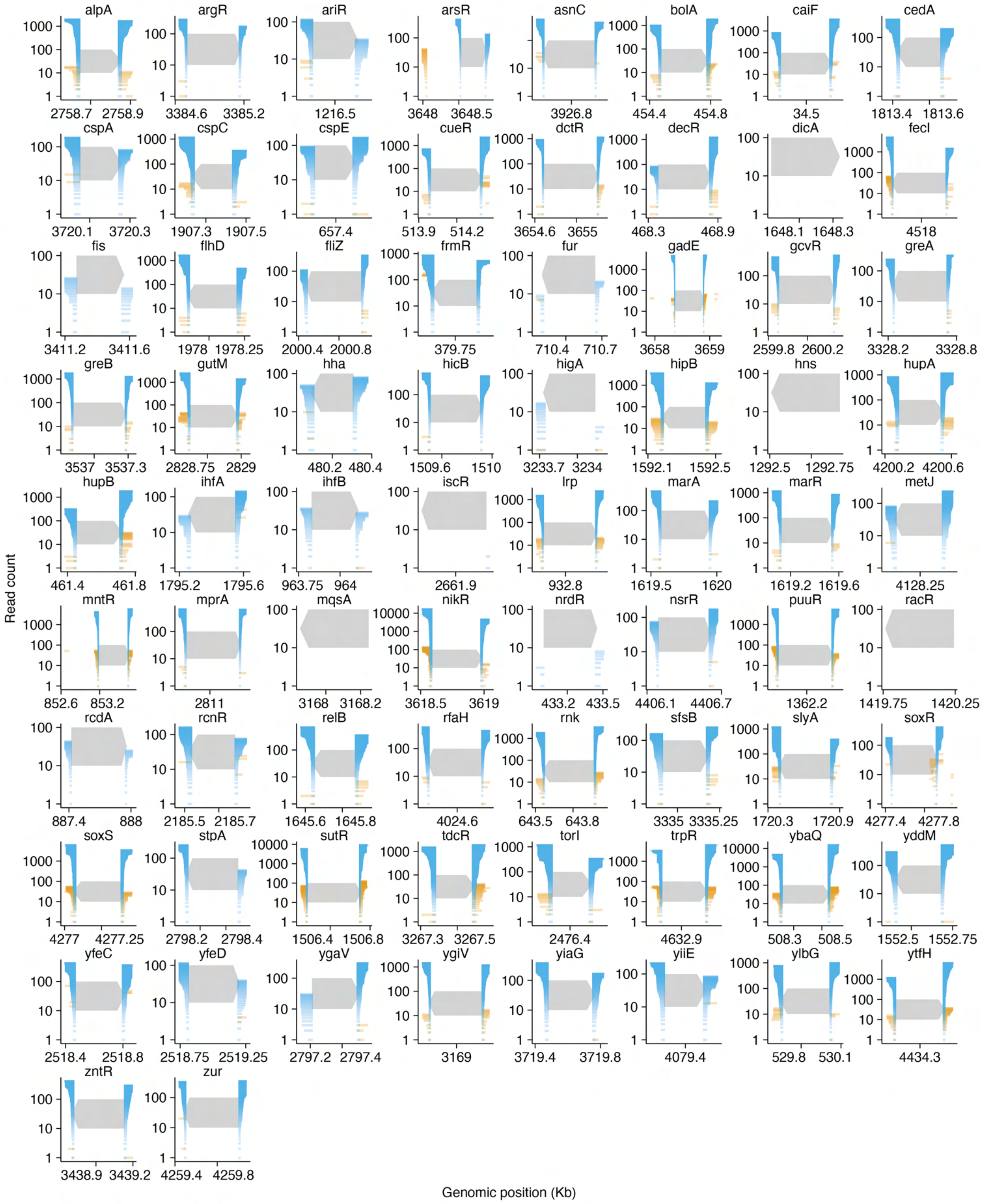


Figure S13. Short transcription factor library - Upstream and downstream reads for target loci. Plots show all reads mapping within 500 bp of either end of the target genes. Orange segments show imperfect reads and blue segments show perfectly matched reads. The number of reads in each category, upstream – perfect, upstream –

imperfect, downstream – perfect, downstream – imperfect are shown on the y-axis and reads are ordered by read length. Genomic coordinates are shown on the x-axis.

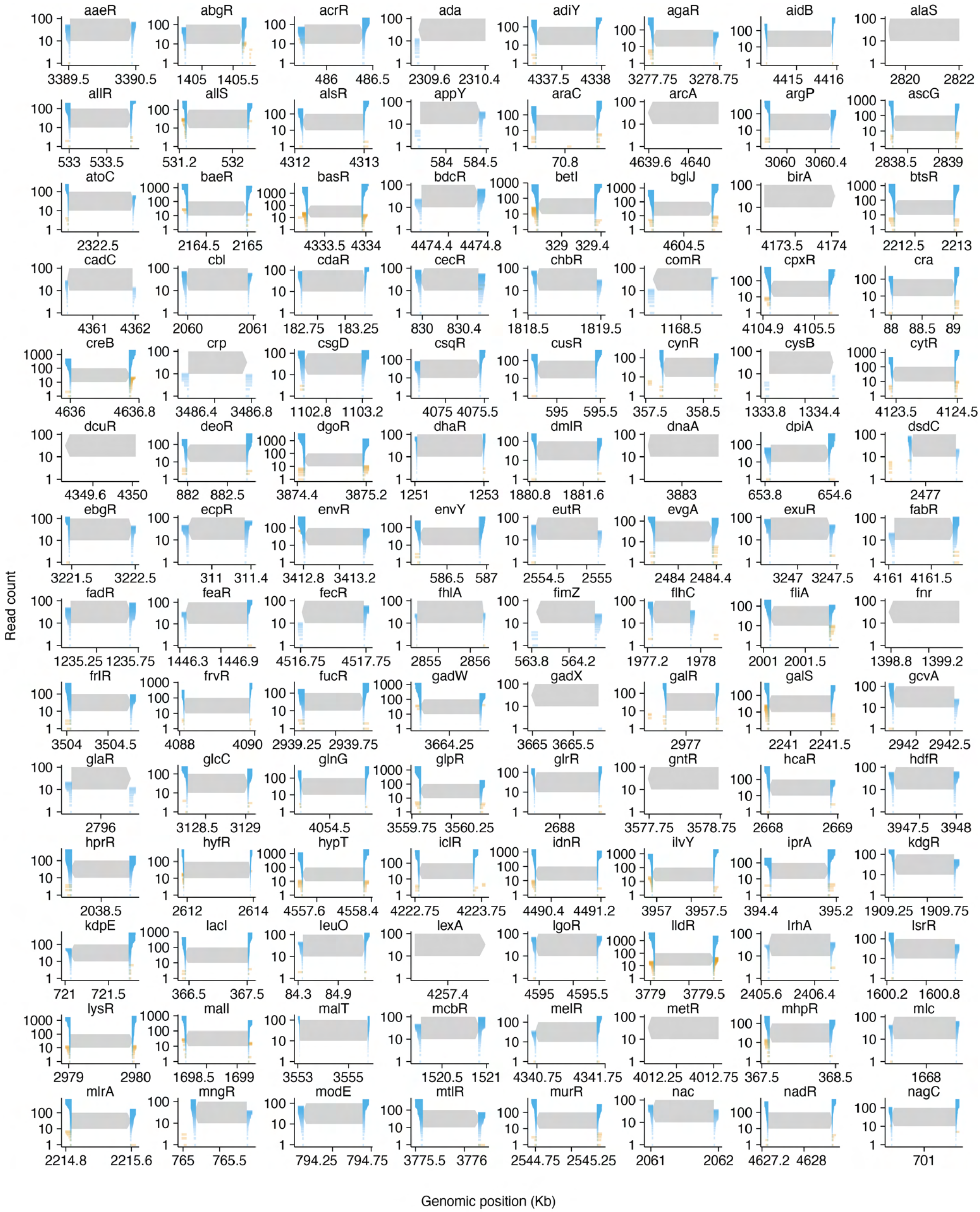


Figure S14a. Long transcription factor deletion library - Upstream and downstream reads for target loci.

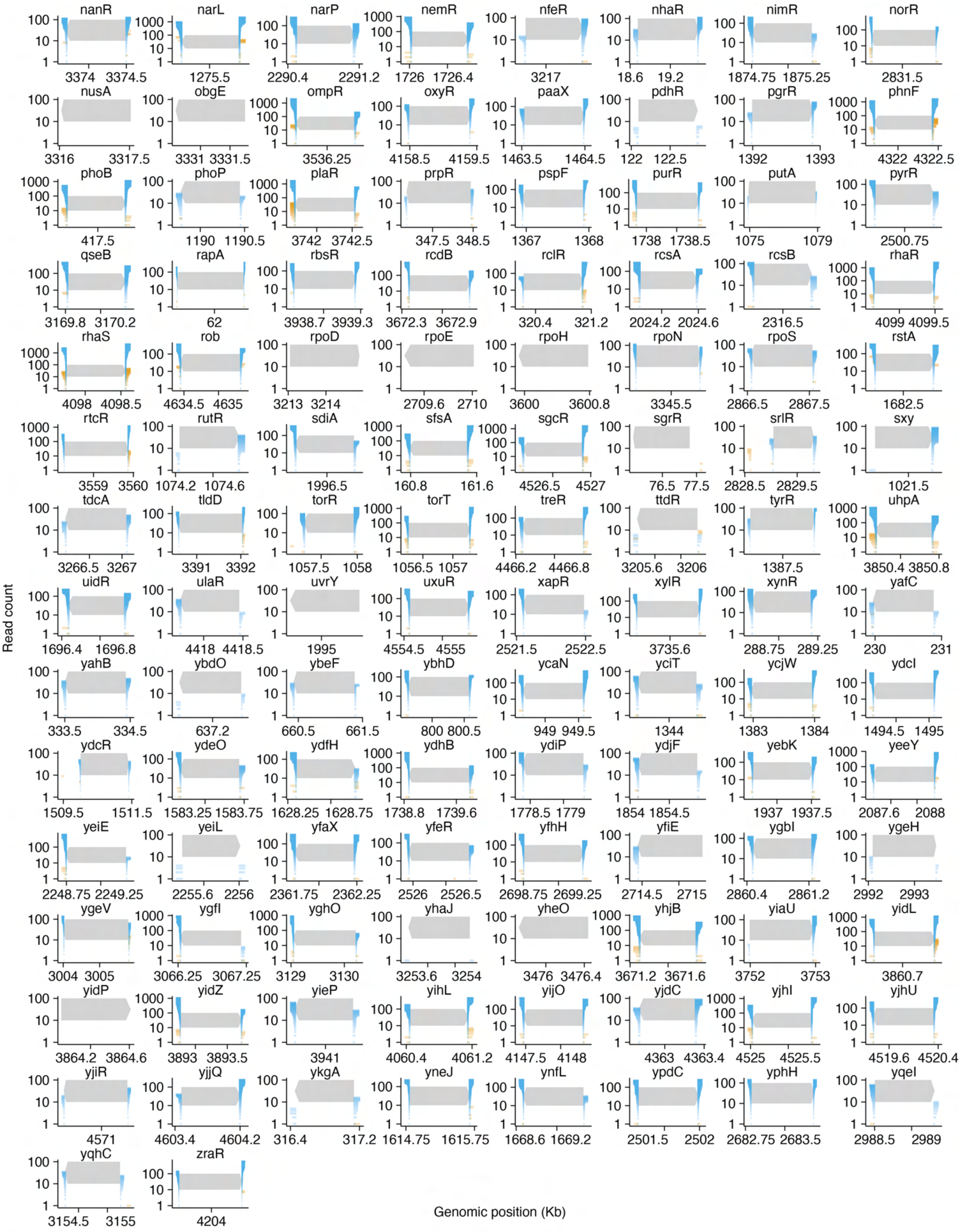
Read count
Genomic position (Kb)
nanR 3374 3374.5
narL 1275.5
narP 2290.4 2291.2
nemR 1726 1726.4
nfeR 3217
nhaR 18.6 19.2
nimR 1874.75 1875.25
norR 2831.5
nusA 3316 3317.5
obgE 3331 3331.5
ompR 3536.25
oxyR 4158.5 4159.5
paaX 1463.5 1464.5
pdhR 122 122.5
pgrR 1392 1393
phnF 4322 4322.5
phoB 417.5
phoP 1190 1190.5
plaR 3742 3742.5
prpR 347.5 348.5
pspF 1367 1368
purR 1738 1738.5
putA 1075 1079
pyrR 2500.75
qseB 3169.8 3170.2
rapA 62
rbsR 3938.7 3939.3
rcdB 3672.3 3672.9
rclR 320.4 321.2
rcsA 2024.2 2024.6
rcsB 2316.5
rhaR 4099 4099.5
rhaS 4098 4098.5
rob 4634.5 4635
rpoD 3213 3214
rpoE 2709.6 2710
rpoH 3600 3600.8
rpoN 3345.5
rpoS 2866.5 2867.5
rstA 1682.5
rtcR 3559 3560
rutR 1074.2 1074.6
sdiA 1996.5
sfsA 160.8 161.6
sgcR 4526.5 4527
sgrR 76.5 77.5
srlR 2828.5 2829.5
sxy 1021.5
tdcA 3266.5 3267
tldD 3391 3392
torR 1057.5 1058
torT 1056.5 1057
treR 4466.2 4466.8
ttdR 3205.6 3206
tyrR 1387.5
uhpA 3850.4 3850.8
uidR 1696.4 1696.8
ulaR 4418 4418.5
uvrY 1995
uxuR 4554.5 4555
xapR 2521.5 2522.5
xylR 3735.6
xynR 288.75 289.25
yafC 230 231
yahB 333.5 334.5
ybdO 637.2
ybeF 660.5 661.5
ybhD 800 800.5
ycaN 949 949.5
yciT 1344
ycjW 1383 1384
ydcI 1494.5 1495
ydcR 1509.5 1511.5
ydeO 1583.25 1583.75
ydfH 1628.25 1628.75
ydhB 1738.8 1739.6
ydiP 1778.5 1779
ydjF 1854 1854.5
yebK 1937 1937.5
yeeY 2087.6 2088
yeiE 2248.75 2249.25
yeiL 2255.6 2256
yfaX 2361.75 2362.25
yfeR 2526 2526.5
yfhH 2698.75 2699.25
yfiE 2714.5 2715
ygbI 2860.4 2861.2
ygeH 2992 2993
ygeV 3004 3005
ygfI 3066.25 3067.25
yghO 3129 3130
yhaJ 3253.6 3254
yheO 3476 3476.4
yhjB 3671.2 3671.6
yiaU 3752 3753
yidL 3860.7
yidP 3864.2 3864.6
yidZ 3893 3893.5
yieP 3941
yihL 4060.4 4061.2
yijO 4147.5 4148
yjdC 4363 4363.4
yjhI 4525 4525.5
yjhU 4519.6 4520.4
yjiR 4571
yjjQ 4603.4 4604.2
ykgA 316.4 317.2
yneJ 1614.75 1615.75
ynfL 1668.6 1669.2
ypdC 2501.5 2502
yphH 2682.75 2683.5
yqeI 2988.5 2989
yqhC 3154.5 3155
zraR 4204

Figure S14b. Long transcription factor deletion library - Upstream and downstream reads for target loci. Plots show all reads mapping within 500 bp of either end of the target genes. Orange segments show imperfect reads and blue segments show perfectly matched reads. The number of reads in each category, upstream – perfect, upstream – imperfect, downstream – perfect, downstream – imperfect are shown on the y-axis and reads are ordered by read length. Genomic coordinates are shown on the x-axis.

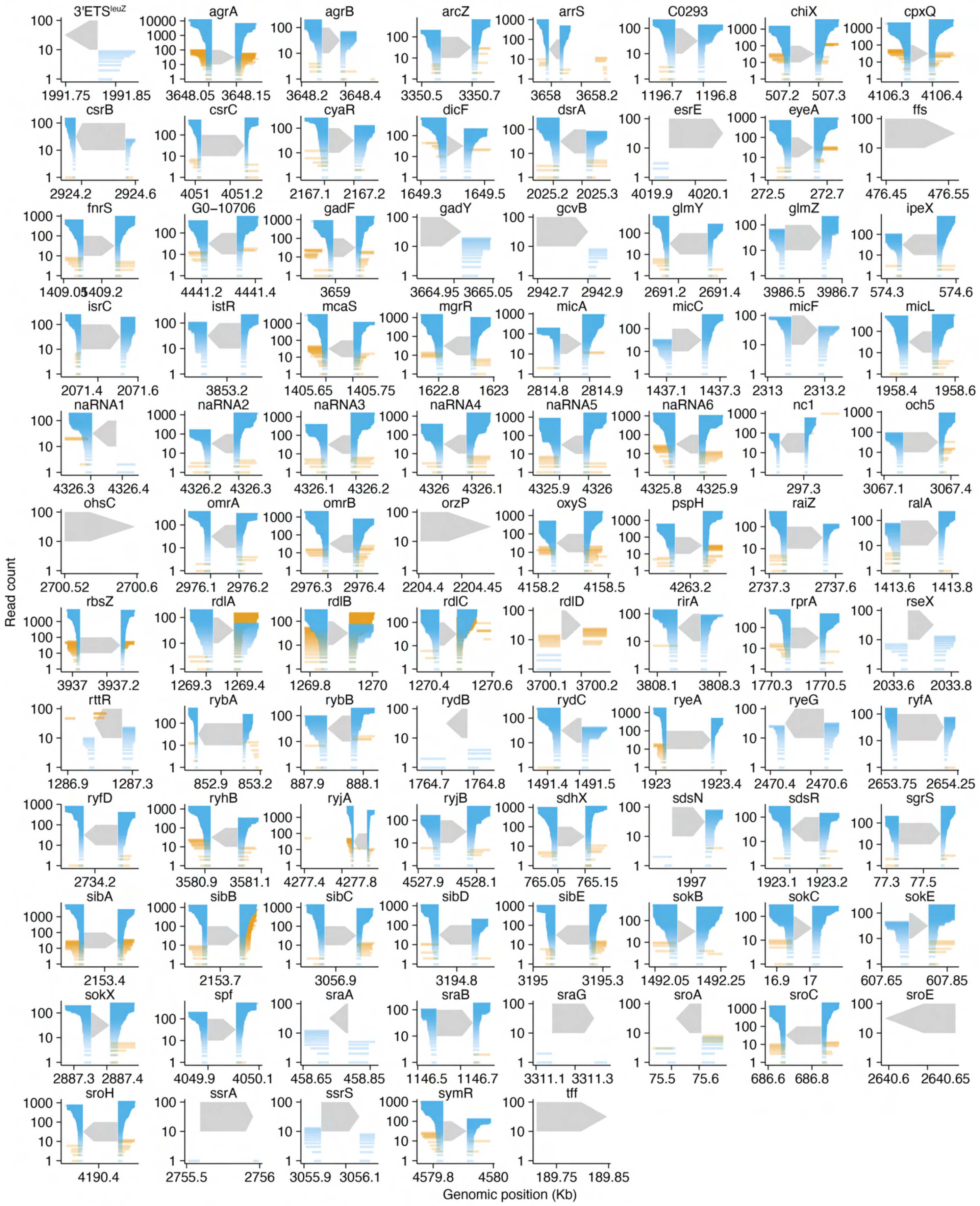


Figure S15. Small RNA deletion library - Upstream and downstream reads for target loci. Plots show all reads mapping within 500 bp of either end of the target genes. Orange segments show imperfect reads and blue segments show perfectly matched reads. The number of reads in each category, upstream – perfect, upstream – imperfect, downstream – perfect, downstream – imperfect are shown on the y-axis and reads are ordered by read length. Genomic coordinates are shown on the x-axis.

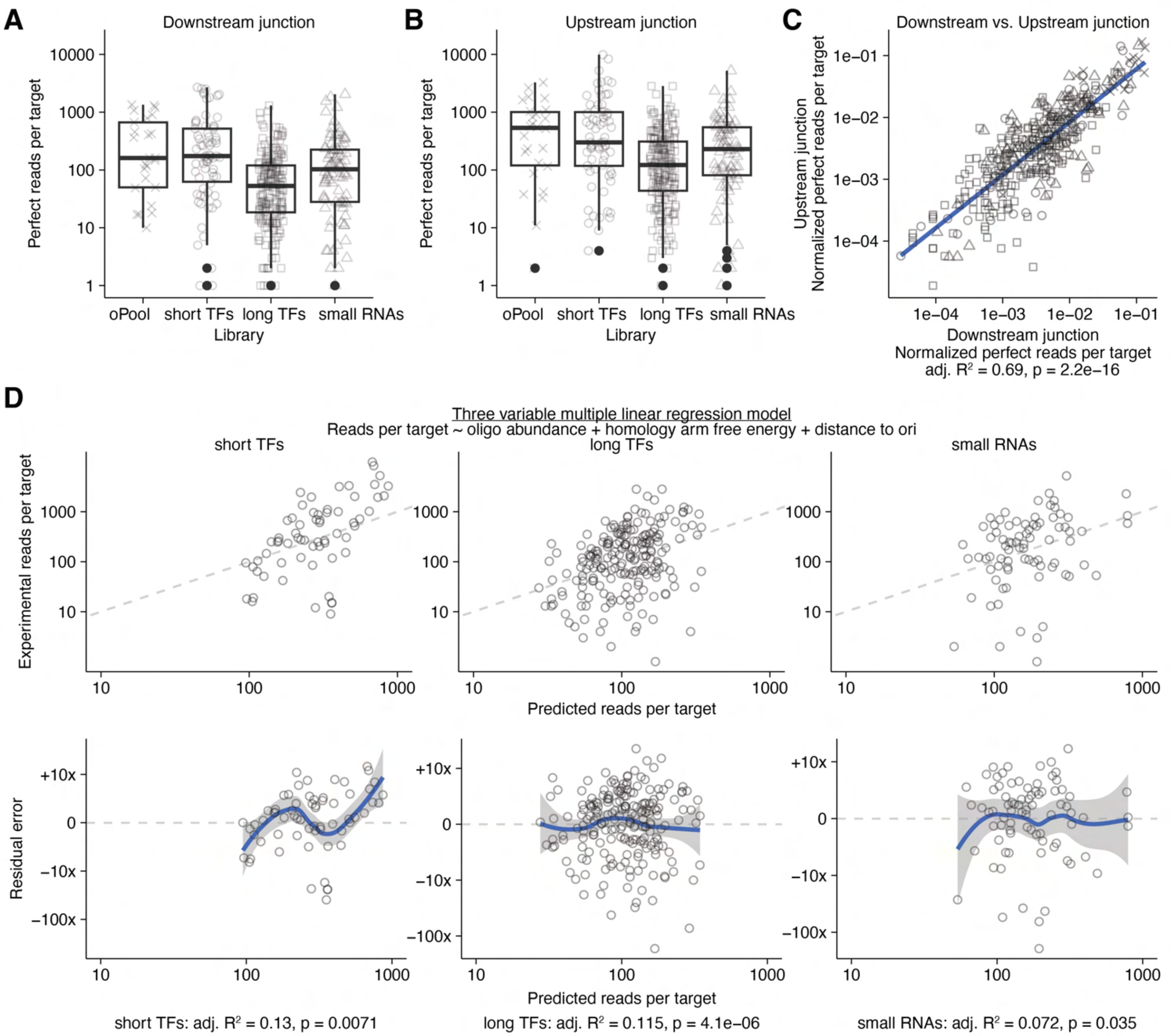


Figure S16. Explaining mutant library abundances. Sequencing results are shown for the downstream (A) and upstream (B) junctions. Perfect reads per target are shown for each of the 4 libraries, including the oPool library from Fig. 7. C) Upstream and downstream read counts correlate for all libraries (shown as different symbols). Read counts were normalized for each library by total reads and therefore represent relative abundance within the libraries. The blue line shows a linear model fit to the data, with $R^2$ and p-values shown below. D) A multiple linear regression model with three independent variables was fit to the reads per target data for the short TFs, long TFs and small RNAs libraries. The top panels show the predicted reads per target from the models compared to experimental observed values. A perfect model would have all points lie on the gray dashed line (slope = 1, intercept = 0). The bottom panels show the residuals from the model. A perfect model would lie on the gray dashed line at zero error. Blue lines show loess smoothing with a 95% confidence interval in gray. The $R^2$ and p-values for each model are shown below. The model was fit on $\log_{10}$(perfect_reads_per_target), therefore the residual error is a log difference.

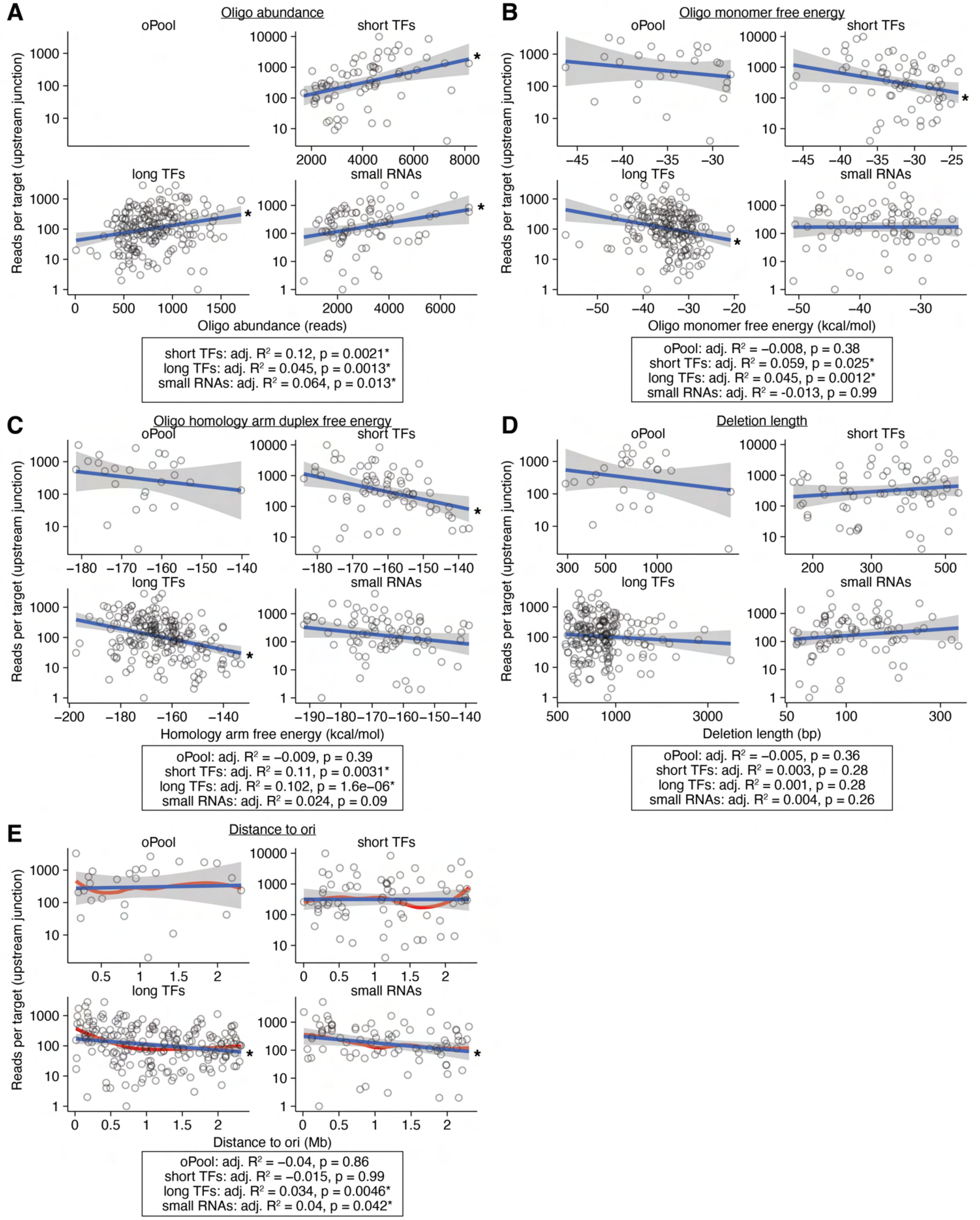


Figure S17. Correlations of oligo parameters and observed reads per target (perfect reads). The correlations between reads per target and oligo abundance (A), monomer free energy (B), homology arm free energy (C), deletion length (D) and distance to ori (E) are shown. Each open circle represents a target gene and the data are shown for each of the four libraries in Fig. 7C-D (oPool) and 8. Blue lines show linear models fit on $\log_{10}$(perfect_reads_per_target) and shaded areas show 95% confident intervals. Results from each statistical test are shown below each panel. Statistically significant correlations are denoted with * ($p < 0.05$). No data was obtained for oligo abundance on the oPool, since there were no conserved flanking primer sites available to construct amplicon sequencing libraries. The red line in panel E shows loess smoothing of the data, which suggests the relationship is nonlinear for the long TFs library (same data as Fig. 8H).
