## Supplementary material for "ORBIT for *E. coli*: Kilobase-scale oligonucleotide recombineering at high throughput and high efficiency": Text S1. Induced electrocompetent cell protocol

### ORBIT electrocompetent cells with induced helper plasmid

Scott H. Saunders, UT Southwestern Medical Center, February 6, 2023

Please send any feedback to

This protocol provides simple instructions for preparing induced electrocompetent cells for ORBIT genetics. It is assumed that you already have your strain of interest with the ORBIT pHelper plasmid. The goal of this protocol is to do two things – make electrocompetent cells that can electroporated with ORBIT materials and induce the oligo recombineering machinery on the helper plasmid. If done correctly, these cells should enable efficient ORBIT modifications to be made, and aliquots can be conveniently stored at -80°C for long time periods. Therefore, you just have to take an aliquot of cells from the freezer and go straight into the ORBIT integration protocol.

This procedure was written for a large prep (500 mL culture), but it can be scaled down to whatever volume is required and cells can be used fresh instead of frozen. Also, labs make competent cells in slightly different ways. Feel free to follow this protocol or modify as necessary.

#### Materials

Specialty materials

- Strain of interest with helper plasmid
- m-toluic acid (1M stock solution in ethanol)
- 10% glycerol (sterilized and ice cold)

*This inducer is cheap and easy to use and available from sigma. Make a 1M stock in ethanol.*

Common materials

- LB broth
- Large flasks
- MilliQ water (ice cold)
- Tray or bath of ice
- Dry ice

Common equipment

- Shaking incubator

**<u>Procedure</u>**

1. **Grow an overnight culture of the helper plasmid strain.** From a frozen stock or a colony, grow a 3 mL LB culture of the helper plasmid strain and make sure to supplement with the appropriate antibiotic.

*The next morning*

2. **Inoculate the main culture with the overnight (1:1000).** Prepare a large flask with 500 mL of LB supplemented with the appropriate antibiotic and add cells from the overnight culture 1:1000. Grow at 37°C with shaking (or modify accordingly) until the culture reaches OD ~0.3. This should take about 3 hours.

*Set up the water and glycerol on ice before you induce to make sure they are very cold.*

3. **Induce with 1 mM m-toluic acid for 30 min.** Add m-toluic acid directly from the 1M ethanol stock into the culture 1:1000 for a final concentration of 1 mM. Let the cells continue to grow and induce in the shaking incubator for 30 min.

4. **Chill culture in ice bath for 15 min.** After induction, transfer the flask to a water + ice bath and swirl the culture every few min for 15 min to evenly cool the cells.

5. **Spin the cells down.** For a 500 mL culture, transfer 45-50 mL into 10x 50mL conical vials. Spin the tubes at 5,000x g for 10 min. Pellets should be small (compared to plasmid preps), but clearly visible along the wall of the tubes. Pour off the supernatant.

6. **Wash 1x with ice cold water.** With a serological pipette resuspend the pellet in 45-50 mL of ice cold water. Again, spin the tubes at 5,000x g for 10 min, and carefully pour off the supernatant.

7. **Wash 2x with ice cold 10% glycerol.** With a serological pipette resuspend the pellet in 45-50 mL of ice cold 10% glycerol. Again, spin the tubes at 5,000x g for 10 min, and carefully pour off the supernatant. **Repeat this step.**

*During the final wash steps, start setting up the microfuge tubes for cell aliquots.*

8. **Resuspend cells and aliquot.** Resuspend the pellet in the residual glycerol solution. This should yield ~5 mL for a 500 mL culture. Of course, more concentrated cells can be made, but 100x concentration generally works well. Arrange many sterile microfuge tubes on ice and aliquot ~50 uL into each. This should yield ~100 aliquots.

9. **Freeze cell aliquots on dry ice, then transfer to -80°C freezer.** Cap the tubes and incubate on dry ice for 5 min before transferring to the -80°C freezer for long term storage.

10. **Test the aliquots.** Try ORBIT with a positive control to test your newly made cell aliquots. If these cells aren’t prepared properly then ORBIT will certainly fail.
