## Supplementary material for "ORBIT for *E. coli*: Kilobase-scale oligonucleotide recombineering at high throughput and high efficiency": Text S2. ORBIT Integration protocol

Scott H. Saunders, UT Southwestern Medical Center, February 6, 2023

Please send any feedback to

This protocol walks through the basic steps of how to perform an ORBIT integration. This protocol is general and should work for a variety of integration plasmids and targeting oligos that can be used for many different types of insertions and deletions on the *E. coli* genome. Please refer to other materials for a detailed description of ORBIT. As an overview, recall that ORBIT modifications require three components that work together: the helper plasmid, an integrating plasmid, and a targeting oligo. The targeting oligo contains a 38 bp attachment site (attB) flanked by homology arms and it gets incorporated into the genome during DNA replication directed by its homology arms. The integrating plasmid contains a 48 bp cognate attachment site (attP), which recombines with the oligo attachment site, thus integrating the entire plasmid at the location targeted in the oligo. The oligo incorporation, also called single stranded DNA recombineering (or MAGE for many oligos), is catalyzed by a single stranded DNA annealing protein (SSAP), and the modification is stabilized by temporarily suppressing the mismatch repair machinery. The recombination between the oligo attB site and the plasmid attP site is catalyzed by the site-specific recombinase Bxb-1. The helper plasmid contains all the necessary genes to achieve these reactions: SSAP - CspRecT, MutL E32K, and Bxb-1. Note that CspRecT and MutL E32K are inducibly controlled by m-toluic acid (XylS - $P_m$ system) and Bxb-1 is separately inducible with L-arabinose (AraC - $P_{araB}$ system). Here we assume that you already have a helper plasmid strain that has been induced with m-toluic acid (CspRecT + MutL E32K) and is electrocompetent. If you do not have this, please first complete the ORBIT helper strain protocol. The helper plasmid also has *sacB*, which confers sucrose sensitivity. Therefore, the helper plasmid can be removed from the newly created strain by selecting on sucrose plates.

### Materials

Specialty materials

- Targeting oligo - 25 µM stock
- Integrating plasmid - ~100 ng/µL stock
- Electrocompetent cells with induced helper plasmid - (~50uL fresh or frozen aliquot)

Common materials

- LB broth
- 10% L-arabinose (filter sterilized)
- LB + Kan plates (30 ug/mL)
- LB + Kan + 7.5% Sucrose plates

*Autoclave 125mL 30% sucrose (wt/vol) and 375mL $H_2O$ + 20g LB agar. Cool agar and mix.*

- LB plates (optional)
- 1 mm Electroporation cuvettes

Common equipment

- Electroporator
- Shaking incubator
- Plate incubator
- Ice maker

### **Procedure**

*Before you start*

1. **Consider setting up a control condition** WITH integrating plasmid but NO targeting oligo. This control will help you assess the likelihood that your ORBIT experiment worked before performing any molecular or phenotypic confirmation. It tests the background off target integration and provides a useful comparison to your + oligo conditions that should be a much higher on target integration. Typically, the OFF target integration rate should be ~ 1% of the ON target integration rate.

*Set up and electroporate*

2. **Set up electroporation cuvettes.** Unwrap and label cuvettes and cool on ice.

3. **Thaw frozen competent cells** (with induced helper plasmid) on ice for 10 min.

4. **Prepare the recovery culture tubes** while competent cells are thawing. With sterile technique, add 30 µL or 300 µL of L- arabinose stock (0.1 - 1% final) and 3 mL of LB to each culture tube. You may also thaw other stock solutions during this time.

Arabinose induces the Bxb-1 integrase. Note that cells need to divide during the recovery, so 3 mL cultures allow recovery cultures to be less dense and more likely to divide than smaller volumes.

5. **Add ORBIT materials to competent cells**. Add ~100 ng of integrating plasmid to each competent cell aliquot (typically ~1 µL). Add 2 µL of targeting oligo stock (~1 µM final) to each competent cell aliquot, as appropriate (do not include for off target control).

6. **Transfer to electroporation cuvettes**. Gently mix competent cell mixtures with pipette (3x) and transfer to electroporation cuvettes - try to avoid bubbles. Tap cuvettes gently to eliminate bubbles and provide good luck. Replace cuvettes in ice.

7. **Electroporate**. Carry cuvettes, recovery tubes, and 1 mL pipette (with tips) to the electroporator. Electroporate with typical E. coli settings 1.8 kV, 25 µF, 200 Ω. Immediately resuspend cuvette cells in 1 mL of recovery medium from the respective culture tube. Be gentle, and transfer the cells to the culture tube.

Electroporate and resuspend all conditions and then proceed to recovery.

*Recover and plate*

8. **Recover electroporated cells**. Transfer the recovery tubes to a 37°C shaker (~250 rpm) and recover for ~ 1 hour. Recovery time has been optimized for a 1 kb deletion with no payload on the integrating plasmid. Lower efficient modifications may require longer recovery periods.

For simply making mutants or cloning, it is always fine to recover longer. You may not be gaining any efficiency (recombinants / total cells), but because the culture grows you may get more recombinants in absolute terms.

9. **Prepare plates**. During recovery make sure agar plates are at least at room temperature. Dry plates with lids open during this time, if necessary.

10. **Plate recovered cells**. Plate a dilution series on Kan plates.

<u>Optional</u>: plate on LB as well as Kan - this allows you to calculate the overall efficiency (of all surviving cells, how many got the modification). Typically, overall efficiency is ~0.5% of all cells for a 1kb galK deletion.

Because of the high efficiency, 50 µL of undiluted culture can yield a lawn of colonies, so I strongly recommend trying multiple dilutions or doing a streak plate with an aliquot of the recovery culture. If low efficiency is observed, concentrate recovery culture by centrifugation and plate all cells.

<u>Optional</u>: The preferred method is the drip plate, which can be used to accurately count 10^1 - 10^9 colonies on a single plate. Simply dilute culture serially (10 fold) ~7 times in a 96 well plate. Plate 10 uL splots on to a plate and then tilt the plate so that the spots drip down it. Use a multichannel pipette to dilute and plate. Dry plates work best for this. The most common mode of failure is that drips run together, making it impossible to count accurately.

*Confirm mutants and remove helper plasmid*

11. **Check mutants.** Pick single colonies into water to set up the PCR and <u>save these suspensions</u>. Typically, we check insertions by colony PCR. There are many different ways to do this using primers to amplify the entire locus + plasmid insertion or using 1 primer that binds the genome and 1 primer that binds the plasmid insertion.

12. **Remove helper plasmid.** Grow correct colonies in LB + Kan (do not add the helper plasmid antibiotic) until dense or overnight and then plate 10-100uL on LB + Kan + 7.5% sucrose.

You should be left with your mutant of interest without the helper plasmid! You can confirm this by restreaking on Kan and separately on the helper plasmid antibiotic (e.g. gent).

If you want to remove the plasmid backbone / kan resistance you may transform with pCP20 and do FLP excision, as described elsewhere.

Note: Instead of plating for integrations on Kan then sucrose counter selecting against the helper plasmid, it is possible to directly plate for cells that incorporated the integrating plasmid and also lost the helper plasmid. To do this, recover ORBIT electroporations at least 3 hours (ideally overnight) then plate directly on LB + Kan + 7.5% sucrose. Concentrate the culture by spinning and plate all cells if culture has not grown overnight. This faster protocol can work, but it takes a surprising number of generations for cells to lose the helper plasmid even without the antibiotic, so the perceived efficiency will be much lower.

Troubleshooting: In general, there are four categories of issues-

1. Cell electrocompetence
2. Helper plasmid induction
3. Integrating plasmid quality
4. Targeting oligo quality

1. Cell electrocompetence

If you don't prepare electrocompetent cells well, they can all die during the electric pulse. You can assess this by plating on LB without antibiotics. You can also try transforming a positive control plasmid that replicates, instead of the ORBIT integrating plasmid. Just make sure it doesn't conflict with the helper plasmid ori.

2. Helper plasmid induction

The oligo recombineering and integrase modules must be properly induced to get ORBIT to work. That means inducing with m-toluic acid for 30 min during competent cell prep and recovering in 0.1% or 1% arabinose. If you forget either one of these you probably will not get any recombinants.

3. Integrating plasmid quality

The number of recombinants scales roughly linearly with the amount of clean plasmid added to the electroporation. If you get very few colonies, it may be because the plasmid is not very concentrated or clean. We have had issues with genomic DNA in our minipreps, which causes the perceived concentration of the plasmid to be much higher that it truly is. Check this by running your supercoiled plasmid on a gel. Also, you can include a positive control ORBIT integration plasmid that you know works well. If the positive control works well, but the new plasmid does not, then you know that's the problem.

4. Targeting oligo quality

If the targeting oligo is mismatched relative to the genome or you try to target a sequence that actually doesn't exist in the strain, then it will fail! This has happened. Assess this by using a positive control oligo, for example the 1kb galK deletion oligo. More broadly, there may be rare issues with targeting oligos that affect efficiency, like strong secondary structure, poor synthesis yields, inefficient regions of the genome etc. These are mostly unknown at this point, but in general longer oligos are more efficient and shorter deletions are more efficient.

If you are confident that everything is right, but efficiency is just low then make sure to plate the entire recovery culture and screen colonies. If this is a mutation you must make repeatedly, then it could be worth optimizing the protocol further (concentrations, comp cell prep, recovery etc.).
