## Supplementary material for "ORBIT for *E. coli*: Kilobase-scale oligonucleotide recombineering at high throughput and high efficiency": File S1. DNA maps (.gb): addgene_plasmid_maps.nb.html

R Notebook


Code 

- Show All Code
- Hide All Code
- Download Rmd

### R Notebook


```
```r
plot(cars)
```

```
<!-- rnb-source-end -->

<!-- rnb-chunk-end -->


<!-- rnb-text-begin -->


<!-- rnb-text-end -->


<!-- rnb-chunk-begin -->


<!-- rnb-frame-begin eyJtZXRhZGF0YSI6eyJjbGFzc2VzIjpbImRhdGEuZnJhbWUiXSwibnJvdyI6MjAsIm5jb2wiOjYsInN1bW1hcnkiOnsiRGVzY3JpcHRpb24iOlsiZGYgWzIwIMOXIDZdIl19fSwicmRmIjoiSDRzSUFBQUFBQUFBQTdWVnpXOGJSUlRmdUhHKzJwVFFSb2lqanlGU1FwTklWVzlrc2pPZVhlM1l6TTRNcy9FcFd0dmIxcXE5YTYwM2FpcHhpRGdneElmRWlRTVNFaGZnaUJBU04xcTFsVGp4RDNEaXpva0xOeWk4OWM2Z3hLblVDZ2xMYjM5djNyeVAzeHZQaDhDSGV5dUhLNDdqWEhMbWEvQ3RnK3JVMzFITnJWdk8xT0xNT2ZQT2Nva25NSDBkbEFXUVZaRDE2YVRqMUtiUjRGMUdtdmxGa0NXUU1yRE1maG5raW9tN0N2SUt5QnJJcXlEWFFLNmJmR3RUckFyVzNXeElkc3pnV3A2bkJ3MjEyeWlTZkRSSTR5TEx6Y3pxblNRdFJHTWpqbnZ1emh2R09CL25zV3YwR3U5WjYrR0RvVFQ2WlQ1cWpQTnNsRUZDWTFweUoyT1I5RlREamx2SEJTTjd1NEVacncvalViY2ZONG9iRjFuVU9WUThNSU5MNHNDV2VYMnYwUjAzSnVPNGwrU05qY2x4TnhzWGcxRTh0RVNYdXlmZHJaMkdueGFWWWU2UGxrQk1hMTh3S2JUQ1NDbkppUWdaa1c0WU1oZUZna1pZYTZvUndReHJTWkhHdklsRklBUnZNeVFpaHBxRWhHRFN2a1lkb2JGZ1NraGZlQXppdFJkQm5CY0VVRUlpUlpEWHhFb3Aza1FJYWlMV29rcWpzRVVZUVg1QVJGdElCQzV0WHlBYWRBU1ZqUE9JZGhoWGtKZVNTREROTWRlaElNSm5SSGM4b1RWdWU0UXpqMm5rWWRZV2dtb21lWUE3VFJTeWtBckN5MDhFeFJFTGhHejVrcUFXb3gya1dKc0dTZ3VNTVVXY2FZRUk5NEU2Z2Q0Umd4SFRnWklDQlJ4cHFiaGtqQW5ObWk3U0xpVWNkWUltb3BTS3dPT2lJMXpJVEpzODhGemFMbHRxSVJKaDFIUmhaVFREakZHTWlHQWswSkZHVkdLSk5HR0k2SFpaQlRNbG1VOGw1SVMySllmdUVLeTA4QkZpQ0ltUXdPcFE0Q1M0cEJHc0FWRTBGRTJLSTFCRHJCQUsyaXBpUUtDdFllbmhmeENVUWgvS3grREVOa21IaEI0VGpFcXRnWU9uYUV0dTJvMGoyOXlvVjhkNU1vcUw0enhwU1BVMlAzOE1kaTV1d0hrWjk4UVovV0RtTksza3lmZ295d2QzQnFtMVhNd0JSeW14VkZ4czkvQ1YwV0RTTzdxZFRObGNuRjZhT1ViUGMzK2U3V0w5bDQxOFdkc0xldmp2cEthSnE1dHMvYTNmbmVsdi84TlBLL3lFVi9qRnp4VitYVG5zZi9OdWhkOHZHdnl5d2g5K3JmRGhrY0hmS255MGFmQ0d3YjN6K1BqUENwL2NOZmk1d1Y5T3AvajA0OU9LV2NWei96M2o5NUhCejB6OFYwOE12NDBLdjEwdy9DTER6L1R4OERXRFB4b2VkWU52R3R3MTlUODRQWWVQL3pMMmdjSHZLbno2Zm9VL1NlY3NUN3VlRmgrOUFHZjkvMitjZVJicmFUeEtKazUxMkJhc2NaRDJreE43cWtvUHF4Y1B4bGF2VDRvNEwreU9TdEsrZlJMNmd6enBGWU1zblNtMW5HZjN0MjI1OGlXdG5WYUUxbWM1OVlieHhIS3l4cFYrWE1UYnQvT1NqT004bXdsWkxOK2xMSVdnV3ZrVzEyZUM1L0ladzlweFdqTHBiL1h1SHFmM3RtN2RkS3FIZnM2czBhclIxOC9vQzFYTjJ0OG1WOTNrV2toU3VKWCtYWlpoM0UyRzloYUVscWNkYjQvemdYMGo0UzdMN2srMmk2eUlyZDlLTHh0YXk3UTU1OWsvV202Yjh0c0lBQUE9In0= -->

<div data-pagedtable="false">
  <script data-pagedtable-source type="application/json">
{"columns":[{"label":["index"],"name":[1],"type":["int"],"align":["right"]},{"label":["name"],"name":[2],"type":["chr"],"align":["left"]},{"label":["type"],"name":[3],"type":["chr"],"align":["left"]},{"label":["start"],"name":[4],"type":["dbl"],"align":["right"]},{"label":["end"],"name":[5],"type":["dbl"],"align":["right"]},{"label":["direction"],"name":[6],"type":["dbl"],"align":["right"]}],"data":[{"1":"1","2":"ColE1","3":"rep_origin","4":"1","5":"589","6":"1"},{"1":"2","2":"rrnB T2 terminator","3":"terminator","4":"850","5":"877","6":"1"},{"1":"3","2":"gentR (aacC1)","3":"gene","4":"938","5":"1471","6":"-1"},{"1":"4","2":"araC","3":"CDS","4":"1907","5":"2785","6":"-1"},{"1":"5","2":"Pc","3":"misc_feature","4":"2936","5":"2964","6":"-1"},{"1":"6","2":"XylS","3":"CDS","4":"3006","5":"3971","6":"-1"},{"1":"7","2":"Pm promoter","3":"promoter","4":"4871","5":"4951","6":"1"},{"1":"8","2":"CspRecT","3":"misc_feature","4":"5024","5":"5836","6":"1"},{"1":"9","2":"MutLE32K","3":"misc_feature","4":"5855","5":"7702","6":"1"},{"1":"10","2":"lambda t0 terminator","3":"terminator","4":"7775","5":"7869","6":"1"},{"1":"11","2":"ParaB","3":"misc_feature","4":"7914","5":"7941","6":"1"},{"1":"12","2":"RBS","3":"misc_feature","4":"7978","5":"7983","6":"1"},{"1":"13","2":"3 bp spacer (suboptimal)","3":"misc_feature","4":"7984","5":"7986","6":"1"},{"1":"14","2":"bxb-1 Int","3":"misc_feature","4":"7987","5":"9489","6":"1"},{"1":"15","2":"MRALVVIRLSRVTDATTSPERQLESCQQLCAQRGWDVVGVAEDLDVSGAVDPFDRKRRPNLARWLAFEEQPFDVIVAYRVDRLTRSIRHLQQLVHWAEDHKKLVVSATEAHFDTTTPFAAVVIALMGTVAQMELEAIKERNRSAAHFNIRAGKYRGSLPPWGYLPTRVDGEWRLVPDPVQRERILEVYHRVVDNHEPLHLVAHDLNRRGVLSPKDYFAQLQGREPQGREWSATALKRSMISEAMLGYATLNGKTVRDDDGAPLVRAEPILTREQLEALRAELVKTSRAKPAVSTPSLLLRVLFCAVCGEPAYKFAGGGRKHPRYRCRSMGFPKHCGNGTVAMAEWDAFCEEQVLDLLGDAERLEKVWVAGSDSAVELAEVNAELVDLTSLIGSPAYRAGSPQREALDVRIAALAARQEELEGLEARPSGWEWRETGQRFGDWWREQDTAAKNTWLRSMNVRLTFDVRGGLTRTIDFGDL*EYEQHLRLGSVVERLHTGMS*","3":"CDS","4":"7987","5":"9489","6":"1"},{"1":"16","2":"SNP","3":"misc_feature","4":"9208","5":"9210","6":"1"},{"1":"17","2":"premature STOP","3":"misc_feature","4":"9424","5":"9426","6":"1"},{"1":"18","2":"rrnB T1 terminator","3":"terminator","4":"9524","5":"9570","6":"1"},{"1":"19","2":"SacR","3":"misc_feature","4":"9651","5":"9996","6":"1"},{"1":"20","2":"SacB","3":"CDS","4":"10009","5":"11430","6":"1"}],"options":{"columns":{"min":{},"max":[10],"total":[6]},"rows":{"min":[10],"max":[10],"total":[20]},"pages":{}}}
  </script>
</div>

<!-- rnb-frame-end -->

<!-- rnb-chunk-end -->


<!-- rnb-text-begin -->


<!-- rnb-text-end -->


<!-- rnb-chunk-begin -->


<!-- rnb-source-begin eyJkYXRhIjoiYGBgclxucGxvdF9wbGFzbWlkKHAyLCBuYW1lID0gJ3BIZWxwZXJfRWMxX1YxX2FtcFInKSArIHNjYWxlX2ZpbGxfbWFudWFsKHZhbHVlcz1jKFwiI0E4QkRFMlwiLCBcIiNEMzZENTVcIiwgXCIjRThCRjYzXCIpKSArIHRoZW1lKHBhbmVsLmJhY2tncm91bmQgPSBlbGVtZW50X3JlY3QoZmlsbCA9ICd3aGl0ZScsIGNvbG9yID0gJ3doaXRlJykpXG5cbmBgYCJ9 -->

```r
plot_plasmid(p2, name = 'pHelper_Ec1_V1_ampR') + scale_fill_manual(values=c("#A8BDE2", "#D36D55", "#E8BF63")) + theme(panel.background = element_rect(fill = 'white', color = 'white'))
```


```
Scale for fill is already present.
Adding another scale for fill, which will replace the existing scale.
```


```
plot_plasmid(p3, name = 'pHelper_NoMutL_V2_gentR') + scale_fill_manual(values=c("#A8BDE2", "#D36D55", "#E8BF63")) + theme(panel.background = element_rect(fill = 'white', color = 'white'))
```


```
Scale for fill is already present.
Adding another scale for fill, which will replace the existing scale.
```


```
minor <- c(2,3,10)

pInt_kan <- read_gb(file = "pint_attp1_kanr.gb") %>% 
  as.data.frame() %>% 
  filter((name != 'pInt_seq_rev') & (name != 'pInt_seq_fwd') & (name != 'central_di_nt')) %>% 
  mutate(type = ifelse(name == 'bxb1_attP', 'mod_1', type)) %>% 
  mutate(type = ifelse(name == 'kanamycin resistance', 'mod_2', type)) %>% 
  mutate(type = ifelse((type != 'mod_1') & (type != 'mod_2'), 'mod_3', type)) %>% 
  mutate(name = ifelse(index %in% minor, '', name))

plot_plasmid(pInt_kan, name = 'pInt_attP1_kanR') + scale_fill_manual(values=c("#D36D55","#E8BF63", "#D3C0D6")) + theme(panel.background = element_rect(fill = 'white', color = 'white'))
```


```
Scale for fill is already present.
Adding another scale for fill, which will replace the existing scale.
```


```
ggsave('pInt_attP1_kanR.png',width = 6, height = 6)
```


```
ggsave('pInt_attP2_kanR.png',width = 6, height = 6)
```


```
NA
```


```
minor <- c(2,3,10)

pInt_kan <- read_gb(file = "pint_attp1_kanr.gb") %>% 
  as.data.frame() %>% 
  filter((name != 'pInt_seq_rev') & (name != 'pInt_seq_fwd') & (name != 'central_di_nt')) %>% 
  mutate(type = ifelse(name == 'bxb1_attP', 'mod_1', type)) %>% 
  mutate(type = ifelse(name == 'kanamycin resistance', 'mod_2', type)) %>% 
  mutate(type = ifelse((type != 'mod_1') & (type != 'mod_2'), 'mod_3', type)) %>% 
  mutate(name = ifelse(index %in% minor, '', name))

plot_plasmid(pInt_kan, name = 'pInt_attP2_kanR') + scale_fill_manual(values=c("#D36D55","#E8BF63", "#D3C0D6")) + theme(panel.background = element_rect(fill = 'white', color = 'white'))
```


```
Scale for fill is already present.
Adding another scale for fill, which will replace the existing scale.
```


```
ggsave('pInt_attP2_kanR.png',width = 6, height = 6)
```


```
NA
```


```
minor <- c(3,7,11)

pInt_chlor <- read_gb(file = "pint_attp1_chlorr.gb") %>% 
  as.data.frame() %>% 
  filter((name != 'pInt_seq_rev') & (name != 'pInt_seq_fwd') & (name != 'central_di_nt')) %>% 
  mutate(type = ifelse(name == 'bxb1_attP', 'mod_1', type)) %>% 
  mutate(type = ifelse(name == 'cat', 'mod_2', type)) %>% 
  mutate(type = ifelse((type != 'mod_1') & (type != 'mod_2'), 'mod_3', type)) %>% 
  mutate(name = ifelse(index %in% minor, '', name))

plot_plasmid(pInt_chlor, name = 'pInt_attP1_chlorR') + scale_fill_manual(values=c("#D36D55","#E8BF63", "#D3C0D6")) + theme(panel.background = element_rect(fill = 'white', color = 'white'))
```


```
Scale for fill is already present.
Adding another scale for fill, which will replace the existing scale.
```


```
ggsave('pInt_attP1_chlorR.png',width = 6, height = 6)
```


```
ggsave('pInt_attP3_chlorR.png',width = 6, height = 6)
```


```
NA
```


```
minor <- c(3,7,11)

pInt_chlor <- read_gb(file = "pint_attp1_chlorr.gb") %>% 
  as.data.frame() %>% 
  filter((name != 'pInt_seq_rev') & (name != 'pInt_seq_fwd') & (name != 'central_di_nt')) %>% 
  mutate(type = ifelse(name == 'bxb1_attP', 'mod_1', type)) %>% 
  mutate(type = ifelse(name == 'cat', 'mod_2', type)) %>% 
  mutate(type = ifelse((type != 'mod_1') & (type != 'mod_2'), 'mod_3', type)) %>% 
  mutate(name = ifelse(index %in% minor, '', name))

plot_plasmid(pInt_chlor, name = 'pInt_attP3_chlorR') + scale_fill_manual(values=c("#D36D55","#E8BF63", "#D3C0D6")) + theme(panel.background = element_rect(fill = 'white', color = 'white'))
```


```
Scale for fill is already present.
Adding another scale for fill, which will replace the existing scale.
```


```
ggsave('pInt_attP3_chlorR.png',width = 6, height = 6)
```


```
NA
```


```
minor <- c(3,7,11)

pInt_amp <- read_gb(file = "pint_attp1_ampr.gb") %>% 
  as.data.frame() %>% 
  filter((name != 'pInt_seq_rev') & (name != 'pInt_seq_fwd') & (name != 'central_di_nt')) %>% 
  mutate(type = ifelse(name == 'bxb1_attP', 'mod_1', type)) %>% 
  mutate(type = ifelse(name == 'bla', 'mod_2', type)) %>% 
  mutate(type = ifelse((type != 'mod_1') & (type != 'mod_2'), 'mod_3', type)) %>% 
  mutate(name = ifelse(index %in% minor, '', name))

plot_plasmid(pInt_amp, name = 'pInt_attP1_ampR') + scale_fill_manual(values=c("#D36D55","#E8BF63", "#D3C0D6")) + theme(panel.background = element_rect(fill = 'white', color = 'white'))
```


```
Scale for fill is already present.
Adding another scale for fill, which will replace the existing scale.
```


```
ggsave('pInt_attP1_ampR.png',width = 6, height = 6)
```


```
ggsave('pInt_attP5_ampR.png',width = 6, height = 6)
```


```
NA
```


```
minor <- c(3,7,11)

pInt_amp <- read_gb(file = "pint_attp1_ampr.gb") %>% 
  as.data.frame() %>% 
  filter((name != 'pInt_seq_rev') & (name != 'pInt_seq_fwd') & (name != 'central_di_nt')) %>% 
  mutate(type = ifelse(name == 'bxb1_attP', 'mod_1', type)) %>% 
  mutate(type = ifelse(name == 'bla', 'mod_2', type)) %>% 
  mutate(type = ifelse((type != 'mod_1') & (type != 'mod_2'), 'mod_3', type)) %>% 
  mutate(name = ifelse(index %in% minor, '', name))

plot_plasmid(pInt_amp, name = 'pInt_attP5_ampR') + scale_fill_manual(values=c("#D36D55","#E8BF63", "#D3C0D6")) + theme(panel.background = element_rect(fill = 'white', color = 'white'))
```


```
Scale for fill is already present.
Adding another scale for fill, which will replace the existing scale.
```


```
ggsave('pInt_attP5_ampR.png',width = 6, height = 6)
```


```
NA
```


```
integration = 12:18
minor <- c(3,10,12,14,16, 18)

pInt_lcs <- read_gb(file = "pint_attp1_lcs_kanr.gb") %>% 
  as.data.frame() %>% 
  filter(name != 'Integration Payload') %>% 
  filter((name != 'pInt_seq_rev') & (name != 'pInt_seq_fwd') & (name != 'central_di_nt')) %>% 
  mutate(type = ifelse(name == 'bxb1_attP', 'mod_1', type)) %>% 
  mutate(type = ifelse(name == 'kanamycin resistance', 'mod_2', type)) %>% 
  mutate(type = ifelse((type != 'mod_1') & (type != 'mod_2'), 'mod_3', type)) %>% 
  mutate(type = ifelse(index %in% integration, 'mod_4', type)) %>% 
  mutate(name = ifelse(index %in% minor, '', name))

plot_plasmid(pInt_lcs, name = 'pInt_attP1_LCS_kanR') + scale_fill_manual(values=c("#D36D55","#E8BF63", "#D3C0D6",'#BED395')) + theme(panel.background = element_rect(fill = 'white', color = 'white'))

ggsave('pInt_attP1_LCS_kanR.png',width = 6, height = 6)
```


```
minor <- c(3,11)

pInt_sacB <- read_gb(file = "pint_attp1_sacb_kanr.gb") %>% 
  as.data.frame() %>% 
  filter((name != 'pInt_seq_rev') & (name != 'pInt_seq_fwd') & (name != 'central_di_nt')) %>% 
  filter(name!='SacR') %>% 
  mutate(type = ifelse(name == 'bxb1_attP', 'mod_1', type)) %>% 
  mutate(type = ifelse(name == 'kanamycin resistance', 'mod_2', type)) %>% 
  mutate(type = ifelse((type != 'mod_1') & (type != 'mod_2'), 'mod_3', type)) %>% 
  mutate(type = ifelse(name == 'SacB', 'mod_4', type)) %>% 
  mutate(name = ifelse(index %in% minor, '', name))

plot_plasmid(pInt_sacB, name = 'pInt_attP1_sacB_kanR') + scale_fill_manual(values=c("#D36D55","#E8BF63", "#D3C0D6",'#BED395')) + theme(panel.background = element_rect(fill = 'white', color = 'white'))

ggsave('pInt_attP1_sacB_kanR.png',width = 6, height = 6)
```


```
plot_plasmid(pInt_xis, name = 'pInt_attP1_tsXis_sacB_kanR') + scale_fill_manual(values=c("#D36D55","#E8BF63", "#D3C0D6",'#BED395')) + theme(panel.background = element_rect(fill = 'white', color = 'white'))
```


```
Scale for fill is already present.
Adding another scale for fill, which will replace the existing scale.
```
