## Supplementary material for "ORBIT for *E. coli*: Kilobase-scale oligonucleotide recombineering at high throughput and high efficiency": File S1. DNA maps (.gb): addgene_plasmid_maps.Rmd

````
---
title: "R Notebook"
output: html_notebook
---


```{r}
library(tidyverse)

library(plasmapR)
```

```{r}

bxb1_mod <- c(4,11,12,13,14,16,17,18)

recT_mod <- c(6,7,8,9, 10)

minor <- c(2,10,12,13,16,18)

p1 <- read_gb(file = "phelper_ec1_v1_gentr.gb") %>% 
  as.data.frame() %>% 
  filter(index != 15) %>% 
  filter(index!=5) %>% 
  filter(name != 'SacR') %>% 
  mutate(type = ifelse(index %in% recT_mod, 'mod_1', type)) %>% 
  mutate(type = ifelse(index %in% bxb1_mod, 'mod_2', type)) %>% 
  mutate(type = ifelse((type != 'mod_1') & (type != 'mod_2'), 'mod_3', type)) %>% 
  mutate(name = ifelse(index %in% minor, '', name))

plot_plasmid(p1, name = 'pHelper_Ec1_V1_gentR') + scale_fill_manual(values=c("#A8BDE2", "#D36D55", "#E8BF63")) + theme(panel.background = element_rect(fill = 'white', color = 'white'))

ggsave('pHelper_Ec1_V1_gentR.png',width = 6, height = 6)
  
```

```{r}

bxb1_mod <- c(4,5,11,12,14,15,16)

recT_mod <- c(6,7,8,9, 10)

minor <- c(2,5,10,14,16)

p2 <- read_gb(file = "phelper_ec1_v1_ampr.gb") %>% 
  as.data.frame() %>% 
  filter(index!=13) %>% 
  filter(name != 'SacR') %>% 
  mutate(type = ifelse(index %in% recT_mod, 'mod_1', type)) %>% 
  mutate(type = ifelse(index %in% bxb1_mod, 'mod_2', type)) %>% 
  mutate(type = ifelse((type != 'mod_1') & (type != 'mod_2'), 'mod_3', type)) %>% 
  mutate(name = ifelse(index %in% minor, '', name))

plot_plasmid(p2, name = 'pHelper_Ec1_V1_ampR') + scale_fill_manual(values=c("#A8BDE2", "#D36D55", "#E8BF63")) + theme(panel.background = element_rect(fill = 'white', color = 'white'))

ggsave('pHelper_Ec1_V1_ampR.png',width = 6, height = 6)
  
```

```{r}

bxb1_mod <- c(4,5,10,11,12,13,16)

recT_mod <- c(6,7,8,9)

minor <- c(2,5,9,11,12,16)

p3 <- read_gb(file = "phelper_nomutl_v2_gentr.gb") %>% 
  as.data.frame() %>% 
  filter(index!=14) %>% 
    filter(index!=15) %>% 
  filter(name != 'SacR') %>% 
  mutate(type = ifelse(index %in% recT_mod, 'mod_1', type)) %>% 
  mutate(type = ifelse(index %in% bxb1_mod, 'mod_2', type)) %>% 
  mutate(type = ifelse((type != 'mod_1') & (type != 'mod_2'), 'mod_3', type)) %>% 
  mutate(name = ifelse(index %in% minor, '', name))

plot_plasmid(p3, name = 'pHelper_NoMutL_V2_gentR') + scale_fill_manual(values=c("#A8BDE2", "#D36D55", "#E8BF63")) + theme(panel.background = element_rect(fill = 'white', color = 'white'))

ggsave('pHelper_NoMutL_V2_gentR.png',width = 6, height = 6)
  
```

```{r}

minor <- c(2,3,10)

pInt_kan <- read_gb(file = "pint_attp1_kanr.gb") %>% 
  as.data.frame() %>% 
  filter((name != 'pInt_seq_rev') & (name != 'pInt_seq_fwd') & (name != 'central_di_nt')) %>% 
  mutate(type = ifelse(name == 'bxb1_attP', 'mod_1', type)) %>% 
  mutate(type = ifelse(name == 'kanamycin resistance', 'mod_2', type)) %>% 
  mutate(type = ifelse((type != 'mod_1') & (type != 'mod_2'), 'mod_3', type)) %>% 
  mutate(name = ifelse(index %in% minor, '', name))

plot_plasmid(pInt_kan, name = 'pInt_attP1_kanR') + scale_fill_manual(values=c("#D36D55","#E8BF63", "#D3C0D6")) + theme(panel.background = element_rect(fill = 'white', color = 'white'))

ggsave('pInt_attP1_kanR.png',width = 6, height = 6)

#ggsave('pInt_attP2_kanR.png',width = 6, height = 6)
  
```

```{r}

minor <- c(2,3,10)

pInt_kan <- read_gb(file = "pint_attp1_kanr.gb") %>% 
  as.data.frame() %>% 
  filter((name != 'pInt_seq_rev') & (name != 'pInt_seq_fwd') & (name != 'central_di_nt')) %>% 
  mutate(type = ifelse(name == 'bxb1_attP', 'mod_1', type)) %>% 
  mutate(type = ifelse(name == 'kanamycin resistance', 'mod_2', type)) %>% 
  mutate(type = ifelse((type != 'mod_1') & (type != 'mod_2'), 'mod_3', type)) %>% 
  mutate(name = ifelse(index %in% minor, '', name))

plot_plasmid(pInt_kan, name = 'pInt_attP2_kanR') + scale_fill_manual(values=c("#D36D55","#E8BF63", "#D3C0D6")) + theme(panel.background = element_rect(fill = 'white', color = 'white'))


ggsave('pInt_attP2_kanR.png',width = 6, height = 6)
  
```

```{r}

minor <- c(3,7,11)

pInt_chlor <- read_gb(file = "pint_attp1_chlorr.gb") %>% 
  as.data.frame() %>% 
  filter((name != 'pInt_seq_rev') & (name != 'pInt_seq_fwd') & (name != 'central_di_nt')) %>% 
  mutate(type = ifelse(name == 'bxb1_attP', 'mod_1', type)) %>% 
  mutate(type = ifelse(name == 'cat', 'mod_2', type)) %>% 
  mutate(type = ifelse((type != 'mod_1') & (type != 'mod_2'), 'mod_3', type)) %>% 
  mutate(name = ifelse(index %in% minor, '', name))

plot_plasmid(pInt_chlor, name = 'pInt_attP1_chlorR') + scale_fill_manual(values=c("#D36D55","#E8BF63", "#D3C0D6")) + theme(panel.background = element_rect(fill = 'white', color = 'white'))

ggsave('pInt_attP1_chlorR.png',width = 6, height = 6)

ggsave('pInt_attP3_chlorR.png',width = 6, height = 6)
  
```

```{r}

minor <- c(3,7,11)

pInt_chlor <- read_gb(file = "pint_attp1_chlorr.gb") %>% 
  as.data.frame() %>% 
  filter((name != 'pInt_seq_rev') & (name != 'pInt_seq_fwd') & (name != 'central_di_nt')) %>% 
  mutate(type = ifelse(name == 'bxb1_attP', 'mod_1', type)) %>% 
  mutate(type = ifelse(name == 'cat', 'mod_2', type)) %>% 
  mutate(type = ifelse((type != 'mod_1') & (type != 'mod_2'), 'mod_3', type)) %>% 
  mutate(name = ifelse(index %in% minor, '', name))

plot_plasmid(pInt_chlor, name = 'pInt_attP3_chlorR') + scale_fill_manual(values=c("#D36D55","#E8BF63", "#D3C0D6")) + theme(panel.background = element_rect(fill = 'white', color = 'white'))

ggsave('pInt_attP3_chlorR.png',width = 6, height = 6)
  
```

```{r}

minor <- c(3,7,11)

pInt_amp <- read_gb(file = "pint_attp1_ampr.gb") %>% 
  as.data.frame() %>% 
  filter((name != 'pInt_seq_rev') & (name != 'pInt_seq_fwd') & (name != 'central_di_nt')) %>% 
  mutate(type = ifelse(name == 'bxb1_attP', 'mod_1', type)) %>% 
  mutate(type = ifelse(name == 'bla', 'mod_2', type)) %>% 
  mutate(type = ifelse((type != 'mod_1') & (type != 'mod_2'), 'mod_3', type)) %>% 
  mutate(name = ifelse(index %in% minor, '', name))

plot_plasmid(pInt_amp, name = 'pInt_attP1_ampR') + scale_fill_manual(values=c("#D36D55","#E8BF63", "#D3C0D6")) + theme(panel.background = element_rect(fill = 'white', color = 'white'))

ggsave('pInt_attP1_ampR.png',width = 6, height = 6)

ggsave('pInt_attP5_ampR.png',width = 6, height = 6)
  
```

```{r}

minor <- c(3,7,11)

pInt_amp <- read_gb(file = "pint_attp1_ampr.gb") %>% 
  as.data.frame() %>% 
  filter((name != 'pInt_seq_rev') & (name != 'pInt_seq_fwd') & (name != 'central_di_nt')) %>% 
  mutate(type = ifelse(name == 'bxb1_attP', 'mod_1', type)) %>% 
  mutate(type = ifelse(name == 'bla', 'mod_2', type)) %>% 
  mutate(type = ifelse((type != 'mod_1') & (type != 'mod_2'), 'mod_3', type)) %>% 
  mutate(name = ifelse(index %in% minor, '', name))

plot_plasmid(pInt_amp, name = 'pInt_attP5_ampR') + scale_fill_manual(values=c("#D36D55","#E8BF63", "#D3C0D6")) + theme(panel.background = element_rect(fill = 'white', color = 'white'))

ggsave('pInt_attP5_ampR.png',width = 6, height = 6)
  
```

```{r}
integration = 12:18
minor <- c(3,10,12,14,16, 18)

pInt_lcs <- read_gb(file = "pint_attp1_lcs_kanr.gb") %>% 
  as.data.frame() %>% 
  filter(name != 'Integration Payload') %>% 
  filter((name != 'pInt_seq_rev') & (name != 'pInt_seq_fwd') & (name != 'central_di_nt')) %>% 
  mutate(type = ifelse(name == 'bxb1_attP', 'mod_1', type)) %>% 
  mutate(type = ifelse(name == 'kanamycin resistance', 'mod_2', type)) %>% 
  mutate(type = ifelse((type != 'mod_1') & (type != 'mod_2'), 'mod_3', type)) %>% 
  mutate(type = ifelse(index %in% integration, 'mod_4', type)) %>% 
  mutate(name = ifelse(index %in% minor, '', name))

plot_plasmid(pInt_lcs, name = 'pInt_attP1_LCS_kanR') + scale_fill_manual(values=c("#D36D55","#E8BF63", "#D3C0D6",'#BED395')) + theme(panel.background = element_rect(fill = 'white', color = 'white'))

ggsave('pInt_attP1_LCS_kanR.png',width = 6, height = 6)
  
```

```{r}

minor <- c(3,11)

pInt_sacB <- read_gb(file = "pint_attp1_sacb_kanr.gb") %>% 
  as.data.frame() %>% 
  filter((name != 'pInt_seq_rev') & (name != 'pInt_seq_fwd') & (name != 'central_di_nt')) %>% 
  filter(name!='SacR') %>% 
  mutate(type = ifelse(name == 'bxb1_attP', 'mod_1', type)) %>% 
  mutate(type = ifelse(name == 'kanamycin resistance', 'mod_2', type)) %>% 
  mutate(type = ifelse((type != 'mod_1') & (type != 'mod_2'), 'mod_3', type)) %>% 
  mutate(type = ifelse(name == 'SacB', 'mod_4', type)) %>% 
  mutate(name = ifelse(index %in% minor, '', name))

plot_plasmid(pInt_sacB, name = 'pInt_attP1_sacB_kanR') + scale_fill_manual(values=c("#D36D55","#E8BF63", "#D3C0D6",'#BED395')) + theme(panel.background = element_rect(fill = 'white', color = 'white'))

ggsave('pInt_attP1_sacB_kanR.png',width = 6, height = 6)
  
```

```{r}

minor <- c(3,7,15)

payload <- 5:8

pInt_xis <- read_gb(file = "pint_attp1_tsxis_sacb_kanr.gb") %>% 
  as.data.frame() %>% 
  filter((name != 'pInt_seq_rev') & (name != 'pInt_seq_fwd') & (name != 'central_di_nt')) %>% 
  filter(name!='SacR') %>% 
  mutate(type = ifelse(name == 'bxb1_attP', 'mod_1', type)) %>% 
  mutate(type = ifelse(name == 'kanamycin resistance', 'mod_2', type)) %>% 
  mutate(type = ifelse((type != 'mod_1') & (type != 'mod_2'), 'mod_3', type)) %>% 
  mutate(type = ifelse(index %in% payload, 'mod_4', type)) %>% 
  mutate(name = ifelse(index %in% minor, '', name))

plot_plasmid(pInt_xis, name = 'pInt_attP1_tsXis_sacB_kanR') + scale_fill_manual(values=c("#D36D55","#E8BF63", "#D3C0D6",'#BED395')) + theme(panel.background = element_rect(fill = 'white', color = 'white'))


ggsave('pInt_attP1_tsXis_sacB_kanR.png',width = 6, height = 6)
  
```
````
