## Supplementary figures and images for "ORBIT for *E. coli*: Kilobase-scale oligonucleotide recombineering at high throughput and high efficiency"

### pHelper_Ec1_V1_ampR.png

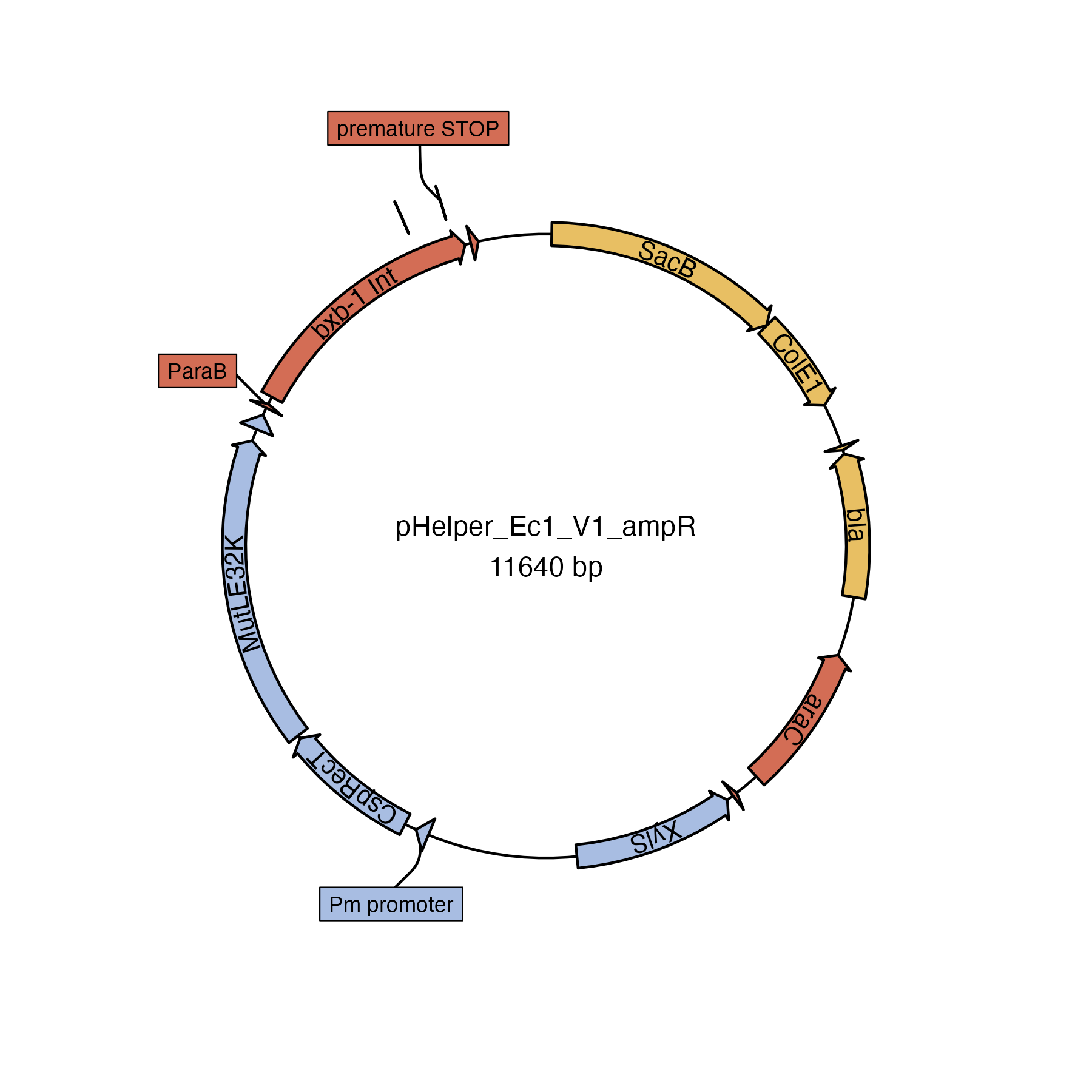

### pHelper_Ec1_V1_gentR.png

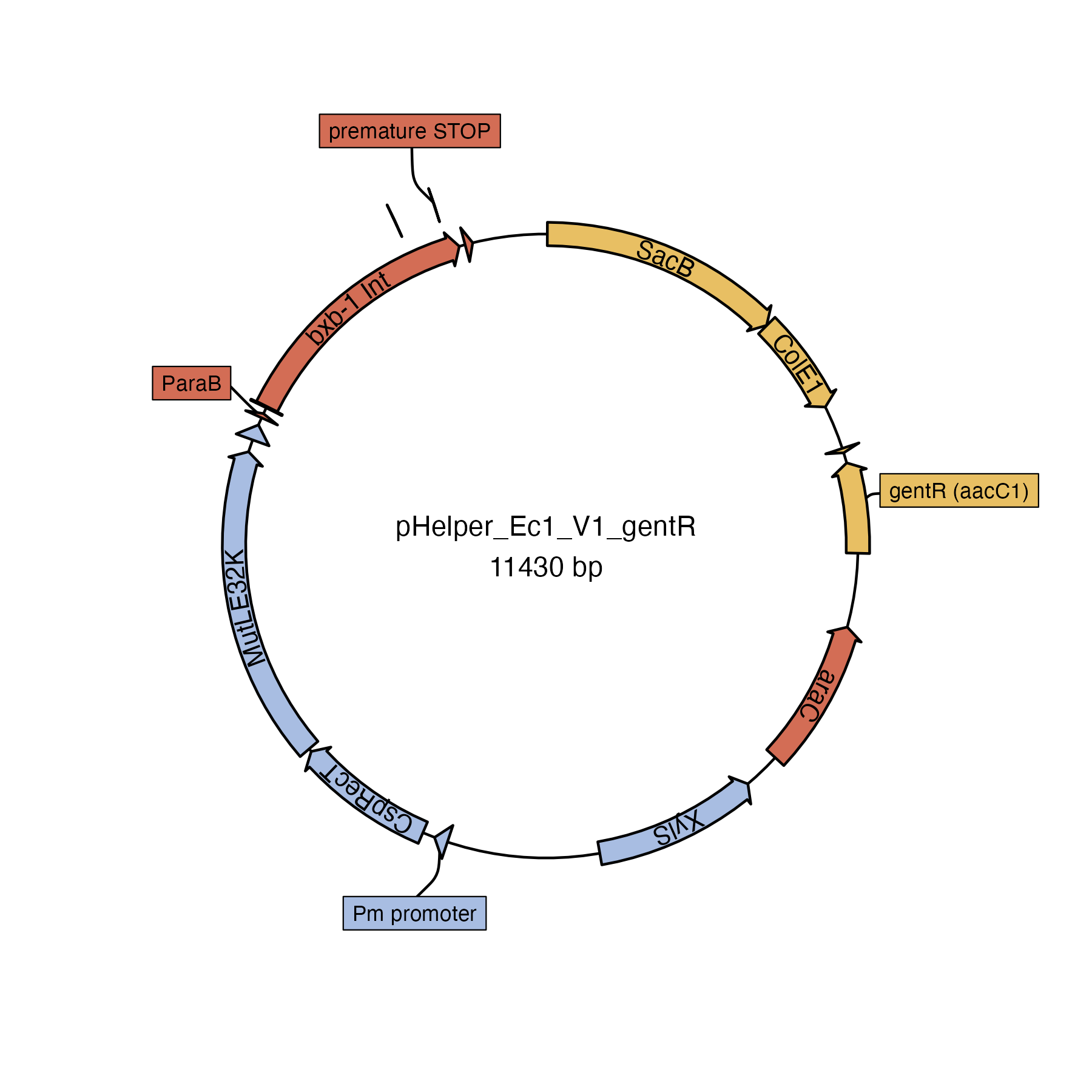

### pHelper_NoMutL_V2_gentR.png

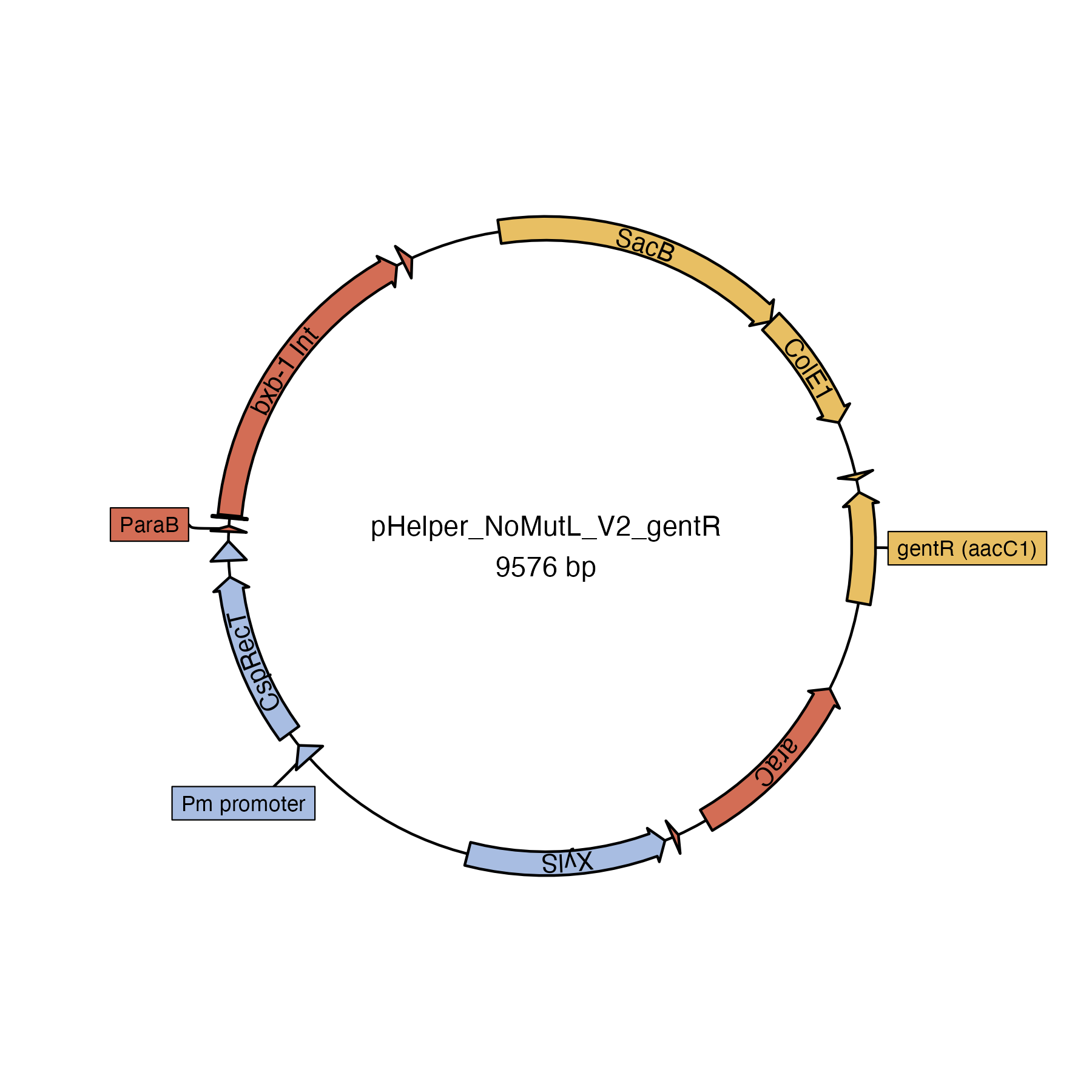

### pInt_attP1_ampR.png

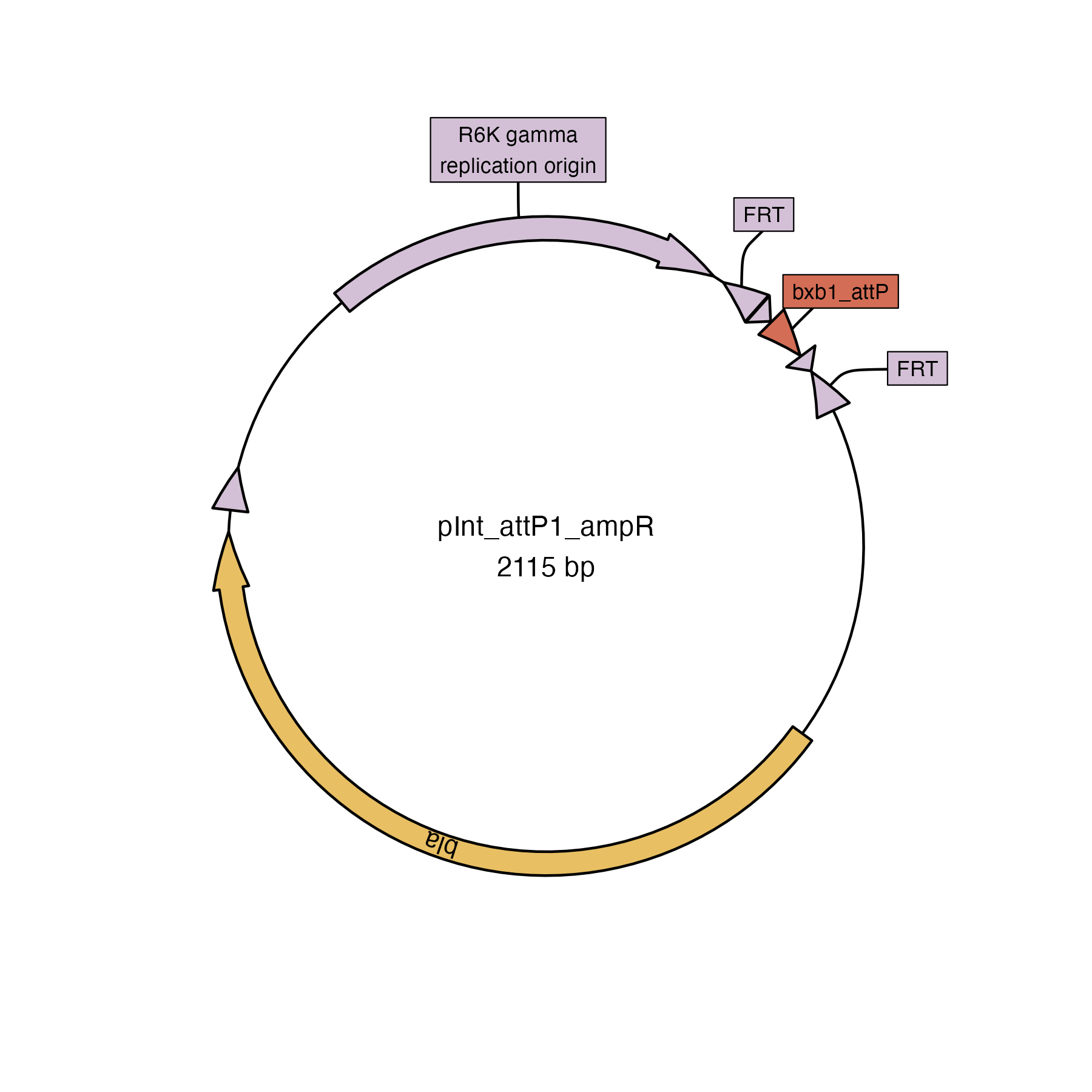

### pInt_attP1_chlorR.png

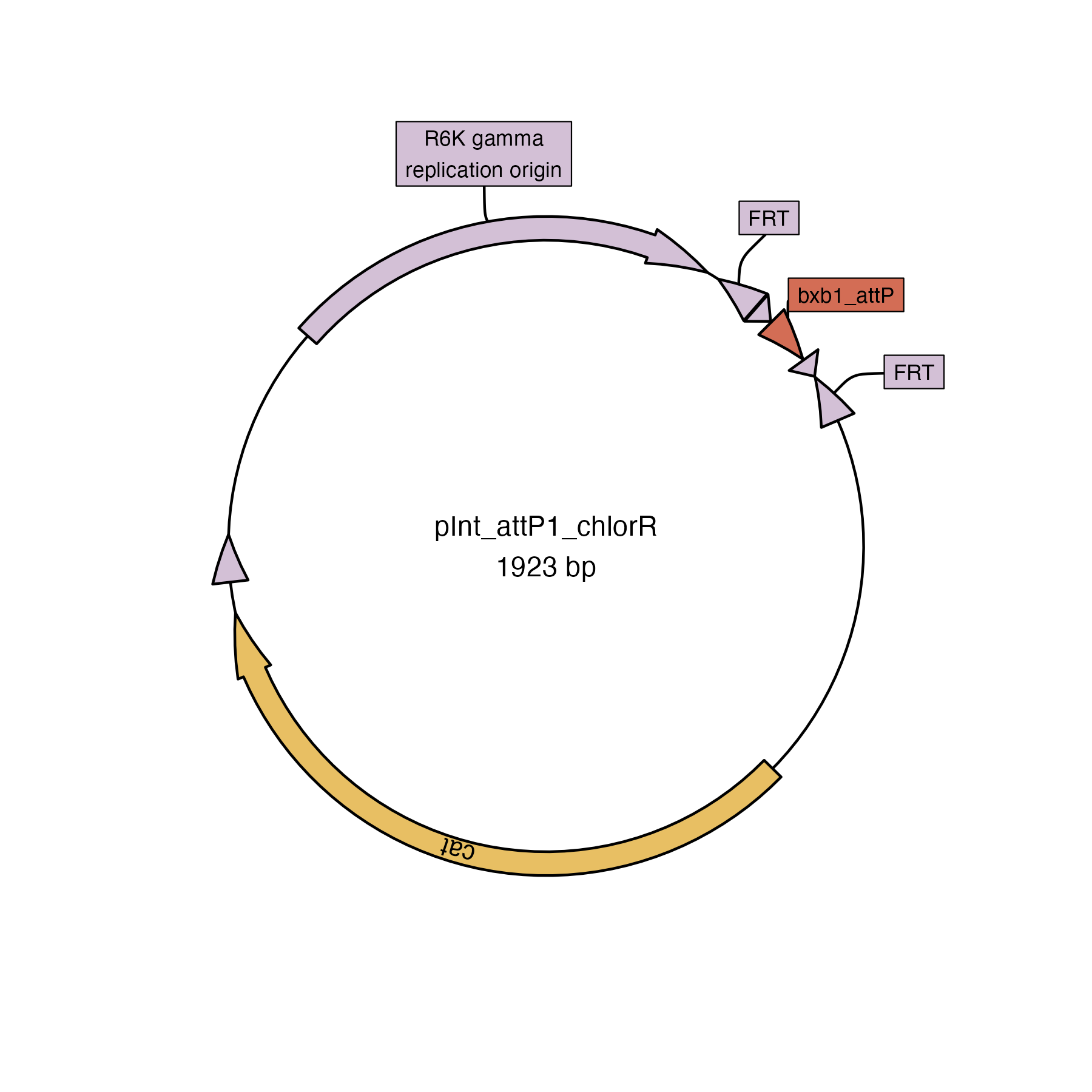

### pInt_attP1_kanR.png

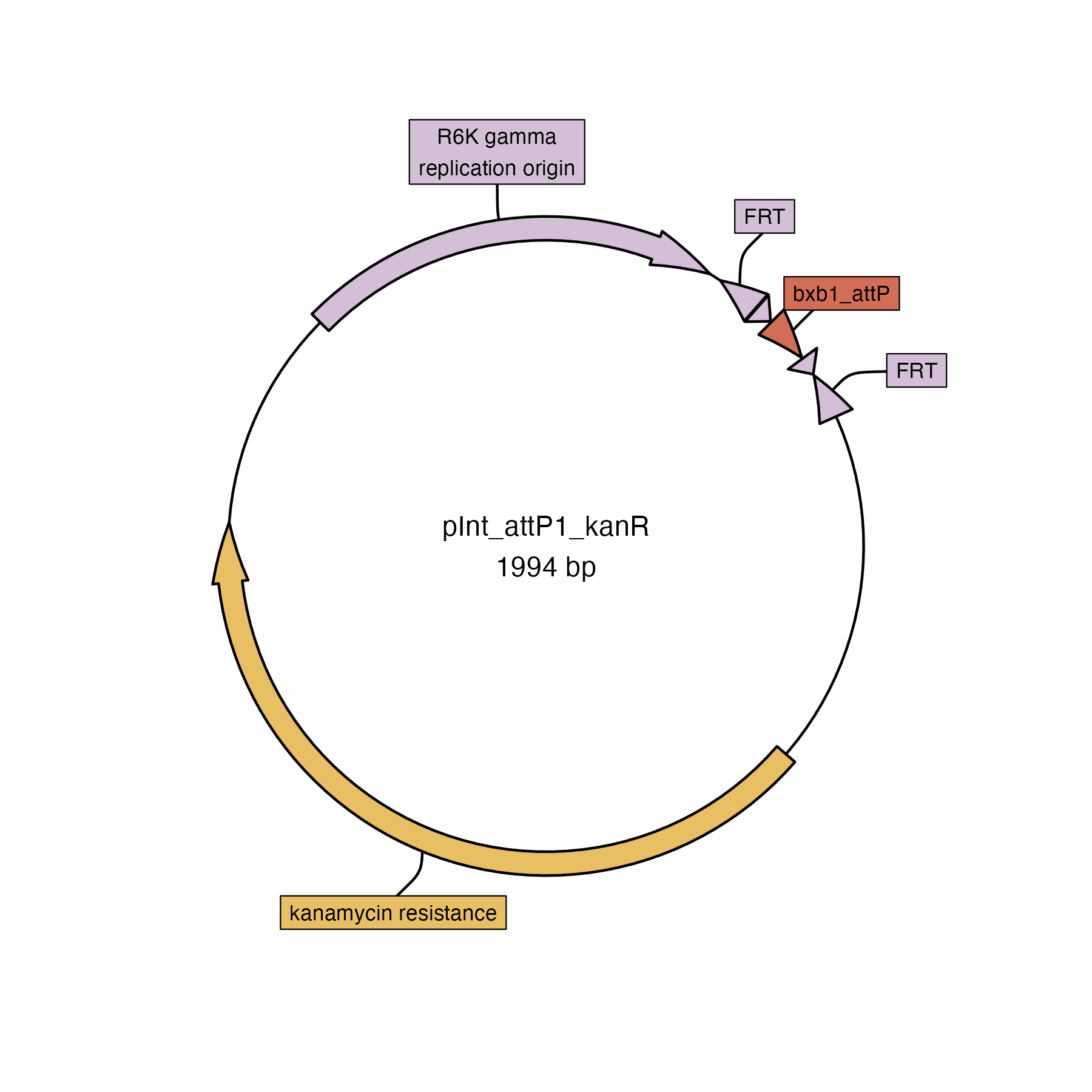

### pInt_attP1_LCS_kanR.png

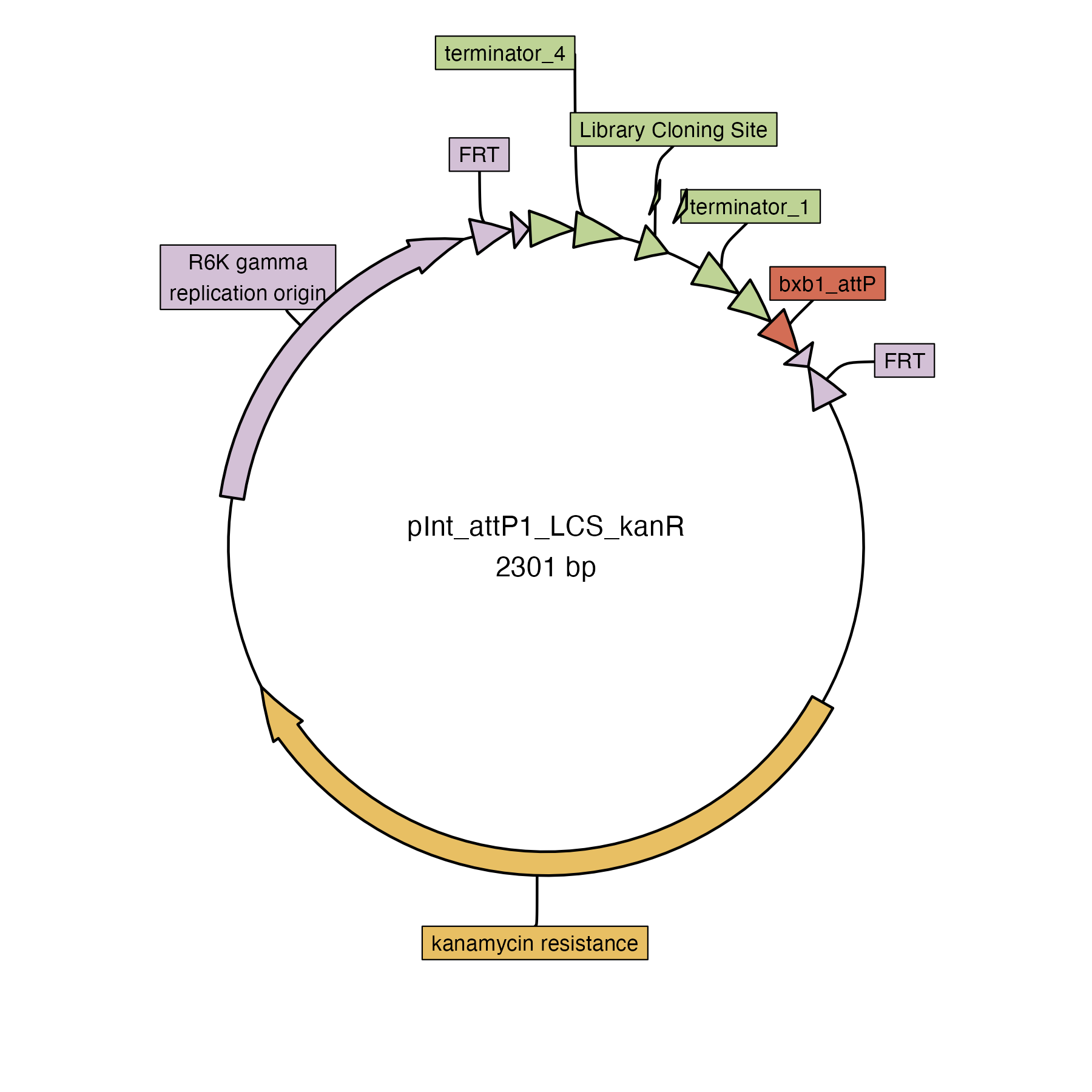

### pInt_attP1_sacB_kanR.png

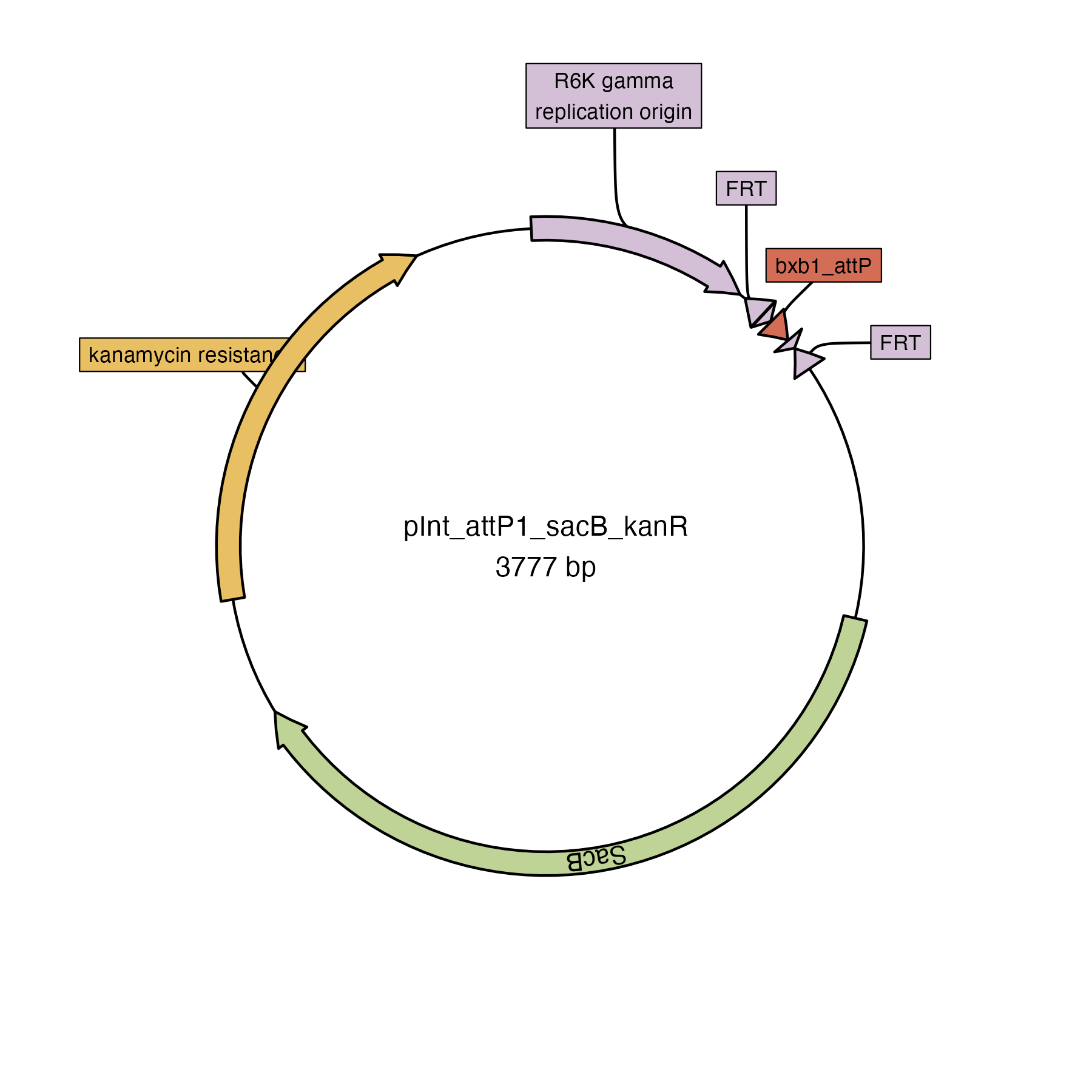

### pInt_attP1_tsXis_sacB_kanR.png

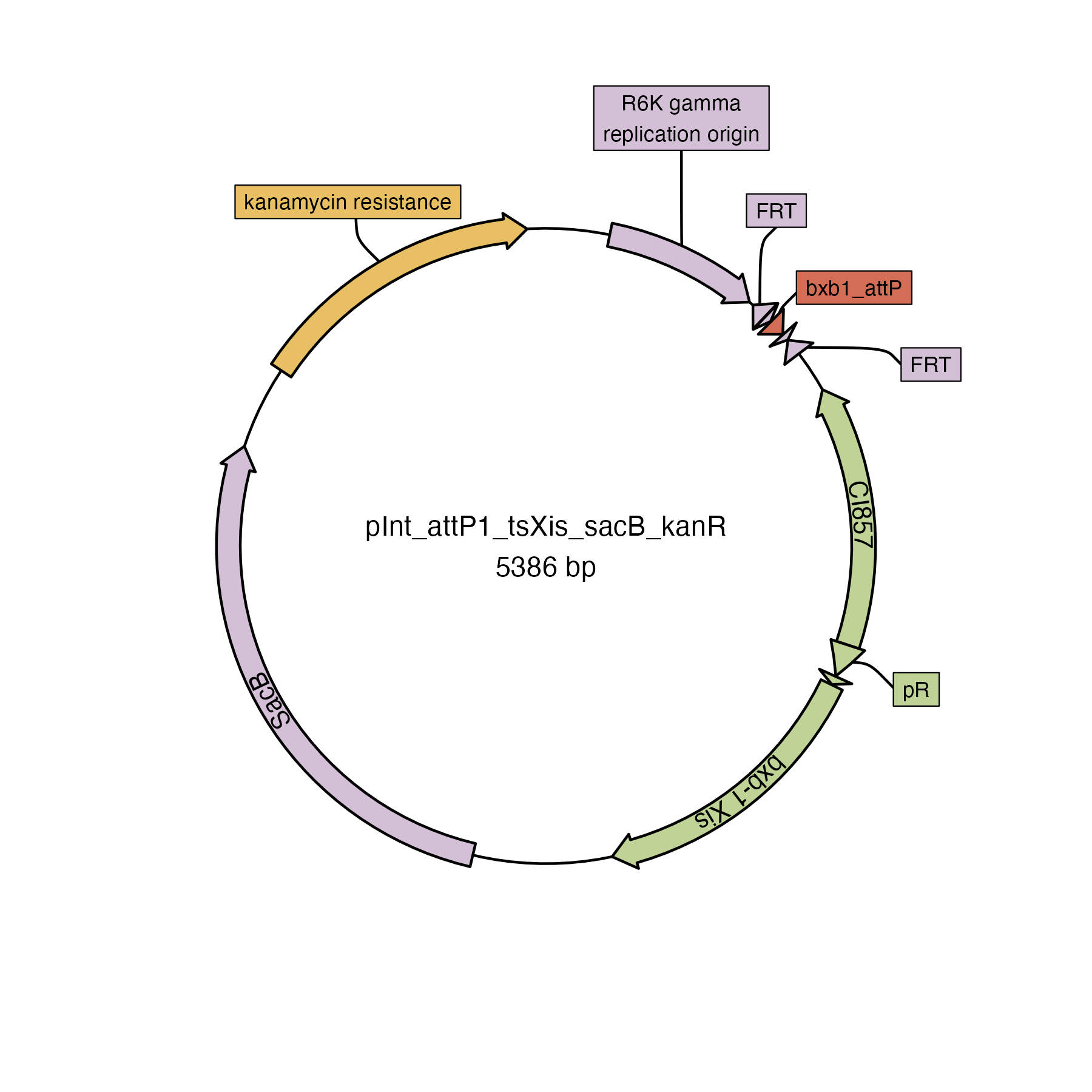

### pInt_attP2_kanR.png

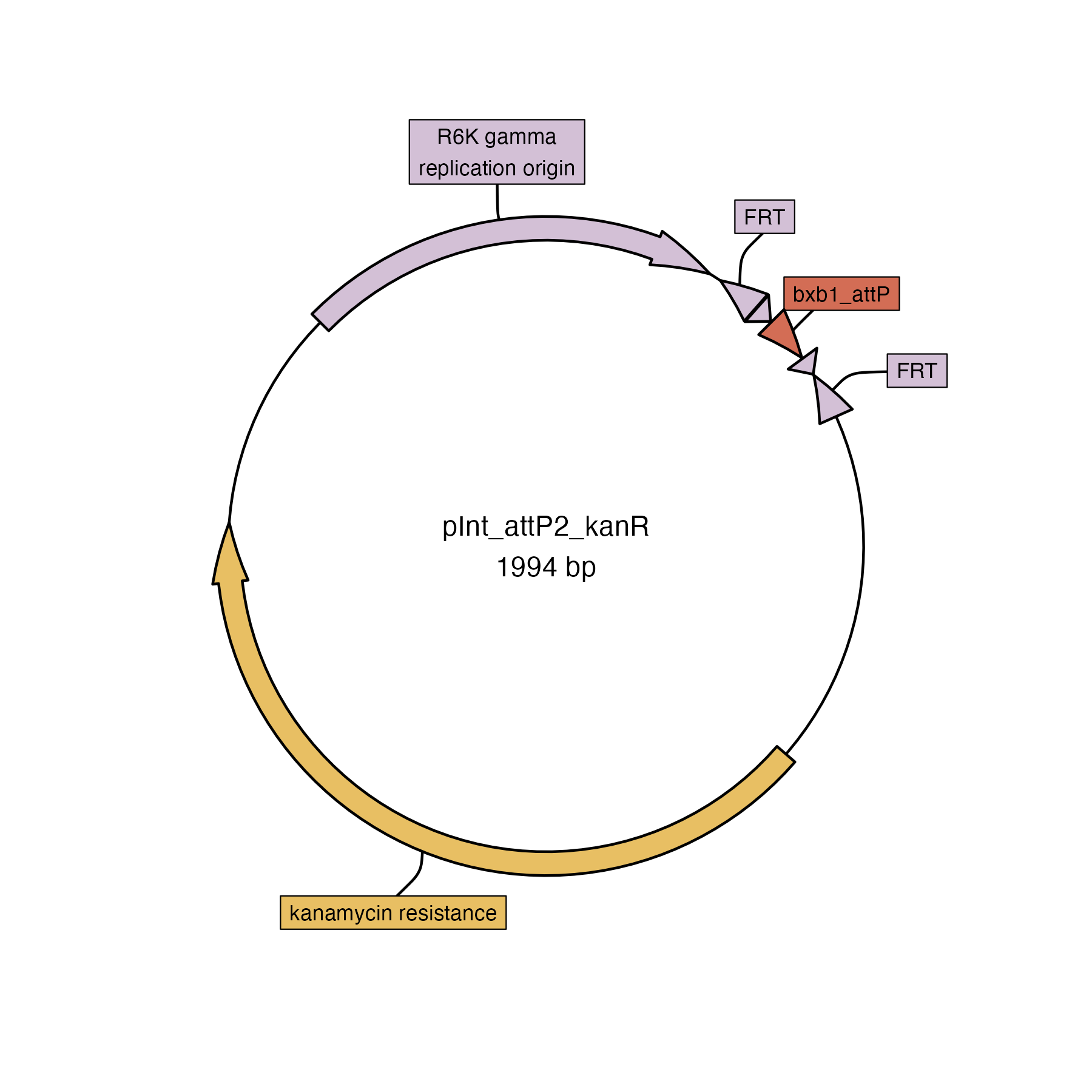

### pInt_attP3_chlorR.png

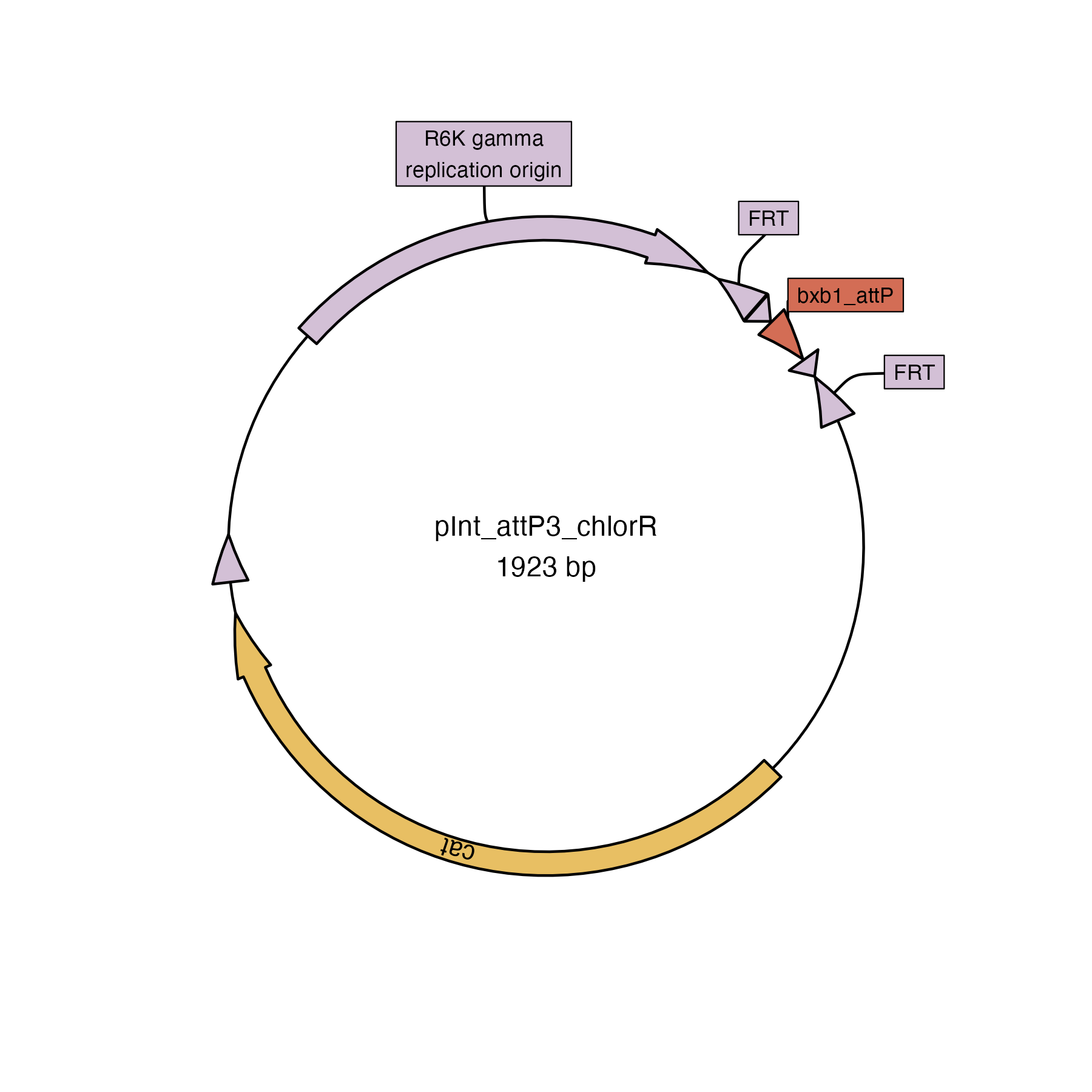

### pInt_attP5_ampR.png

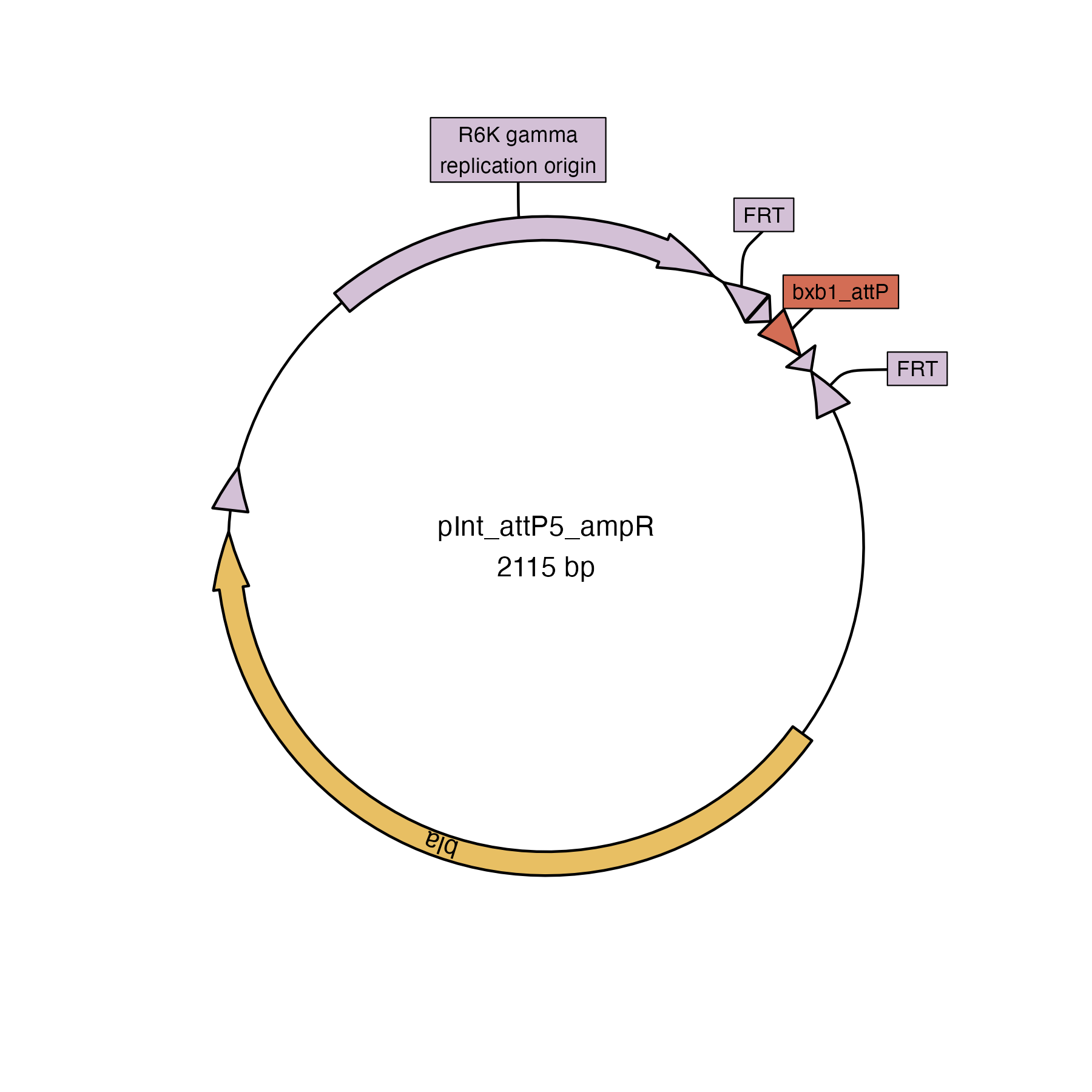
